# Unsupervised removal of systematic background noise from droplet-based single-cell experiments using CellBender

**DOI:** 10.1101/791699

**Authors:** Stephen J. Fleming, Mark D. Chaffin, Alessandro Arduini, Amer-Denis Akkad, Eric Banks, John C. Marioni, Anthony A. Philippakis, Patrick T. Ellinor, Mehrtash Babadi

## Abstract

Droplet-based single-cell assays, including scRNA-seq, snRNA-seq, and CITE-seq, produce a significant amount of background noise counts, the hallmark of which is non-zero counts in cell-free droplets and off-target gene expression in unexpected cell types. The presence of such systematic background noise is a potential source of batch effect and spurious differential gene expression. Here we develop a deep generative model for noise-contaminated data that is structured to reflect the phenomenology of background noise generation in droplet-based single-cell assays. The proposed model successfully distinguishes cell-containing from cell-free droplets without supervision, learns the profile of background noise, and retrieves a noise-free quantification in an end-to-end fashion. We present a scalable and robust implementation of our method as a module in the open-source software package `CellBender`. We show that `CellBender` operates close to the theoretically optimal denoising limit in simulated datasets, and present extensive evaluations using real datasets and experimental benchmarks drawn from different tissues, protocols, and modalities to show that `CellBender` significantly improves the agreement of droplet-based single-cell data with established gene expression patterns, and that the learned background noise profile provides evidence for degraded or uncaptured cell types.

## 1 Introduction

Droplet-based assays have enabled transcriptome-wide quantification of gene expression at the resolution of single cells [1, 2]. In a typical single-cell RNA sequencing (scRNA-seq) experiment, a suspension of cells is prepared and loaded into individual droplets. Poly(A)-tailed mRNAs in each droplet are uniquely barcoded and reverse-transcribed, followed by PCR amplification, library preparation, and ultimately sequencing. Quantifying gene expression in each cell is achieved by identifying and counting unique cDNA fragments that have a particular droplet barcode. The differential PCR amplification bias on different molecules can be reduced by using unique molecular identifier barcodes (UMIs), and counting the number of unique UMIs as a proxy for unique endogenous transcripts. This count information is then summarized in a count matrix, where counts of each gene are recorded for each cell barcode. The count matrix is the starting point for downstream analyses such as batch correction, clustering, and differential expression [3, 4]. In additional to cellular mRNA, other cell-endogenous molecules or incorporated perturbations (hereafter referred to as cell “features” for brevity) can be assayed using a similar setup by conjugating the desired feature with a cellular barcode. Examples include CITE-seq [5], Perturb-Seq [6], scCAT-seq [7], SNARE-seq [8], SHARE-seq [9], ECCITE-seq [10], 10x Multiome, among many other recently introduced droplet-based assays.

In order to reduce the rate of events where multiple cells are encapsulated in the same droplet, the cell suspension is appropriately diluted, and as a result, a typical droplet-based single-cell experiment produces hundreds of thousands of cell-free droplets. In an ideal scenario, a cell-free droplet is expected to be truly devoid of capturable molecules whereas a cell-containing droplet will yield features originating only from the encapsulated cell. In reality, however, neither expectation is met. On the one hand, the cell suspension contains a low to moderate concentration of cell-free mRNA molecules or other capturable features (Fig. 1a) which leads to non-zero molecule counts even in cell-free droplets [11] (Fig. 1b). These cell-free molecules, also referred to as “ambient” molecules, have their origin in either ruptured or degraded cells, residual cytoplasmic debris (e.g. in single-nuclei RNA-seq), or exogenous sources such as unbound ssDNA-conjugated antibodies or sample contamination. On the other hand, the shedding of capture oligos by beads in microfluidic channels as well as the formation of spurious chimeric molecules during the bulk mixed-template PCR amplification [12, 13] effectively lead to “swapping” of transcripts and barcodes across droplets. The severity of these problems depends on the tissue isolation protocol, as well as library preparation steps, including purification, size selection, PCR amplification conditioning and the number of cycles [14]. For a more thorough discussion, see Sec. S.2 in Supplementary Methods.

**Figure 1:**
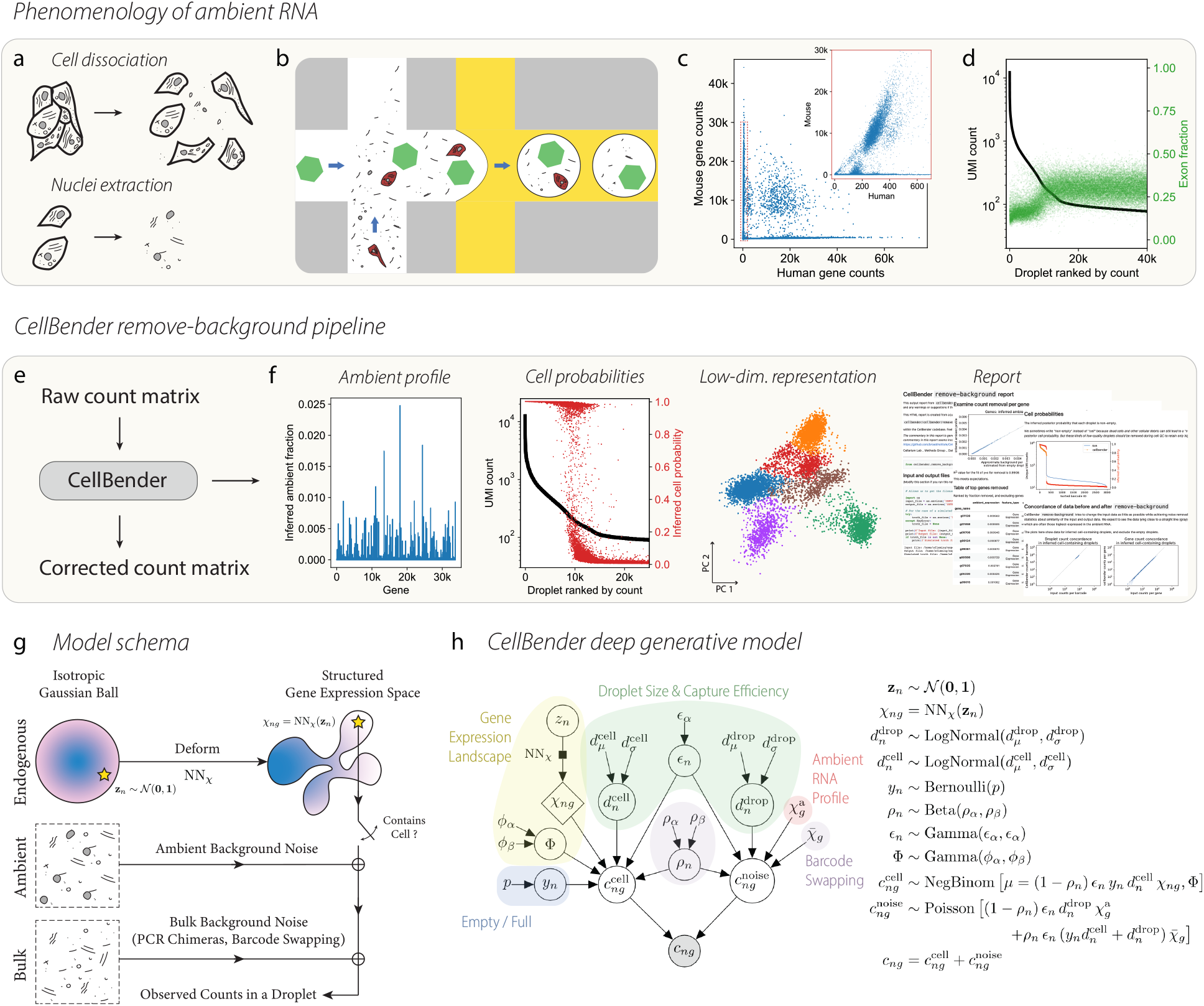
The phenomenology of ambient RNA and its deep generative modeling using `CellBender remove-background`. (a) Cell dissociation and nuclei extraction lead to the presence of cell-free RNA in solution. (b) Schematic diagram of the proposed source of ambient RNA background counts. Cell-free “ambient” RNAs (black lines) and other cellular debris are present in the cell-containing solution, and these RNAs are packaged up into the same droplet as a cell (red), or into an otherwise empty droplet that contains only a barcoded capture oligo bead (green hexagon). (c) Unique UMI counts per droplet that map to human and mouse genes for the publicly available `hgmm12k` dataset from 10x Genomics. The experiment is a mixture of human and mouse cells, and the inset (red box) shows that there are hundreds of human counts in droplets that contain mouse cells. (d) snRNA-seq Wistar rat heart dataset `rat6k`, showing unique UMI counts per droplet (black) with fraction of reads from exonic regions superimposed (green). The “ambient plateau” is the region of the rank-ordered plot with ranked barcode ID greater than about 15,000, where there are approximately 100 unique UMI counts per droplet. The increase in the fraction of exonic mapped reads coinciding with the onset of cell-free droplets shows that, in snRNA-seq, the ambient RNA is enriched for cytoplasmic material, where fewer intronic reads remain due to splicing. (e) Running `CellBender` is as simple sending a raw count matrix in and receiving a corrected count matrix in return. (f) Additional useful outputs include inferred latent variables of the model, such as the ambient RNA profile, probabilities that each droplet is non-empty, a low-dimensional embedding of gene expression per cell, and a summary report. (g) Schematic diagram explaining the rationale for our model. “True” cell counts are modeled using a flexible prior parameterized by a neural network NN_χ_. These counts (if a cell is present in a given droplet) are added to two constant noise sources: ambient background noise and bulk background noise. (h) The generative model for count data in the presence of background RNA.

Mixed-species experiments provide a direct demonstration of the effects of systematic background noise, as shown in Fig. 1c, where an experiment with a mixture of human and mouse cells is observed to have hundreds of off-target human transcripts in all droplets that contain mouse cells (inset), when ideally, mouse-cell-containing droplets would have zero human transcripts (excluding doublets, where two cells are captured in one droplet). The issue of background counts is particularly problematic in single-nuclei RNA sequencing (snRNA-seq). The harsh nuclear isolation protocols produce a significant number of ruptured nuclei and a high concentration of cytoplasmic RNA in the suspension (Fig. 1d, green dots). In severe cases, the typical total UMI count distinction between droplets with and without nuclei nearly disappears and all droplets lie on a continuum of counts. In such situations, successful downstream analysis hinges on our ability (1) to tell apart empty from non-empty droplets, and (2) to correctly recover the counts from encapsulated cells or nuclei while removing background counts.

The presence of background counts can reduce both the magnitude and the specificity of differential signal across different cell types. In cases where quantitative accuracy or specificity is required, e.g. for identification of exclusive marker genes as a part of drug target discovery, or the study of subtle phenotypic alterations in a case/control setting, background counts can obscure or even completely mask the signal of interest. In some experiments, extremely high expression of a particular gene in one cell type can give rise to a large amount of background, making it seem as though all cells express the gene at a low level. This issue is common to antibody features in CITE-seq and sgRNA CRISPR guides in Perturb-seq.

As the field of single-cell omics is rapidly extending beyond unimodal measurements and toward multimodality [15], the issue of systematic background noise remains a ubiquitous artifact that negatively impacts all such assays, regardless of the measured feature. A general-purpose *in silico* mitigation strategy is therefore expected to be of wide applicability. Here, we introduce a deep generative model for inferring cell-free and cell-containing droplets, learning the background noise profile, and retrieving uncontaminated counts from cell-containing droplets. Our proposed algorithm operates end-to-end starting from the raw counts, is fully unsupervised, is agnostic to the nature of the measured molecular feature (e.g. mRNA, protein, etc.), and requires no assumptions or prior biological knowledge of either cell types or cell-type-specific gene expression profiles. A major challenge in distinguishing background noise counts from biological counts for single droplets is the extreme sparsity of counts, such that without a strong informative prior, the counts obtained from a single droplet do not provide sufficient statistical power to allow inference of background contamination. Here, we utilize a neural network to learn the distribution of gene expression across all droplets. The learned distribution acts as a prior over cell-endogenous counts, provides a mechanism to share statistical power between similar cells, and ultimately improves the estimation of background noise counts. Learning this neural prior of cell states and estimating the background noise profile is performed simultaneously and self-consistently within a variational inference framework, allowing progressively improved separation of endogenous and background counts during model training.

We present extensive evaluation of our algorithm on both simulated and real datasets (wholecell, single-nuclei, mixed-species, and CITE-seq). We show that (1) our method is superior to the currently existing methods in distinguishing empty and cell-containing droplets, in particular, in ambiguous regimes and challenging single-nuclei RNA-seq datasets, and that (2) our method successfully learns and subtracts background noise counts from cell-containing droplets and leads to significantly increased amplitude and specificity of differential expression, both for RNA and CITE-seq antibody counts, and increases the correlation between the two modalities.

Our method is made available as a production-grade, easy-to-use command line tool (Fig. 1e,f). We utilize the Pyro probabilistic programming framework [16] for Bayesian inference. GPU acceleration is necessary for fast operation of this method. We refer to this method as `remove-background`, which constitutes the first computational module in `CellBender`, an open-source software package developed by the authors for pre-processing and quality controlling single-cell omics data. Several community-standard file formats, including CellRanger, DropSeq, AnnData [17], and Loom, are accepted as input. `CellBender` workflows are available on Terra (app.terra.bio), a secure open platform for collaborative omics analysis, and can be run on the cloud on a GPU with zero setup.

Since the time our method was first made available as an open-source project in 2019, it has been extensively used by the single-cell omics community in several large-scale studies, including primary research articles on the mouse brain [18], human brain organoids [19], human intestine [20], human heart [21–23], human and mouse adipocytes [24, 25], several recent studies on SARS-CoV-2 in human tissues [26–30], and a large snRNA-seq human cross-tissue atlas [31]. Background noise removal remains a crucial step in single-cell data analysis, and other authors have developed methods for remedying ambient RNA as well. In particular, DecontX by Yang *et al*. [32] is another principled method, which we benchmark together with our method here.

## 2 Methods

### 2.1 A generative model for noisy single-cell droplet-based count data

We build a probabilistic model of noise-contaminated single-cell data by examining the key steps of data generation process from first principles, including droplet formation and cell encapsulation, reverse transcription, PCR amplification, and the consequent ambient molecules and chimeric library fragments. These mechanisms, along with the empirical evidence for each, are discussed in detail in Sec. S.2 in Supplementary Methods. A simplified schematic of our model is shown in Fig. 1g, along with the formal probabilistic graphical model in Fig. 1h. Our general approach to modeling is discussed in Sec. S.1.1 in Online Methods. We review key elements of the probabilistic model in this section and refer the reader to Online Methods for supplementary details.

Our starting point is the observed feature count matrix *c_ng_*, where *n* and *g* denote cell index and feature index (e.g. gene), respectively. We interpret *c_ng_* as the sum of two non-negative contributions: the true biological counts originating from cells 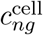, and the background noise counts 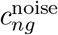. The background noise counts are drawn from a Poisson distribution:

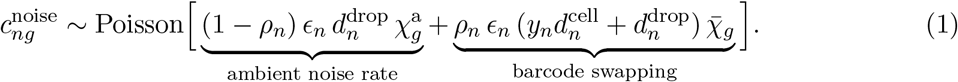

where the noise rate stems from two distinct processes: physically encapsulated ambient molecules, and barcode-swapped molecules, e.g. PCR chimeras. The ambient rate is determined by a learnable ambient profile 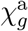, droplet size factor 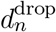, and droplet-specific capture efficiency factor *ϵ_n_*. We model barcode swapping as a diffusion process with a droplet-specific rate *ρ_n_* that is additionally modulated by the total amount of physically captured molecules in the droplet, i.e. 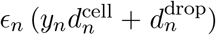, and the dataset-wide average gene expression (“pseudo-bulk”), 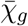. Here, *y_n_* ∈ {0, 1} is a binary variable that indicates cell presence in the droplet and 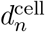 is the cell size factor.

The true biological counts 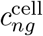 are modeled as a negative binomial distribution with a rate that depends on droplet-specific capture efficiency *ϵ_n_*, non-chimeric fraction 1 – *ρ_n_*, the cell presence indicator *y_n_*, cell size factor 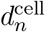, and a prior on true gene expression rate of the cell *χ_ng_*[**z**_*n*_]:

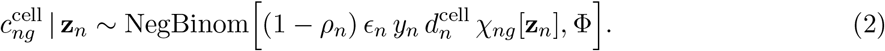

Here, Φ is a global learnable overdispersion parameter that modulates the uncertainty of the cell gene expression prior, and **z**_*n*_ is a droplet-specific latent variable that determines the gene expression rate prior *χ_ng_*. Crucially, the way we construct this prior is one of the components that makes the our model unique among noise removal approaches for count data. We utilize a neural network to learn a flexible prior for biological counts, which is realized as a deformation of a low-dimensional Gaussian latent space **z**_*n*_ (Fig. 1g). We fit the model using the stochastic variational inference (SVI) technique and leverage additional “encoding” neural networks for amortizing the approximate inference of droplet-specific (“local”) latent variables, see Supplementary Fig. S1b. Put together, our framework resembles a variational auto-encoder (VAE) [33] within a structured probabilistic model of noisy single-cell data.

We use the probabilistic programming language Pyro [16] to implement our model and the approximate variational inference algorithm. Our choice of variational posterior is shown graphically in Supplementary Fig. S1b and the details are provided in Sec. S.1.3 in Online Methods.

### 2.2 Constructing a denoised integer count matrix

`CellBender` generates several outputs following model fitting and inference, including the learned profile of ambient noise, cell containment probability per droplet, the low-dimensional latent space representation of cell states, and importantly, the estimated denoised integer count matrix, 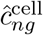. It is worth emphasizing that our sought-after denoised count matrix 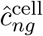 is not obtained by decoding the underlying low-dimensional latent embeddings of observed counts. This is a fundamental difference between our approach and VAE-based denoising and imputation methods [34–36]: encoding into and out of a low-dimensional latent space acts as an information bottleneck, smooths the data to varying degrees, and potentially masks subtle biological features such as transcriptional bursting, infrequent cell states, and other rare fluctuations of potential functional importance. In our approach, the low-dimensional latent space of cell states acts as a prior, which together with the observed data, determines the Bayesian posterior 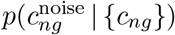. We estimate an integer matrix of likely-noise counts 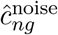 from the latter and obtain the denoised counts by subtracting off noise counts from observed counts.

Given the explicit partitioning of the observed data as a sum of non-negative signal and noise contributions, our approach explicitly guarantees the following: (1) each entry in the output count matrix will be less than or equal to the corresponding entry in the raw input matrix *c_ng_*; (2) the results are largely insensitive to the representational capacity of the encoding and decoding neural networks; (3) importantly, in a clean dataset where 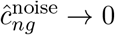, we obtain 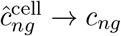, i.e. the data is not deformed, smoothed, or imputed. Our conservative approach to denoising is crucial for safe operation of our method in automated analysis pipelines, in particular, in application to clinical data and reference atlas building efforts.

Any noise removal algorithm involves a trade-off between removing actual noise (sensitivity) and retaining signal (specificity). In `CellBender`, we control this trade-off by means of a user-defined “nominal false positive rate” (nFPR) parameter (see Sec. S.1.5 in Online Methods). The nFPR parameter provides a transparent and interpretable handle to impose an upper bound on the amount of erroneously removed signal counts in aggregate (“false positive” counts), which could be either imposed separately on each feature, or globally. Larger nFPR values imply removing more noise at the expense of more signal. The ability to control denoising nFPR, regardless of the inherent noise of a given dataset, is desirable for integrative analysis of heterogeneous datasets such as clinical patient samples generated at multiple centers [26].

Finally, we note that reducing the posterior distribution of noise counts 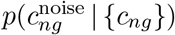, which is the natural output of a Bayesian model, to an integer point estimate 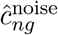, is a non-trivial and subtle task. The widely-used maximum *a posteriori* (MAP) estimator 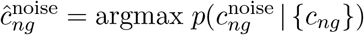, even though is a canonical Bayesian choice, leads to systematic under-estimation of noise counts for genes that are present in the ambient profile at low levels (see Sec. S.1.10 in Online Methods). Meeting the specified total noise target implied by nFPR while attaining the maximum modelbased posterior probability turns the estimation of 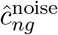 into a secondary optimization problem. We discuss and evaluate several such estimation algorithms in Sec. S.1.4 in Online Methods. By default (as of `CellBender` version v0.3.0), we use a constrained estimator that is formally equivalent to the multiple-choice knapsack problem (MCKP), which we show is exactly solvable using a fast and greedy coordinate ascent algorithm under mild assumptions, see Sec. S.1.9 in Online Methods.

## 3 Results

We present several evaluations of `CellBender` using real and simulated datasets in the following sections. Benchmarking on real scRNA-seq and snRNA-seq datasets shows that `CellBender` increases the specificity of known marker genes and significantly diminishes off-target gene expression. Compared to widely-used cell calling strategies, `CellBender` retrieves substantially more high-quality cells. Benchmarking on mixed-species scRNA-seq experiments demonstrates that `CellBender` removes the majority of off-target cross-species counts. Experiments using simulated noisy datasets with known ground truth show that `CellBender` operates close to the theoretically optimal limit. Finally, we show that `CellBender` significantly reduces background noise in CITE-seq data and increases the correlation between the cell-type specificity of mRNA and protein.

### 3.1 CellBender increases the specificity of marker genes and attenuates off-target expression

Removal of systematic noise from a dataset results in clearer biological insights by enhancing the specificity of gene expression and reducing spurious off-target counts. We demonstrate this by preprocessing a single-cell and a single-nuclei RNA-seq dataset with `CellBender` prior to downstream analysis, and assessing the biological soundness of the results.

We carried out a standard analysis workflow on the publicly available peripheral blood mononuclear cell (PBMC) scRNA-seq dataset (`pbmc8k`) from 10x Genomics dataset using scanpy [17]. We identified cell-containing droplets as having posterior cell probability *q_n_* > 0.5, and we used these cells in analyzing raw data and data pre-processed with `CellBender`. We further filtered cells using cutoffs for number of nonzero genes, percent mitochondrial counts, and an upper limit for UMI counts (details in Sec. S.1.12 in Online Methods). The results of the exact same analysis, with and without `CellBender` pre-processing, are shown in Fig. 2a-d, including the expression of several immune marker genes.

**Figure 2:**
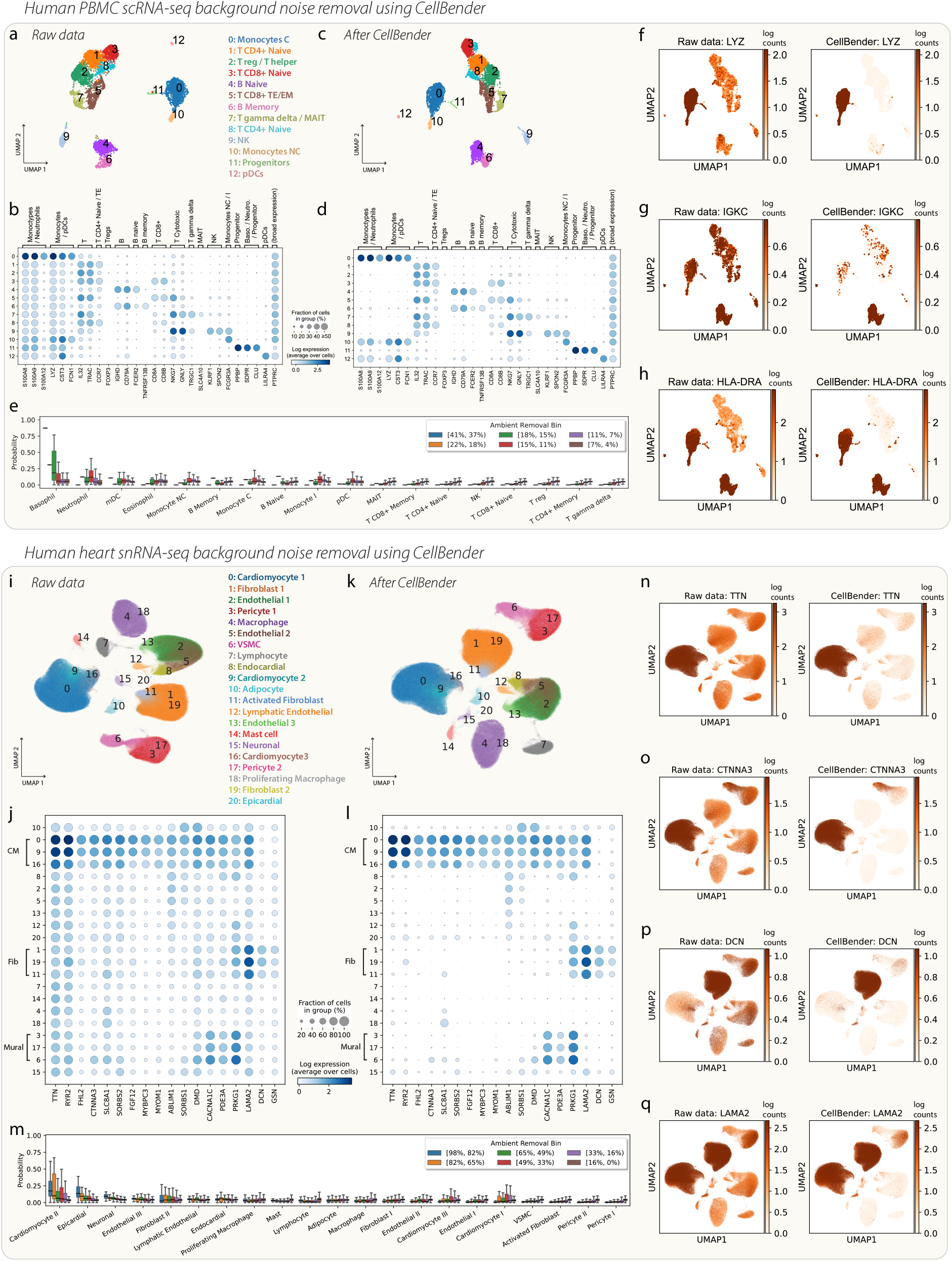
(a-h) Standard scanpy analysis of publicly-available 10x Genomics dataset `pbmc8k` with and without `CellBender`. UMAP visualizations of (a) the raw data and (c) the data pre-processed with `CellBender`. The dot plots display the expression of pre-defined marker genes for PBMCs for (b) the raw dataset and (d) the dataset processed with `CellBender`. (e) Removal of each gene has been mapped to cell type, indicating that cell types do not necessarily contribute equally to ambient RNA. (f-h) UMAP plots of the expression of *LYZ, IGKC*, and *HLA-DRA* in each cell before and after `CellBender`. Colorbar axes are truncated at the 80th percentile of per-cell expression. (i-q) Removal of background RNA from a published human heart snRNA-seq atlas, `heart600k`. (i) UMAP of raw data for nearly 600k nuclei. (j) Dotplot showing several highly-expressed genes in the raw dataset. (k-l) UMAP and dotplot after `CellBender`. (m) Similar to (e), this plot demonstrates that many of the removed counts are attributable to cardiomyocyte genes. (n-q) UMAP plots of the expression of *TTN, CTNNA3, DCN*, and *LAMA2* before and after `CellBender`. Colorbar axes are truncated at the 80th percentile of per-cell expression.

Raw gene expression data, as shown in Fig. 2b, indicates that the genes *S100A8, S100A9, LYZ, CST3*, and *PTPRC* are found to be abundantly and ubiquitously expressed in all clusters. While *PTPRC* (*CD45*) is a glycoprotein expressed on all nucleated hematopoietic cells, *LYZ* and *CST3* are known to be specific markers for monocytes and plasmacytoid dendritic cells (pDCs), whereas *S100A8* and *S100A9* are known to be specific markers of neutrophils, monocytes, and pDCs [37, 38], see Fig. S7. We hypothesized that the off-target expression of these genes was a result of systematic background noise. Fig. 2d shows the denoised counts obtained using `CellBender` at nFPR = 0.01 and demonstrates both sensitivity and specificity of `CellBender`: on the one hand, we observe that the expression of *S100A8, S100A9*, and *LYZ* is now largely concentrated on monocytes and pDCs, as expected. Conversely, we note that the biologically expected ubiquitous expression of *PTPRC* has remained unchanged. Supplementary Fig. S8 the expression of *LYZ* across clusters before and after `CellBender` in more detail. Supplementary Table S2 shows the differential expression for *S100A8, S100A9, LYZ*, and *CST3*, between monocytes C (cluster 0, where expression is expected) and naive B cells (cluster 4, where expression is not expected), calculated as the log2-fold-change using a Wilcoxon test. The log2-fold-changes (LFC) increase by a factor of two after subtracting background RNAl; in contrast, the LFC of *PTPRC* hardly changes at all. Another visualization of the effect of background noise removal is shown in Fig. 2f-h, where the expression of *LYZ* (increased specificity for monocytes), *IGKC* (increased specificity for B cells), and *HLA-DRA* (increased specificity for both monocytes and B cells) are plotted per-cell before and after `CellBender`. The increase in specificity is striking for these and many other examples, see e.g. Supplementary Fig. S9.

Next, we explored the origin of background counts in the PBMC dataset. Note that neutrophils and other granulocytes are absent from 10x PBMC cell clusters. The difficulty of capturing granulocytes is attributed to their sensitivity to rapid degradation after collection and poor isolation via density gradient centrifugation [39]. As such, we hypothesized that ambient counts might be enriched with granulocyte lysates. To test this hypothesis, we examined the fraction of counts removed by `CellBender` for each gene, and accordingly assigned each gene to the blood cell type with the highest consensus normalized TPM expression value obtained from the Human Protein Atlas (HPA) immune reference [38]. We binned the genes according to the fraction removed by `CellBender` as ambient noise, and interpreted the empirical frequency of assigning different cell types to the genes within each bin as the probability of contributing to the ambient soup. The result is shown in Fig. 2e and indicates that genes in the topmost ambient removal bins are associated with granulocytes at a significantly higher frequency. The top- and bottom-ten genes ranked by `CellBender` ambient removal are shown in Supplementary Fig. S6 along with the expression of each in the HPA immune reference, further demonstrating the enrichment of top-most ambient genes in basophils and neutrophils and the relative cell-type non-specificity of bottom-most genes.

PBMC scRNA-seq datasets are considered relatively clean in terms of ambient RNA contamination (see the UMI curve in Supplementary Fig. S4a-b as compared with Fig. S4e-f). Next, we examined a more challenging snRNA-seq dataset in which nuclei were extracted from frozen human heart tissue [23], `heart600k`. Nuclei preparations are more susceptible to ambient RNA contamination since the cells are all lysed and cytoplasmic mRNA becomes free in solution.

UMAP projections of the `heart600k` dataset were re-computed using Harmony-pytorch for batch effect correction [40], starting with either the raw counts (Fig. 2i) or the post-`CellBender` counts (Fig. 2k). The overall shape and appearance of the UMAP is qualitatively quite similar in both cases. However, an examination of gene expression shows that the dataset has been cleaned up quite significantly post `CellBender` (Fig. 2j,l). Fig. 2j demonstrates that, for many highly-expressed marker genes, the raw data would indicate that these genes are expressed in every cell type. However, it has been well established that the role of *TTN*, for example, is in the sarcomere of striated muscle cells including cardiomyocytes, and it is not expressed in the other cell types present in this experiment. Fig. 2l,n demonstrate that, after `CellBender`, the expression of *TTN* becomes much more specific to the cardiomyocyte clusters. Similarly, *CTNNA3*, involved in cell-cell adhesion in muscle, appears much more specific to cardiomyocytes and vascular smooth muscle cells (cluster 6: VSMC) after `CellBender` (Fig. 2l,o), in agreement with existing heart snRNA-seq atlases [21, 41]. The expression of *DCN*, which plays a role in collagen fibril assembly in the extracellular matrix, becomes much more specific to fibroblasts (Fig. 2l,p), also consistent with Refs. [21, 41]. Finally, the expression of *LAMA2*, another component of the extracellular matrix, is found after `CellBender` to be much more specific to fibroblasts and cardiomyocytes, with some lower-level expression in pericytes, adipocytes, and neuronal cells, again in agreement with Refs. [21, 41].

Cardiomyocytes have high UMI counts as compared to other cell types (see, for example, Supplementary Fig. 2b from Ref. [21], where the cardiomyocytes can have an order of magnitude higher UMI counts than other cell types in snRNA-seq). We hypothesized that we should see a disproportionately high amount cardiomyocyte genes in the background RNA removed by `CellBender`. An examination of genes preferentially removed by `CellBender` shows that the top genes in terms of removed fraction are in fact associated mainly with cardiomyocytes, and to a lesser extent with epicardial cells; see Fig. 2m. Many of the genes plotted in Fig. 2j,l are cardiomyocyte marker genes, including some of the most highly-expressed genes in the dataset, *TTN* and *RYR2*. This highlights the importance of learning the ambient RNA profile from the dataset itself: the large amount of ambient cardiomyocyte mRNA, which is packaged into each droplet as background counts, is appropriately targeted and removed by `CellBender`, vastly improving the specificity of gene expression for downstream biological analyses.

### 3.2 CellBender accurately identifies cell-containing droplets

As a part of model training and inference, `CellBender` produces a posterior probability *q_n_* that droplet *n* contains a cell. While this determination can be rather trivial in some pristine datasets (e.g. the PBMC dataset `pbmc8k`, see Supplementary Fig. S4a-b), complicated experimental factors and excessive amounts of ambient RNA contamination often make this determination rather challenging (see, e.g., the snRNA-seq dataset `rat6k` in Supplementary Fig. S4e-f). A variety of heuristics are typically employed in order determine cutoffs for thresholding cells versus empty droplets, as in CellRanger v2. More principled approaches have been developed, including CellRanger v3+ and `EmptyDrops`. CellRanger v3+ and `EmptyDrops` [42] use statistical tests to ascertain which droplets have expression profiles significantly different from empty droplets. In our algorithm, the determination of empty vs. non-empty droplet is a result of disentangling background counts from endogenous feature counts during model training, where both the gene expression and total UMI counts of all droplets are taken into account.

Fig. 1f panel 2 shows the posterior cell probabilities for the first 25,000 droplets of the `rat6k` rat heart snRNA-seq dataset. Note that the algorithm in general identifies cells and empty droplets as expected, and that the transition between the two is not based on a hard UMI cutoff. A determination of cell-free vs. cell-containing can be obtained by thresholding based on the posterior probability, *q_n_*. The algorithm converges to largely binary probability values for the majority of droplets and the precise choice of threshold value affects relatively very few droplets in practice.

We compare the cell calls made by `CellBender` with three other methods in common use (CellRanger v2, CellRanger v3, and `EmptyDrops`) in Fig. 3. Fig. 3a shows that `CellBender` generally calls more cells than CellRanger (Sec. S.6.6 in Supplementary Methods), many of which lie farther down the UMI curve (black) and are not called by other methods.

**Figure 3:**
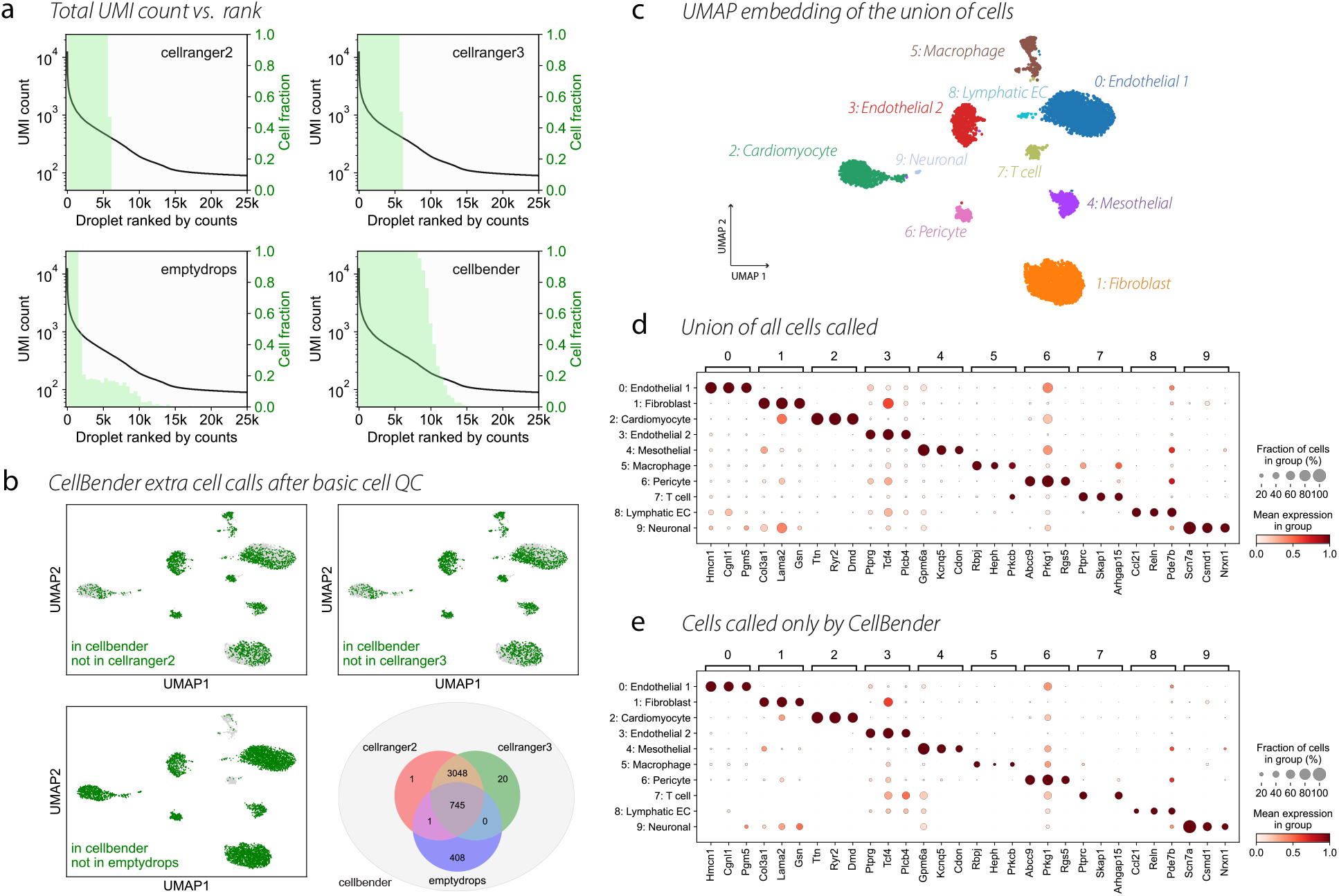
Comparing four cell calling algorithms (CellRanger v2, v3, EmptyDrops, and `CellBender`) on the `rat6k` snRNA-seq dataset. (a) Detected cells for different algorithms: the UMI vs. barcode rank curve (black line) is superimposed on the fraction of detected cell-containing droplets in different barcode rank bins (green bars). CellRanger results indicate imposing a nearly hard cutoff on the barcode rank, while EmptyDrops calls several cells between 6000 and 10,000 in UMI count rank (x-axis). (b) `CellBender` detects all cells called by the other algorithms (after cell QC) and many more. UMAP embeddings were generated after performing cell QC. All cells are shown in gray, with green dots superimposed to denote cells which were not detected by the method in question but detected by `CellBender`. The Venn diagram quantifies the agreement between various methods. (c) UMAP with cell type labels at Leiden resolution 0.5. All clusters appear to be biologically meaningful. (d) Top three marker genes for each cluster (scanpy Wilcoxon test) are shown for the union of all cells called by any algorithm (which coincides with `CellBender` cell calls). (e) Same marker gene dotplot as in (d), but now showing only those cells which were exclusively detected by `CellBender`. The similarity to (d) and presence of real marker genes indicates that the extra cell calls made by `CellBender` are real.

The set of cells called by `CellBender` contains all the cells called by CellRanger v2, v3, and `EmptyDrops` after cell quality control (see Venn diagram in panel Fig. 3b). In addition, `CellBender` detects more than 50% extra cells compared to CellRanger v3, and more than five times as many cells as `EmptyDrops`. Given the significant ambient RNA contamination in this dataset, we naturally hypothesized that many of the extra cell calls made by `CellBender` might have been cytoplasmic debris which were nevertheless statistically different from the ambient RNA in terms of gene expression makeup. To evaluate this hypothesis, we obtained a UMAP embedding of cells detected only by `CellBender` together with the cells detected by other methods (Fig. 3b) after typical filtering by gene complexity and mitochondrial fraction (see Sec. S.1.12 in Online Methods). To our surprise, (1) over 40% of the extra cells called by `CellBender` pass quality control filters, amounting to over 1600 cells (Supplementary Table S3), and (2) the extra cell calls made by `CellBender` clustered together with cells called by the other algorithms. Fig. 3c shows the UMAP embedding obtained from the union of all cells called by any algorithm (after cell QC filtering) with putative cell type labels, and it can be seen that the cells called exclusively by `CellBender` have a marker gene distribution (Fig. 3e) similar to the dotplot created using the union of all cells called by any algorithm (Fig. 3d). `EmptyDrops` calls many low-UMI-count cells that CellRanger v2 and v3 miss, though it also misses a large number of relatively high-UMI-count droplets along the rank-ordered UMI plot. This is likely due to the similarity between gene expression of the empty drops and the most populous cell types in this particular experiment (Sec. S.6.6 in Supplementary Methods). As such, the Dirichlet-multinomial likelihood model employed in `EmptyDrops` does not yield a statistically significant probability of being non-empty for cardiomyocyte-containing droplets. In contrast, `CellBender` learns the expression profile of cardiomyoctyes from high-count droplets and is not impacted.

Finally, we recommend performing additional biologically-motivated and tissue-specific quality control on `CellBender` cell calls whenever possible, e.g. using mitochondrial read fraction, exonic read fraction, and gene complexity, as suggested by previous authors [42, 43]. We have deliberately avoided including such filters in `CellBender` to allow broad applicability of this method. Post-`CellBender` quality controlling strategies must be informed by the studied biological system and the protocol. To emphasize the importance of post-filtering, we show a plot of the fraction of reads per droplet that come from mitochondrial genes in the `hgmm12k` dataset in Supplementary Fig. S12. It can be clearly seen that many low-UMI droplets exhibit a high fraction of mitochondrial genes (possibly dead or dying cells), and because they are distinct from empty droplets, they are nevertheless assigned a high probability of containing cells by `CellBender`. After filtering the detected cells based on mitochondrial read fraction, some of these lowest-count and degraded cells will be naturally filtered out. The analysis shown in Fig. 3 includes such post-filtering criteria.

### 3.3 CellBender significantly reduces off-target gene expression in mixed-species experiments

A definitive and straightforward experimental benchmark to evaluate the level of background noise and the efficacy of mitigation strategies is a mixed-species experiment, where two cell types from different species are combined and assayed together. This would ideally result in droplets containing exclusively feature counts from one species or the other, but due to the presence of background noise, this is not the case (as shown in Fig. 1c). Here, we use the publicly available human-mouse mixture dataset from 10x Genomics (`hgmm12k`) to evaluate `CellBender` and also compare to DecontX [32], another method for removing background noise.

Fig. 4a shows a scatter plot of human and mouse gene expression in each droplet (doublet droplets not shown) in raw data and for `CellBender` processed data at different nFPR settings on a logarithmic scale (data plotted on linear axes in Supplementary Fig. S13). The raw data shows hundreds of off-target cross-species counts in each droplet (best visible in the side histograms). After removing background noise, we would ideally expect all cross-species counts to be removed. Indeed, `CellBender` (with default nFPR of 0.01) reduces off-target counts to a median of 19 per cell, i.e. by over an order of magnitude from the raw data, with a median of 225. At a nFPR setting of 0.1, the median off-target counts per cell drops to 4 (Supplementary Table S5 and Fig. S14). It is worth re-emphasizing that `CellBender` is a completely unsupervised model, and that the algorithm achieves this level of denoising without the knowledge of human genes, mouse genes, or that this is a mixture-species experiment.

**Figure 4:**
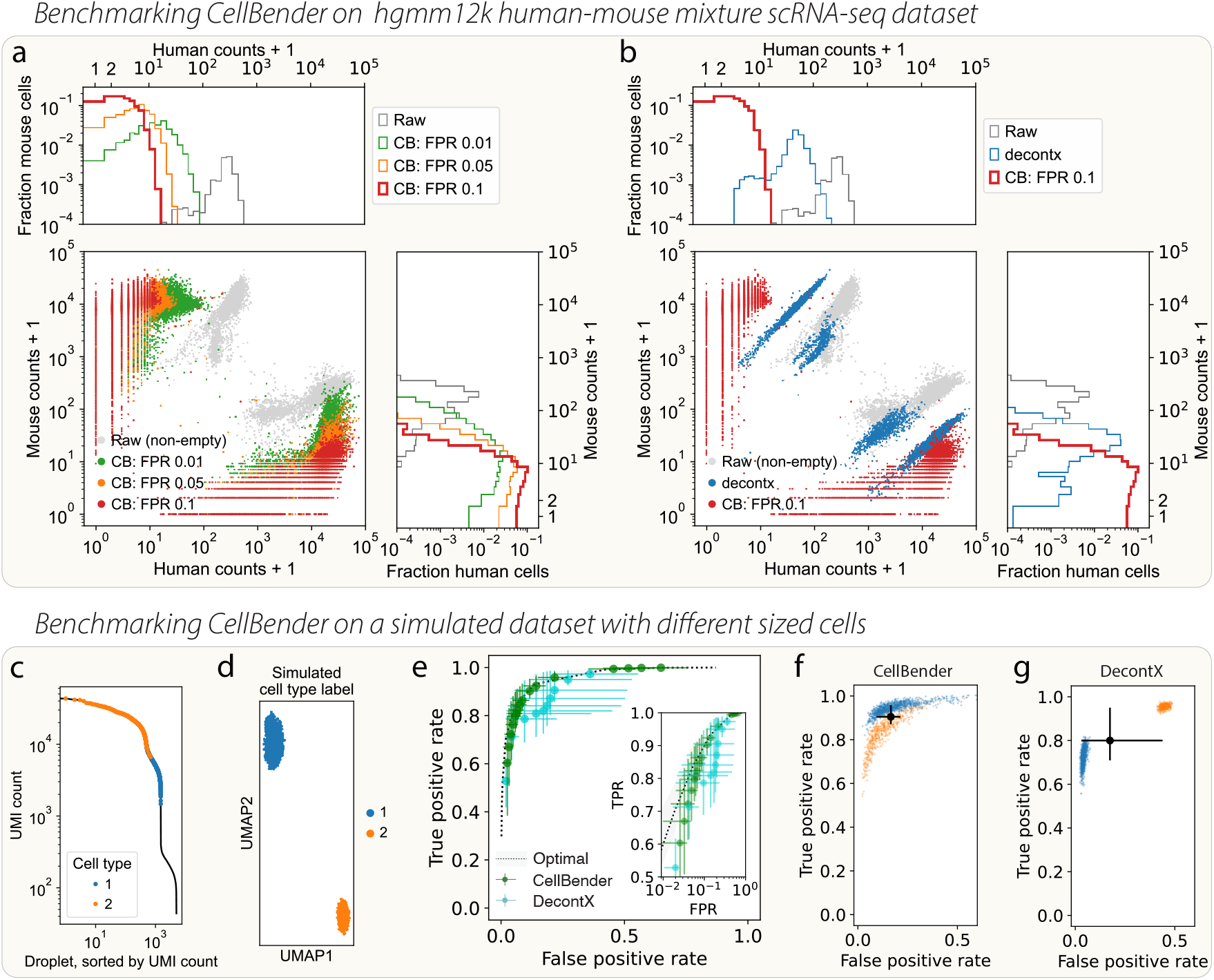
Benchmarking `CellBender` on denoising the `hgmm12k` human-mouse mixture dataset (a-b), and a simulated dataset with different sized cells (c-g). (a) Log-scale plot of species mixing shows that raw data (gray) contains several hundred counts of mouse transcripts in human cells and vice versa. CB removes most of the off-target noise. The marginal histograms show that many human cells end up with zero mouse counts and vice versa. CB denoised counts are shown for several nominal false positive rates (nFPR) choices. (b) Same plot as in (a) but with DecontX included for comparison. (c) The UMI curve for the simulated dataset, showing cells and empty droplets. Simulated cell type 2 has many more UMI counts than cell type 1. (d) The UMAP created from cells called by CB. (e) ROC curve quantifying per-cell noise removal performance. Black dashed line with gray shading (one standard deviation in per-cell performance) represents the best possible performance given perfect knowledge of all latent variables in the simulation and is only limited by sampling noise. Large green dots (mean) with error bars (first and third quartiles in per-cell performance) show CB outputs at a variety of expected nFPR values. Cyan dots with error bars show DecontX output using different values of the parameter `delta`. (f-g) Comparison of per-cell performance of DecontX (default settings) and CB (matching the output FPR of default DecontX), where cells are colored by cell type. DecontX treats the different cell types rather differently in terms of FPR (blue and orange colors are cell types from c-d). `CellBender` is abbreviated as CB on the plots.

Fig. 4b compares the performance of `CellBender` with DecontX [32]. We find that while DecontX removes a large number of cross-species counts, `CellBender` has a significantly higher sensitivity: in fact, at nFPR 0.1 (in red), `CellBender` removes all cross-species counts from 16% of cells (see the marginal histograms in Fig. 4a, where “1” means there are zero cross-species counts). In addition, the results obtained using `CellBender` show other important characteristics which are worth emphasizing:

- The amount of background noise which gets removed can be tuned using the interpretable expected nFPR parameter, as shown in Fig. 4a,e.
- `CellBender` largely removes the linear trend in the relationship between cross-species counts and cell endogenous counts (the linear trend seen in raw data shown in gray, see also Supplementary Fig. S13). The proportional relation between background noise counts and cell endogenous counts has been associated with library PCR chimeras formed during mixed-template amplification [13] which effectively leads to random barcode swapping between library fragments. Another potential mechanism is droplet-to-droplet variability in capture efficiency which also leads to a proportional relation between endogenous and noise counts. Both of these phenomena are modeled in `CellBender`, see Sec. S.1.2 in Online Methods. Note that this linear trend remains largely unmitigated by DecontX, see Fig. 4b and Supplementary Fig. S13b.
- We find that DecontX treats different groups of cells from the same species differently, which can be seen as the fragmentation of blue points in Fig. 4b. We hypothesize this non-uniform performance to be associated with the hard clustering preprocessing step in DecontX. While the user can provide their own clustering to DecontX to mitigate this issue, `CellBender` sidesteps such issues altogether by avoiding hard clustering entirely, and instead allows similar cells to share statistical power via a low-dimensional continuous latent space.

### 3.4 CellBender operates near the theoretically optimal limit on simulated datasets

So far, we have shown evaluations of `CellBender` using real datasets and resorted to prior biological knowledge (e.g. marker genes) or expected outcomes (as in mixed-species experiments) to assess the soundness of the results. Here, we additionally show experiments using simulated data, with known noise and signal contributions, to evaluate the performance of `CellBender` theoretically and in a more controlled setting. Fig. 4c-g shows the results of inference using a simulated dataset with 10,000 genes, generated according to a noise model that includes both ambient sources and barcode swapping (see Sec. S.1.14 in Online Methods for simulation details). Importantly, the `CellBender` model is slightly mis-specified for this simulated data on purpose, as the simulation draws “true” gene expression *χ_ng_* from a Dirichlet distribution with fixed concentration parameter per cell type. Panels (c-g) show a simulation with two “cell types” with unique underlying expression profiles, where the cell types have a very different number of UMI counts. The ambient profile in the simulation is a weighted average of total expression.

Fig. 4e shows the noise removal performance as a receiver operating characteristic (ROC) curve. Noise counts that are correctly removed are counted as “true positives”, and a “false positive” is a cell-endogenous count that is erroneously removed. A hypothetical model with perfect knowledge of every real and noise count would be represented by the point (0, 1) in the FPR-TPR plane. The stochasticity of the data generating process and finite sequencing depth, however, make this perfect limit theoretically out of reach, even with prefect knowledge of all latent variables.

We show the “best theoretically achievable performance”, given perfect knowledge of all latent variables, as the black dashed line. `CellBender` comes quite close to this optimal performance (green dots, obtained by running at increasing nFPR parameters). Supplementary Table S6 shows a decent agreement between the specified nFPR and empirical FPR. The DecontX ROC curve was created by running the tool with several values of hyperparameter `delta`. Default DecontX parameters were found to correspond to an empirical FPR of 0.142 and TPR of 0.809. Run with nFPR=0.0442, `CellBender` was found to have exactly the same TPR of 0.809, but the FPR was 0.062. This means that, for the same amount of removal of noise, DecontX removed more than twice as much signal as `CellBender`. At nFPR=0.125, `CellBender` matched the DecontX FPR of 0.142, but the TPR was 0.923. This means that, for the same value of removal of real signal, `CellBender` was able to remove 92.3% of the noise, while DecontX removed 80.9%. This seems to be due to DecontX treating the two simulated cell types differently in terms of where they land on the ROC curve (see Fig. 4f,g).

### 3.5 CellBender removes ambient antibody counts from CITE-seq data and increases correlation between protein and RNA levels

As mentioned in the introduction, `CellBender` makes no assumption about the nature of the captured molecules and is generally applicable to all barcoded features using within the same model. This generality results from the common phenomenological origin of the technical noise we aim to remove. To demonstrate this, we evaluate `CellBender` for denoising CITE-seq data. We treat cell surface protein and RNA measurements on an equal footing as a unified count matrix, and denoise the two modalities simultaneously using `CellBender`. Empirically, antibody counts exhibit a very high level of background noise which may be attributed to unbound and unwashed antibodies in the cell suspension. We show a publicly-available 10x Genomics CITE-seq dataset of PBMCs (`pbmc5k`) in Fig. 5. We have grouped antibodies together with their associated genes for ease of visual evaluation. In panel (a), the antibody features (red) have such a large amount of background noise that it is challenging to discern a clear pattern. The gene expression counts (blue), in contrast, have a very low amount of background noise in this dataset. Panel (b) shows the output of `CellBender` run with nFPR 0.1, where a pattern clearly emerges, and visually it appears that the red dots (protein antibody) very often line up with the blue dots (mRNA).

**Figure 5:**
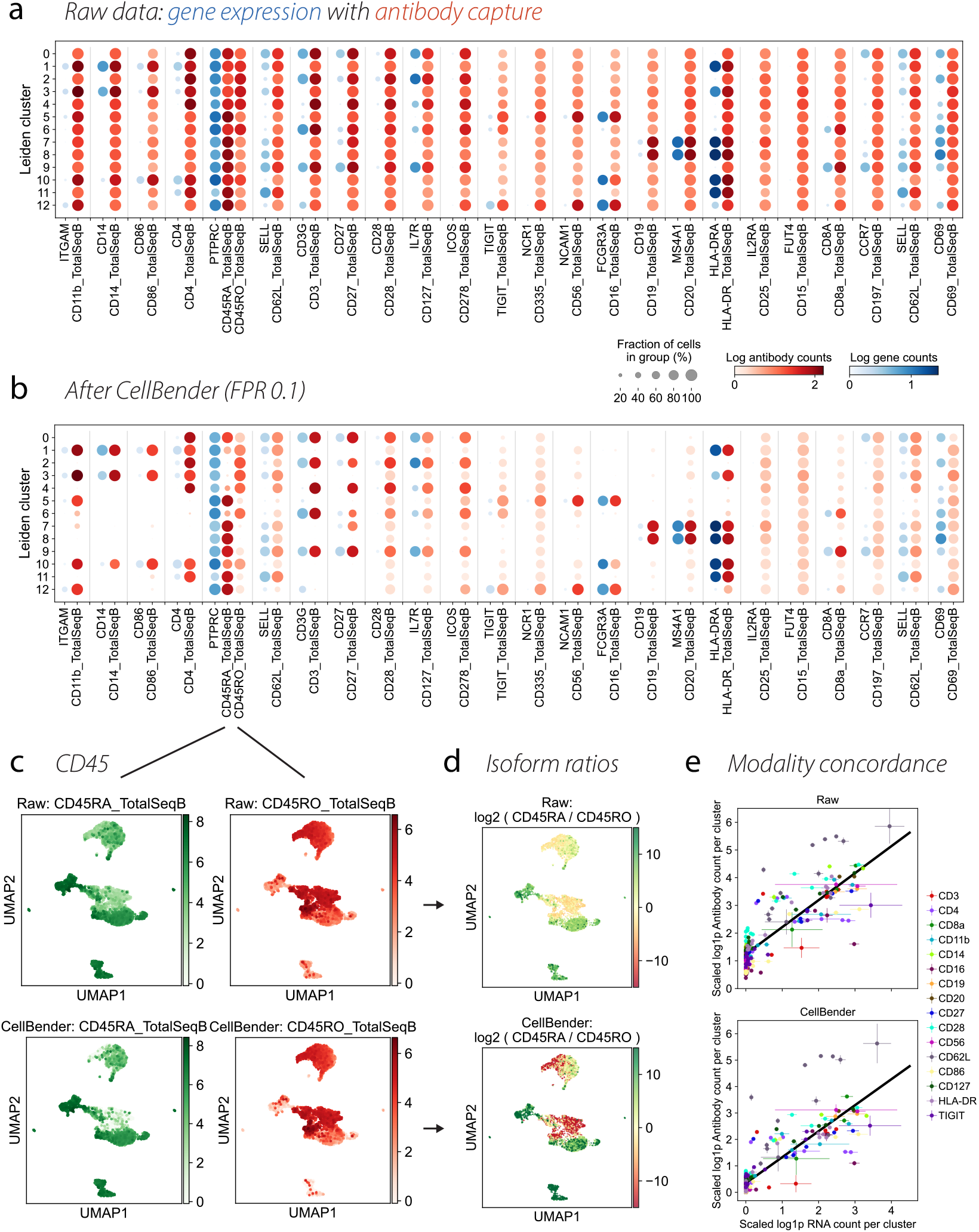
Performance of `CellBender` on denoising a CITE-seq PBMC dataset from 10x Genomics `pbmc5k`. (a) Raw data. The dotplot includes antibody capture features (red), along with the relevant gene expression features (blue) for all measured antibodies whose corresponding gene had max expression in any cell type above 0.05 mean counts. Groupings of related features are delineated by the gray vertical lines. (b) Same as (a) but for `CellBender` denoised counts. In both (a) and (b), the clustering is obtained at Leiden 0.6 resolution based on the `CellBender` output; see Supplementary Fig. S 15 for UMAP and cluster labels. (c) Examining CD45RA and CD45RO isoforms of CD45 as log normalized counts superimposed on the UMAP embedding. The expected anti-correlation of the two isoforms is significantly enhanced by `CellBender`. (d) UMAP embedding showing the log ratio of CD45RA and CD45RO expression and indicating the increased specificity afforded by `CellBender`. (e) Comparing the relationship between antibody counts and gene expression after scaling to collapse all data to the same line (see Sec. S.1.13 in Online Methods) for the raw data (upper) and `CellBender` denoised data (lower). By removing background counts, `CellBender` moves the intercept down toward zero and makes antibody counts more cluster-specific.

**Figure 6:**
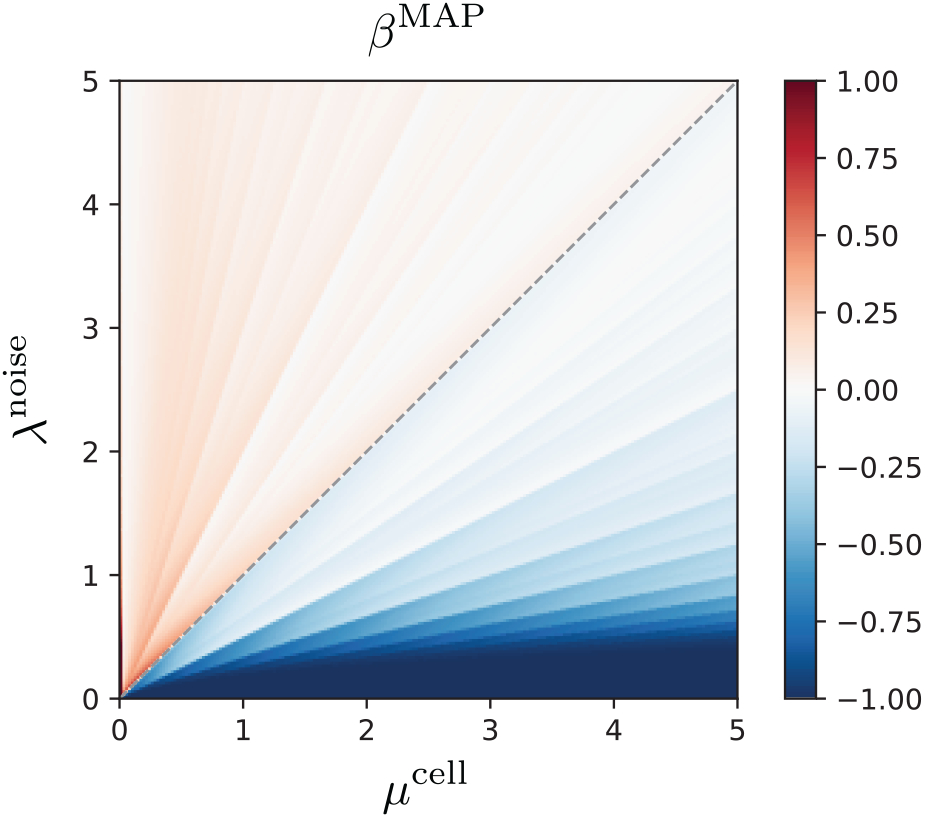
The relative asymptotic bias *β*^MAP^ of the MAP estimator of noise counts for a Poisson model of noise and cell counts. The axes show the respective prior rates. Note the extreme asymptotic bias *β*^MAP^ ≈ – 1 corresponding to estimating zero noise counts in the regime λ^noise^ ≪ *μ*^cell^.

Antibody counts and the corresponding RNA counts exhibit an expected linear relationship for most antibodies, and the impact of `CellBender` on this relationship is shown in Fig. 5e. In the raw data, the presence of background noise leads to a relatively large nonzero intercept, such that cells with zero RNA counts have nonzero antibody counts. `CellBender` effectively reduces the magnitude of this intercept while maintaining the biological linear relationship; additional results are given in Supplementary Fig. S16a-b. The specificity of antibodies to particular cell types improves as a direct consequence. Supplementary Fig. S16c shows that the Pearson correlation between the fraction of cells per cluster expressing antibody and the corresponding RNA increases markedly after `CellBender`. We note that the presence of large intercepts poses a challenge for comparing cell types across different batches and datasets, which may have different levels of background counts.

As a specific case study, we highlight two antibodies for different isoforms of *PTPRC* (also known as CD45): CD45RA and CD45RO, shown with the corresponding mRNA *PTPRC*. The removal of background noise (Fig. 5c) highlights a clear pattern of mutually exclusive differential expression of the two isoforms in different immune cell types: compare panel (d) top (raw) and bottom (`CellBender`). The expression of *HNRNPLL*, a splicing factor associated with the CD45RO isoform [44], is shown in Supplementary Fig. S15. We find that effector T cells states, i.e. T CD8+ EM/TE, and T regs, have both relatively higher level of *HNRNPLL* and CD45RO expression, as expected. `CellBender` increases the relative enrichment of CD45RO in such clusters as shown in Fig. 5d.

## 4 Discussion

We presented `CellBender`, an unsupervised method for removing systematic background noise from droplet-based single-cell experiments. `CellBender` learns the profile of noise counts from the data and subsequently estimates denoised counts. This is achieved by leveraging a deep generative model of noisy single-cell data that combines the flexibility of deep neural networks for learning the landscape of cell states with a structured probabilistic model of noise generation processes. `CellBender` can be used as a pre-processing step in any droplet-based single-cell omics analysis pipeline that involves an unfiltered count matrix, and is especially helpful for analysing datasets severely contaminated with background noise. These include snRNA-seq experiments that are subject to hash nuclear isolation protocols, and CITE-seq experiments that may produce large amounts of ambient antibodies. Removal of ambient noise has been advocated as an important step in single-cell analysis workflows and protocols [45, 46] and is increasingly becoming a standard part of single-cell data analysis.

Other authors have addressed the removal of background noise in scRNA-seq datasets in the past few years, including DecontX [32] and SoupX [11] for removal of ambient RNA, and methods for attenuating background counts due to chimeric molecules [13]. In practice, the operation of SoupX involves manual input and relies on the user’s prior knowledge of cell-type-specific gene expression, as well as providing (or calculating) a list of genes for estimating background RNA fraction in cells. The method introduced in Ref. 13 leverages read-per-UMI frequency data to detect library PCR chimeras. While this approach is highly effective at reducing the number of chimeric counts, it cannot detect physically encapsulated ambient molecules, which are indistinguishable from cell-endogenous molecules based on read-per-UMI frequency data alone. DecontX represents an unsupervised alternative for background noise removal. We have demonstrated that `CellBender` operates near the theoretically optimal limit and surpasses the performance of DecontX on several benchmarks. Other practical advantages of `CellBender` over DecontX include a tunable “nominal false positive rate” parameter for controlling the trade-off between denoising sensitivity and specificity in a principled fashion; automatic probabilistic determination of cell-containing droplets; and generation of a low-dimensional latent space embedding of cells which can be used in downstream analyses.

`CellBender` makes no assumptions about the profile of ambient noise and infers it from observed data, including counts in cell-free droplets. The analysis accompanying Fig. 2e,m demonstrated that studying the ambient profile produced by `CellBender` might be of value in and of itself, and could be used for instance to study the transcriptional makeup of extracellular vesicles, and to diagnose degraded and uncaptured cells. Ziegler *et al*., for example, made use of the `CellBender`-inferred ambient profile to help call high-confidence SARS-CoV-2 RNA^+^ cells in a scRNA-seq study of human nasopharyngeal swabs [28].

By mitigating background noise, `CellBender` eliminates a source of batch variation and spurious differential expression signals. This is particularly important for performing differential analysis of similar cell types between samples in a cohort. Since the systematic background noise is dataset-specific and is influenced by the circumstances around each batch, unmitigated noise can then appear as differential signal across batches. Sec. S.6.2 in Supplementary Methods includes a clear demonstration of this phenomenon and shows how `CellBender` effectively mitigates this source of batch variation and spurious differential expression. Removing systematic noise from individual datasets prior to integration is becoming increasingly crucial as the field is progressing from homogeneous small-scale experiments toward large-scale data integration and atlasing efforts, where datasets from many batches and tissue processing centers are being combined and analyzed jointly, see e.g. Eraslan *et al*. [31].

Field applications of `CellBender`, which include aiding the discovery of novel biology and resolving inconsistent findings, can be found in the works of other authors who have adopted our method since the time it was made publicly available as open-source software in 2019. We would like to highlight Ref. [47] where `CellBender` was applied to remove ambient RNA from brain snRNA-seq samples, resulting in the removal of neuronal marker genes from glial cell types, and identification of previous annotations of immature oligodendrocytes as potentially glial cells contaminated with ambient RNAs. A selection of other works which have used `CellBender` include primary research articles on the mouse brain [18], human brain organoids [19], human intestine [20], human heart [21–23], human and mouse adipocytes [24, 25], several recent studies on SARS-CoV-2 in human tissues [26–30], and a large snRNA-seq human cross-tissue atlas [31]. For particularly compelling example figures demonstrating the effects of `CellBender`, see Supplementary Fig. 1 in Eraslan *et al*. [31] and Extended Data Fig. 1e-h in Delorey *et al*. [26]. In cases where the raw data are relatively clean to begin with, Di Bella *et al*. observe that processing with `CellBender` will (appropriately) change the count matrix very little [48].

Future research directions include extending `CellBender` beyond the count matrix of unique UMIs, and modeling the data at the finer granularity of individual sequenced reads. For instance, chimeric reads can be identified much more effectively when read-per-UMI counts are taken into account [13]. This information is not contained in the conventional primary quantification of single-cell data as a count matrix of unique UMI counts. Additional interesting directions include evaluating the utility of `CellBender` on additional single-cell data modalities, including Perturb-seq [6] where background CRISPR guides can make the determination of perturbation challenging.

## Acknowledgements

The authors thank Luca D’Alessio, Carolina Roselli, Caroline Porter, Eli Bingham, Fritz Obermeyer, James Nemesh, Brice Wang, Behtash Babadi, Victoria Popic, Alec Wysoker, Ayshwarya Subramanian, Nathan Tucker, Yossi Farjoun, Timothy Tickle, and Ambrose Carr for insightful discussions at various stages of this project. S.J.F., M.D.C., and M.B. acknowledge financial support from the Broad-Bayer Precision Cardiology Lab (PCL). M.B. acknowledges additional support from the SPARC Grant *Development of Production-Grade Computational Methods for Single-Cell Genomics* from the Broad Institute. The (as yet) unpublished `rat6k` snRNA-seq dataset was generated by the PCL, and the experiment was performed by A.A. and A-D.A.

## S Supplemental Material

### S.1 Online Methods

#### S.1.1 Why a deep generative model?

Before we take a deeper dive into the `CellBender` model and inference algorithm, we would like to clearly motivate our choice of modeling framework. The approach taken here, i.e. deep generative models and stochastic variational inference (SVI), typically requires more computational resources than conventional deterministic algorithms and thus, must be conceptually justified.

First, we note that since the ambient molecules are aliquoted from the same cell suspension, they correspond to the same fixed distribution, and our many observations of cell-free droplets provide sufficient statistics to make it possible to infer that distribution with very high accuracy–in principle. In challenging cases such as highly contaminated snRNA-seq experiments where background noise removal is most needed, cell-free droplets are defined only in relation to cell-containing droplets (see Sec. 3.2). Therefore, we are obligated to model the landscape of cell feature counts (mRNA, protein, etc.) on par with the fixed distribution of ambient molecules. Cell states, however, are typically much more variable than the fixed distribution of ambient molecules. The challenging issue is our lack of *a priori* knowledge of the process that generates true biological counts in a cell, and the *a priori* unknown biological complexity of the assayed sample.

Furthermore, the fraction of captured mRNAs and other targeted features is on the order of 10% or less of expected counts (using 10x Genomics v2 or v3 chemistry, which generates approximately tens of thousands of feature counts per cell). Such sparse sampling is referred to as “dropout” in the context of droplet-based cell assays. For our purposes, dropout poses a particularly difficult challenge: even if we are provided with the knowledge of the true distribution of ambient molecules and other systematic background noises, “deconvolving” the observed count data from any given droplet into noise and signal contributions is a non-trivial task, given that both contributions are deep in the discrete regime and are subject to extreme sampling stochastic noise. We must, therefore, come up with a *prior* estimate of both contributions. An imbalanced model, e.g. one that has a stronger prior for noise and weaker prior for signal, or vice versa, will lead to over- or under-estimation of noise.

For these two main reasons, i.e. (1) *a priori* unknown landscape of cell states, and (2) sparse sampling of the content of each droplet (dropout), we are naturally led to a modeling choice that includes the following ingredients: (1) a flexible class of distributions to learn the landscape of cell states; (2) the ability to allow cells to share statistical power and leverage the observation from all cells to act as a prior; (3) the ability to automatically determine whether or not a droplet contains a cell.

Grouping of cells into clusters in order to share statistical weight may be achieved in multiple ways, including a nearest-neighbors clustering (as in a traditional scRNA-seq analysis) and other graph-based methods [49]. Using information learned from similar cells to build a *prior* belief is most rigorously done within the Bayesian framework. Bayesian methods for modeling complex distributions include auto-encoders and normalizing flows. Finally, automatic determination of cell-free vs. cell-containing droplets requires model comparison which may also be rigorously done within the Bayesian framework. We have found the common denominator of these requirements, together with the expressibility of the Bayesian framework for turning mechanistic insights into structured probabilistic models, to naturally lead to a model that is no more or no less complex than `CellBender`.

#### S.1.2 Model

Our generative model for noisy droplet-based count data is shown in Fig. S1a, along with a schematic of the rationale in Fig. 1g. Throughout this section, we use *n* and *g* subscripts to refer to cell and molecular feature (e.g. gene, protein) indices on various vector and matrix variables. 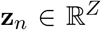 is the latent variable that encodes endogenous cell states in a lower-dimensional space. *χ_ng_* is the fractional molecular feature frequency (i.e. normalized to 1) in cell *n*, and lives on a (*G* – 1)-simplex in 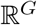, where *G* is the dimensionality of the raw molecular feature space (e.g. number of genes in scRNA-seq). NN_χ_, shown as a factor (black square) in the graphical model, is the “decoder” neural network that deforms the low-dimensional embedding **z**_*n*_ to the raw data feature space *χ_ng_*. 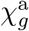 is the normalized abundance of ambient molecules and is a learnable parameter. 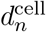 is a cell-specific size factor. 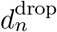 is a droplet-specific size factor for ambient counts. *y_n_* is a discrete binary random variable which is 1 if there is a cell in droplet *n* and 0 otherwise. *ρ_n_* is the proportion of reads that are assigned to droplet *n* but are exogenous to droplet *n* and have been randomly swapped e.g. due to PCR chimera formation. *ϵ_n_* is a droplet-specific capture efficiency parameter, close to 1, that reflects how efficiently the targeted molecules in droplet *n* are captured, barcoded, and reverse-transcribed. In other words, *ϵ_n_* is a technical confounder that affects the total UMI counts in a droplet, endogenous and ambient alike. 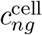 and 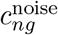 denote the latent counts per droplet that come from the cell and from background sources, respectively. Finally, *c_ng_* is the observed counts of feature *g* in cell *n*. The generative process is as follows:

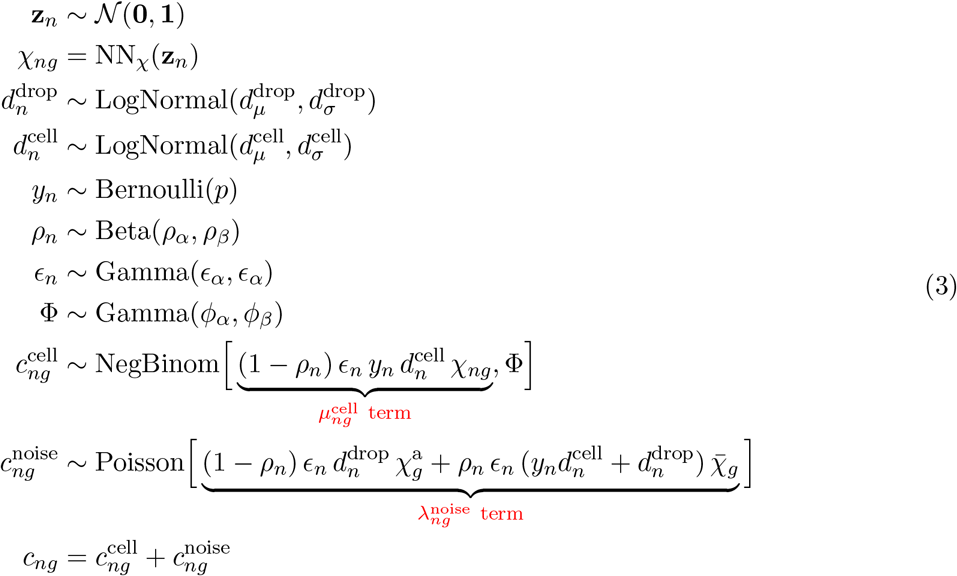

**Figure S 1:**
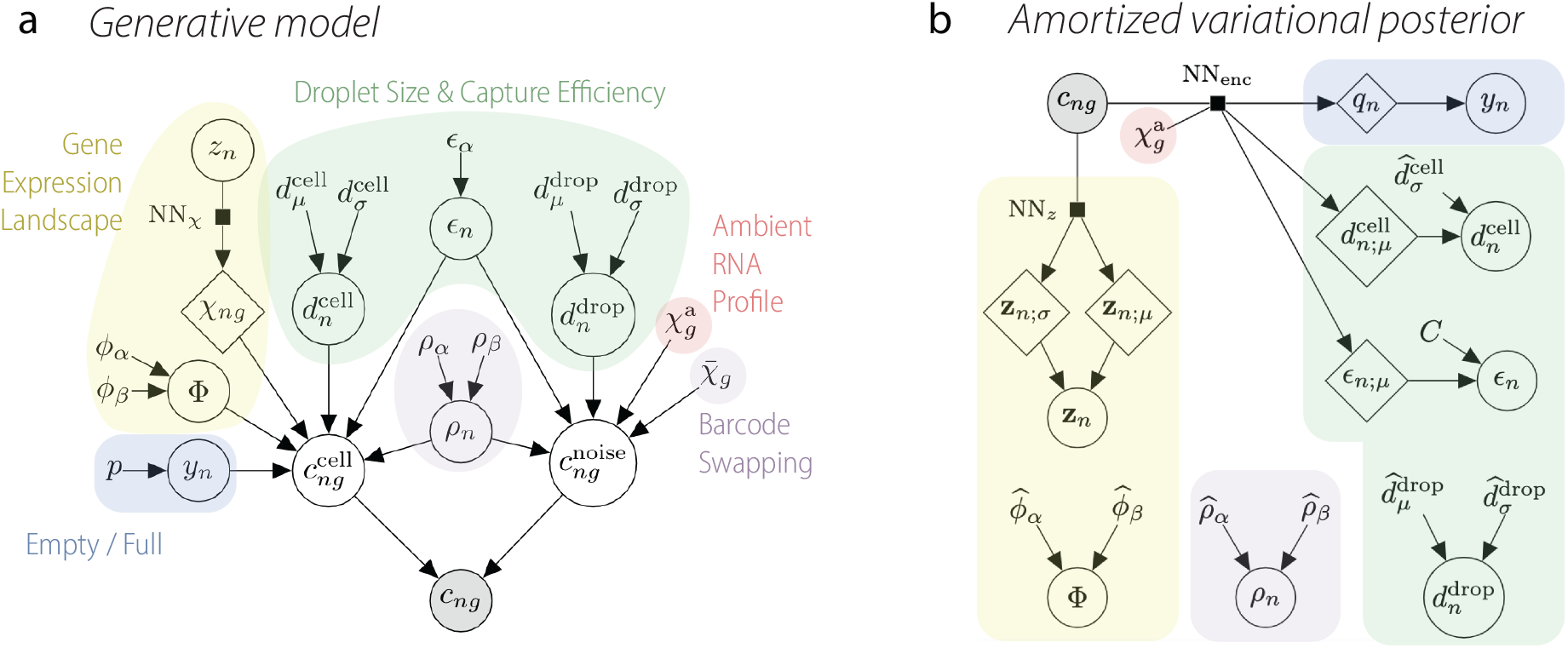
(a) `CellBender` generative model for noisy single-cell count data. (b) The variational posterior used by `CellBender`. The neural network NN_enc_ takes the observed data as input and yields the parameters of various variational distributions assumed for the local latent variables. The global latent variables are treated in the usual mean-field approximation.

##### Modeling the rate of endogenous and exogenous feature counts

We will discuss our parametric choices for count likelihoods, i.e. negative binomial for endogenous counts and Poisson for ambient counts, in the next section. Here, we focus on the expressions given for the “rates” of the two contributions, 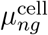 and 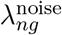, respectively. The rate of endogenous counts in a droplet, 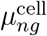, straightforwardly follows from the definitions: 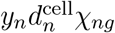 represents the expected counts from the cell in droplet *n*. The rate is modulated by the droplet’s efficiency *ϵ_n_*, and the term (1 – *ρ_n_*) is the fraction of library fragments originating from the cell that are *not* swapped to a different droplet, maintaining the interpretation of *ρ_n_* as the fraction of swapped counts exogenous to droplet *n*. The rate of exogenous counts in a droplet, 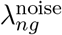, has two parts: ambient molecules and randomly swapped barcodes. The barcode swapping process results in a certain fraction of counts in each droplet, *ρ_n_* ∈ [0, 1], having actually originated in other droplets. We assume it is equally likely to swap any two barcodes, and so the net effect is that the swapped molecules in any given droplet are effectively sampled from the average (“bulk”) features over the entire experiment, denoted by 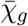. Ambient molecules, on the other hand, may have a distinct composition as argued in Supplementary Section S.2 and demonstrated in Section 3.1, and therefore are sampled from a different and learnable profile, denoted by 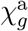. Accordingly, we decompose the rate into two main parts. The first part is the ambient counts that physically originate in droplet 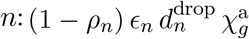. The second part is the counts that did not physically originate in droplet n, but were erroneously assigned there later: 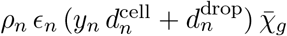. This expression is the product of three terms: the contamination fraction *ρ_n_*, the term in parentheses together with *ϵ_n_* that is proportional to the expected number of molecules physically encapsulated in the droplet, and finally the average (“bulk”) molecular profile, 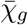.

##### Count Likelihood Models

The fundamental noise governing count data in single-cell sequencing is Poisson, rooted in the empirical fact that each molecule has only a small probability of being successfully captured and sequenced. We refer the reader to the excellent analysis of Refs. [50, 51] on this matter and the nuances and hazards of employing more flexible count likelihood models.

Accordingly, we model the noise statistics of background noise counts, 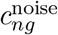, as a Poisson distribution. We do not accommodate for additional overdispersion in addition to what is implicitly induced by the stochasticity of the latent variables that appear in the Poisson rate of exogenous counts, see Eq. (3): we believe our theoretical model of ambient counts and barcode swapping to be flexible enough, and to be a fairly faithful representation of the simple underlying physical process, such that any additional overdispersion is likely to result in model under-specification.

On the other hand, we purposefully endow endogenous counts 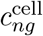 with extra overdispersion, signified by the overdispersion parameter Φ of a negative binomial (Poisson-Gamma) distribution. In the context of our problem, this inclusion is motivated as follows: as mentioned earlier, imposing a prior distribution over 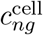 is to provide a mechanism to share statistical power across cells, help overcome data sparsity, and ultimately aid deconvolving observed counts into exogenous and endogenous compartments. Crucially, the prior imposed on endogenous counts must be data-driven and endowed with a tunable parameter to balance the model’s prior belief over endogenous counts with exogenous counts, as dictated by the structure of the data and the maximum likelihood principle that we use to fit the model. The extra overdispersion parameter provides precisely such a mechanism to balance the prior beliefs and desensitize the results on the representational capacity of the underlying neural networks that encode the structure of endogenous counts. Faced with a dataset that contains a large number of the same cell types in the same state, the model will benefit from reducing Φ and strengthening its prior belief of endogenous counts. By contrast, prior belief will be commensurately “softer” when faced with a complex dataset, in particular, if the size of the latent space is not large enough to afford the complexity of the dataset.

##### Model Hyperparameters

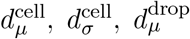, and 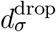 are all fixed hyperparameters that we determine automatically from the provided data using a number of heuristics. A cutoff in UMI counts (`--low-count-threshold`) is used to remove very low UMI count barcodes. The mode of the remaining UMI count distribution is then used to approximate 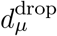. A gaussian mixture model is fit to the UMI counts per droplet, and mixture components larger than 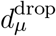 are identified and combined to obtain an estimate of 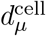. The variance hyperparameters are also estimated from the gaussian mixture components, and scaled down to account for the dispersion induced by *ϵ_n_*. These hyperparameters specificy the prior for endogenous and ambient rate scale factors, 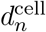 and 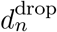, both of which are modeled as LogNormal distributions on an empirical basis. *p* is a hyperparameter representing the prior probability that any given droplet contains a cell, and it is derived from the expected number of cells in the experiment and the total number of droplets included in the analysis. (*ρ_α_, ρ_β_*) are general priors for the contamination fraction *ρ_*n*_*, with default values of (1.5, 50), motivated by the fact that the shape of this beta distribution matches our expectations, from observations of many datasets, that barcode swapping is typically in the range of a few percent. The hyperparameter *ϵ_α_* controls how concentrated the droplet-specific capture efficiency will be around 1. We use a fixed value of 50, motivated by examination of the overdispersion of the droplet sizes in the 10x Genomics `ercc` dataset, compared to a Poisson.

##### Choice of Contamination Model

The `CellBender` model can be restricted to only ambient background noise by setting *ρ_n_* = 0 for all *n*, or it can be restricted to barcode swapping background noise only by removing the “endogenous ambient” term, 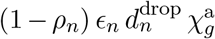, from the Poisson rate for 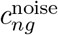. The default mode in `CellBender` uses the full model as specified in Eq. (3), but the user can specify the ambient-only or swapping-only model via command line arguments in our provided implementation.

#### S.1.3 Inference

The probabilistic model described in the previous section entails several global (experiment-wide) and local (one for each droplet) latent variables. Scalable approximate inference can be achieved using stochastic variational inference (SVI) [52] and amortization [53]. We provide a brief account of the inference strategy in this section. We note that other authors have also successfully applied SVI techniques for scalable probabilistic modeling of single-cell data [34–36]. The objective function which is optimized in SVI is the evidence lower bound (ELBO):

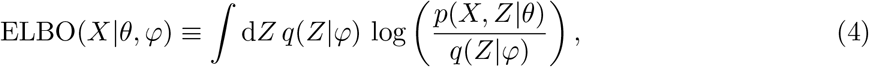

where *X* = {*c_ng_*} is the observed data, 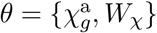 is the bundle of tunable model hyperparameters, including the weights of the neural network NN_χ_ (denoted by W_χ_), 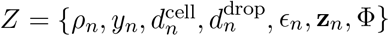 is the bundle of latent variables, and *q*(*Z*|*φ*) is the variational ansatz shown in Supplementary Fig. S1b and parameterized by 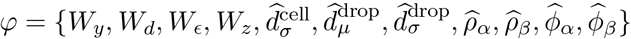. In the SVI methodology, one obtains argmax_*θ,φ*_ ELBO(*X*|*θ, φ*) via successive sub-sampling of data *X* and incremental updates of (*θ, φ*) using a stochastic optimizer. We refer the reader to Ref. 54 for a review.

##### Constructing a Variational Posterior Distribution

The faithfulness of the approximate posterior to the true posterior is ultimately dependent on one’s choice of the variational ansatz, *q*(*Z*|*φ*). Supplementary Fig. S1b shows the structure of our proposed ansatz. Generally speaking, we impose tunable parametric distributions over global latent variables while we infer local latent variables using auxiliary neural networks (often referred to as recognition or encoder networks). The latter technique is referred to as amortization and is the key to the scalability of our algorithm to a theoretically unbounded number of data points (cells).

The posterior for **z**_*n*_ is encoded by a neural network NN_*z*_ which takes in observed counts *c_ng_*, along with the current estimate of the ambient profile 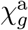, and outputs 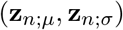; the latter parameterize the mean and scale of an assumed Gaussian posterior distribution for **z**_*n*_:

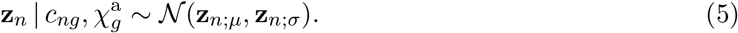

Note that this *encoder* network for **z**_*n*_, together with the *decoder* network that maps **z**_*n*_ to *χ_ng_*, form the auto-encoder structure mentioned earlier, in the spirit of Kingma *et al*. [33].

The variational posteriors for the cell presence indicator variable *y_n_*, cell scale-factor 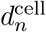, and the droplet-specific capture efficiency *ϵ_n_* are parameterized via additional neural networks (shown together as NN_enc_ in Fig. 1). These auxiliary encoder neural networks each take *c_ng_* and 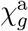 as input, and estimate all or some of the parameters of specified posterior distributions. In practice, we found it beneficial to further provide a few hand-crafted features constructed from *c_ng_* and 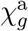 as inputs to each of the encoder neural networks (see Sec. S.1.11). The posterior for *y_n_* is assumed to be Bernoulli and is parameterized by the neural network NN_*y*_ that outputs *q_n_*, the cell presence posterior probability:

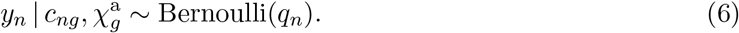

The posterior for 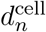 is assumed to be LogNormal and is parameterized by the neural network NN_*d*_ which outputs 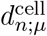, a strictly-positive scale-factor per droplet:

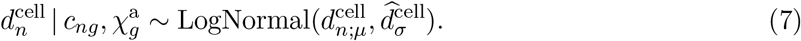

We have additionally introduced a learnable posterior parameter 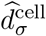 to characterize the uncertainty in estimating cell scale-factors. The posterior for *ϵ_n_* is assumed to be Gamma and is parameterized by the neural network NN_*ϵ*_ which outputs *ϵ_n;μ_*, the posterior mean capture efficiency:

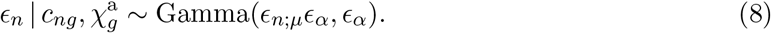

Here *ϵ_α_* is the same hyperparameter from the model, controlling the uncertainty in droplet efficiencies. Finally, the variational posteriors for Φ, *ρ_n_*, and 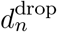 are assumed as follows:

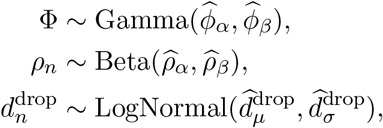

each of which involve two trainable parameters. Note that we have assumed the barcode-swapping rate *ρ_n_* and droplet size 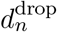 to have the same posterior distribution for all droplets *n*, even though these are droplet-specific (local) latent variables. We have found this more restrictive posterior to work well in practice while allowing more robust SVI fits.

##### Approximate treatment of Poisson and Negative Binomial convolution

Details aside, the structure of our generative model for endogenous and exogenous counts is as follows (see Eq. 3):

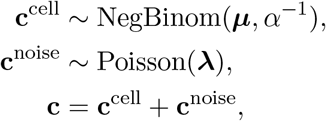

where we have dropped the common *ng* indices and used bold symbols as a shorthand for cell × feature matrices. Here, ***μ*** and **λ** refer to the endogenous and exogenous count rates, and *α* = Φ^−1^ is the inverse overdispersion. This parametric decomposition into non-negative endogenous and exogenous contributions ensures that the inferred endogenous counts **c**^cell^| **c** to be ≤ **c**. This desirable property, however, poses a technical challenge: as a part of variational inference, we need to be able to compute the probability density of **c** in a differentiable fashion; however, the sum of a general Poisson and a general negative binomial distribution does not admit a closed probability density expression. Formally, the latter is given by the convolution of the two probability densities. Computing this convolution explicitly, while doable, is prohibitively slow. We therefore resort to the following approximation during model training:

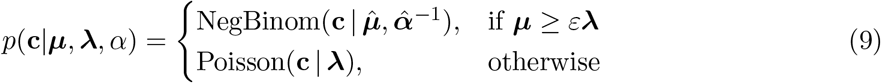

where we set *ε* = 10^−5^, and 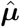 and 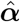 are obtained by matching the first two moments of an “effective” negative binomial distribution to **c**^cell^ + **c**^noise^:

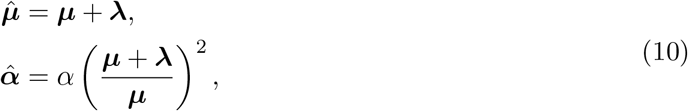

where all algebraic operations involving matrices are element-wise. The rationale for switching from a moment-matched negative binomial to Poisson when ***μ*** < *ε***λ** is for numerical stability: when ***μ*** → 0, i.e. vanishing prior rate of endogenous counts, we obtain 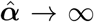, which leads to numerical instability. At the same time, the observed count is dominated by noise counts in this regime, i.e. 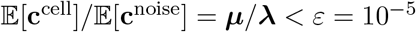, justifying the switch.

#### S.1.4 Constructing the denoised integer count matrix: preliminaries

Our Bayesian model, following fitting of model and posterior parameters, allows us to compute the posterior probability of having a specified number of noise counts in each entry of the count matrix. Even though we marginalize 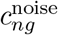 during inference, we can recover its posterior after model fitting via posterior sampling. We formally have:

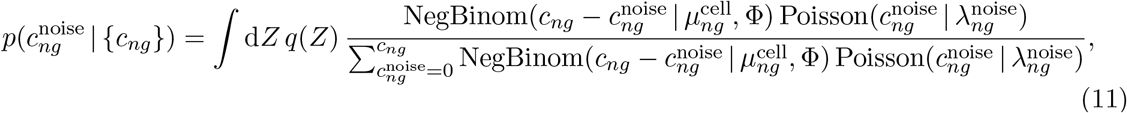

where *Z* is the bundle of all other latent variables along with their approximate posterior distribution *q*(*Z*). The terms and expressions appearing in the integrand are evaluated at *Z*. In practice, we approximate the integral via *N* discrete Monte Carlo samples drawn from *q*(*Z*) and keep track of the marginal posterior of noise counts for each of the *n* × *g* count matrix entries. We compute the probabilities in log space for numerical stability, truncate the allowed range of 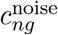 to a safe upper bound, normalize each MC sample via the LogSumExp operation, and keep track of the running total over MC samples via sequential LogSumExp operations for memory efficiency.

The obtained *n* × *g* marginal posterior distributions comprise our full probabilistic knowledge of noise counts for each entry in the count matrix. Standard single-cell downstream analysis workflows, however, by and large expect a single point estimate for input, as opposed to a distribution. Furthermore, a plurality of widely-used algorithms such as voom [55] for differential expression (DE) analysis, Seurat v3’s highly-variable gene (HVG) selection [56], and scVI [34] for latent space learning, explicitly expect *integer* counts as input due to employment of discrete likelihood models such as negative binomial. These expectations motivate us to estimate a single integer matrix of noise counts 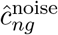 from the obtained Bayesian posterior 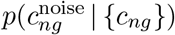, and produce an integer matrix of denoised counts 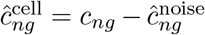 as the primary output of `CellBender`. The strict satisfaction of 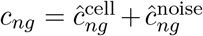 implies the complementary of noise and signal estimators. Hereafter, we focus on estimating the noise matrix for concreteness.

Canonical Bayes estimators for summarizing 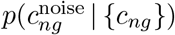 as a single point estimate include: (1) the posterior mean, 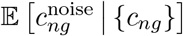, and (2) the posterior mode, argmax 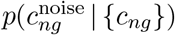, also known as the maximum *a posteriori* (MAP) estimator. The posterior mean estimator is an unbiased estimator; however, it yields non-integer values, which is undesirable. The MAP estimator yields integer values, however, is a biased estimator. For example, the MAP estimator systematically underestimates noise counts for genes that have lower noise prior rate compared to cell expression prior rate, see Sec. S.1.10. Neither of these canonical estimators provide a tunable parameter for increasing or decreasing the strength of denoising and controlling the trade-off between denoising sensitivity and specificity.

In order to address these shortcomings, we introduce a number of application-specific estimators to meet our specific needs. In general, we aim to develop estimation strategies for attaining the highest posterior probability subject to specified population-level constraints such as gene-wise or dataset-wise total noise budgets or expected false positive rate. Having such handles is useful in many downstream applications such as ascertaining the specificity of marker genes. We note that the true Bayesian recipe for conveying the results of `CellBender` is the full posterior and not a point estimate, and that the optimality an integer noise estimator is not universal and depends on the downstream application. For example, the desire to have an estimator suitable for differential expression testing between samples imposes a different set of constraints than the desire to have a given degree of certainty that each count in the output is not noise. We examine the merits and drawbacks of each strategy using different metrics in the following sections.

#### S.1.5 Estimating the integer noise matrix as a multiple-choice knapsack problem

We show that the problem of estimating an integer noise matrix 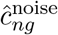 that attains maximum posterior probability subject to linear constraints is equivalent to the multiple-choice knapsack problem (MCKP), which is a classical combinatorial optimization problem. To set the stage, we assume a linear index map 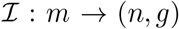 from *m* ∈ {1, …, *N* × *G*} to the entries of the count matrix (*n, g*), for 1 ≤ *n* ≤ *N* and 1 ≤ *g* ≤ *G*. Let 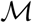 be the index set of noise count matrix elements we wish to perform constrained estimation over. Choices include the entire count matrix 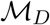, a row (cell) 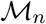, or a column (gene) 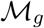. We define *X_mc_* ∈ {0, 1} to be a binary variable that is 1 if the noise count for matrix element at 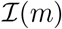 is set to *c*, and is 0 otherwise. Since there is a unique choice to be made for each matrix entry, we require 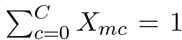, where *C* is the maximum specified noise count and is upper bounded by max *c_ng_*. We further define a “reward” for each assignment as 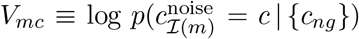, i.e. the log posterior probability for that assignment. Finally, we wish to impose a lower bound *L* on the sum total of noise counts. This is readily expressed as 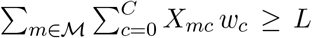, where *w_c_* = (0, 1, …,*C*) is an integer-valued weight vector. Maximizing the log posterior probability, which is given as 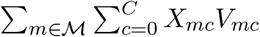 subject to the aforementioned constraints is expressed as:

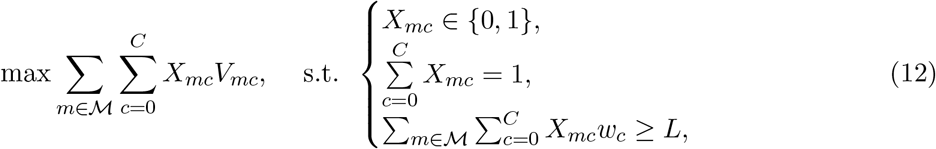

which is precisely the MCKP problem. MCKP is a classical NP-hard problem which admits a pseudo-polynomial dynamic programming solution. In our specific case, we show that subject to mild assumptions, a fast and exact solution is feasible with time complexity 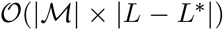, where 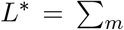 argmax_*c*_ *V_mc_*, see S.1.9. Note that *L*\* is the sum of MAP estimates over the specified count matrix entries 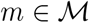. Since the noise rate is typically lower than the endogenous expression rate, *L*\* is typically an underestimate (see Sec. S.1.10) and as such, we are generally interested in cases where *L* > *L*\* to overcome the asymptotic bias of the MAP estimator. Moving away from the MAP estimator, by definition, decreases the posterior probability. As such, the inequality constraint is realized at the threshold *L* and thus, we refer to *L* as the “noise target”.

##### Concrete MCKP problems for enforcing gene-wise and dataset-wise noise count constraints

The MCKP framework allows us to impose noise targets over arbitrary selections of count matrix entries. For concreteness, we consider two scenarios: (1) imposing gene-wise constraints, where each column *g* of the noise count matrix is constrained to sum to ≥ *L_g_* and is estimated independently; and (2) dataset-wise constraints, where all count matrix entries are estimated at once subject to a global constraint that the sum total of noise counts ≥ *L*. Setting the noise target may also be done in a different ways. Here, we consider two strategies: (1) a noise target based on a nominal false positive rate (nFPR), and (2) a noise target based on the cumulative distribution function (CDF) of the posterior of the aggregated noise counts. These strategies are described below.

##### Using the nominal false positive rate (nFPR) to specify the noise target

We introduce a single tunable parameter nFPR ∈ [0, 1] to specify the noise target. We define this parameter such that nFPR = 1 implies allocating all raw counts as noise counts whereas nFPR = 0 implies removing as many noise counts as what is inferred from the model posterior aggregated over the appropriate slice of the dataset, i.e. either gene-wise or for the full dataset. Specifically, we define nFPR as follows. For each gene *g*, we estimate the expected noise count per likely cell-containing droplet as follows:

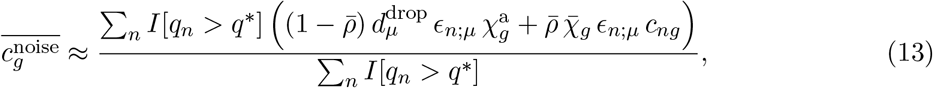

where *q*\* = 0.5 is the threshold we have chosen for determining likely cell-containing droplets, and 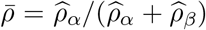 is the posterior mean of the barcode-swapping rate. The two terms in the numerator correspond to ambient and barcode-swapping contributions to noise counts. Likewise, we estimate cell counts as follows:

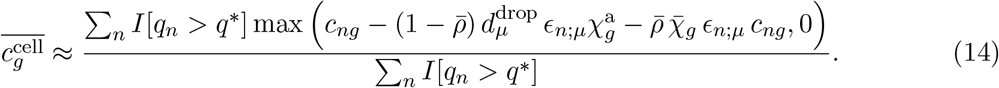

Equipped with these two aggregate estimates, we define the nFPR recipe for specifying the per-cell per-gene noise target ℓ_*g*_ as:

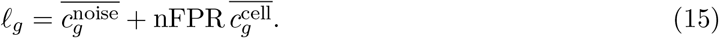

The gene-wise total noise target for *N* cells is given as *L_g_* = *N ℓ_g_*, and the dataset-wise total noise target for *N* cells is given as 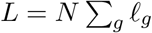.

##### Using the aggregated noise posterior CDF quantiles to specify the noise target

Another strategy for setting a total noise target over a slice of the dataset is via the quantiles of the the aggregated noise posterior. The aggregated noise over the desired set of count matrix entries 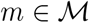 defined as:

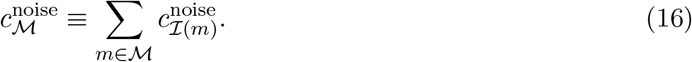

The posterior distribution of 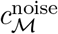 is formally given as the convolution of the posterior distribution of the included noise count matrix entries. The latter can be obtained numerically using fast Fourier transform (FFT). In practice, we have found that calculating the first two moments of 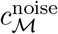 and appealing to the central limit theorem (CLT) yields virtually identical results. These moments are given as:

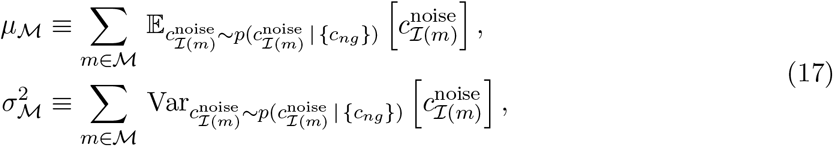

and the CLT implies 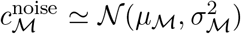. Given a total noise CDF quantile *q*, we set the noise target to:

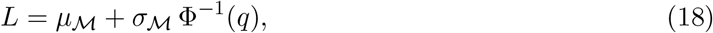

where Φ^−1^(*q*) is the inverse CDF of the normal distribution. Like before, we can set 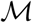 to either 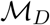 or 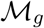 for imposing dataset-wise or gene-wise noise targets, respectively.

#### S.1.6 Estimating the integer noise matrix via element-wise noise posterior CDF quantiles

A straightforward strategy for estimating the integer noise count matrix is to pick the noise count for each entry of the noise count matrix according to a specified CDF quantile *q*. In particular, the choice *q* = 0.5 corresponds to the posterior median estimator, which is a canonical Bayes estimator. Specifying a higher (lower) valuer for *q* results in removing more (fewer) noise counts and as such, *q* serves as a handle for setting the denoising eagerness of `CellBender`. This algorithm is implemented as follows. For each cell *n* and gene *g*, we calculate the CDF of noise counts 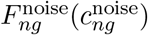 from the noise posterior:

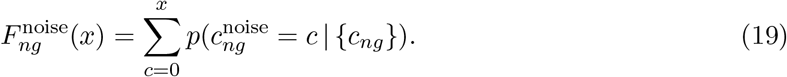

The estimated integer noise count matrix is then obtained as:

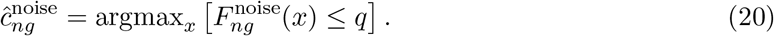

This estimator, as opposed to the MCKP approach discussed in the previous section, does not involve solving a global constrained optimization problem and as such, does not allow targeting noise counts in aggregate, either gene-wise or dataset-wise. While it is possible to fine-tune the quantile threshold *q* to achieve the desired nFPR, we did not attempt it: MCKP achieves the same goal by allocating the total noise budget more globally rather than locally, and as such, can achieve a higher total posterior probability.

#### S.1.7 Estimating the integer noise matrix via posterior regularization

Another strategy for estimating an integer noise matrix subject to external constraints, such as dataset-level or gene-wise nFPR, is provided by the framework of posterior regularization of Ganchev *et al*. [57] and is another optimization-based approach. This is the framework we had adopted in `CellBender` v0.2.0 and we provide it here for completeness. Concretely, following the setup of Eq. (4) in Ganchev *et al*. [57] (with no slack, i.e. *ε* = 0), given data *X* = *{c_ng_*} and latent variables *Z*, we seek a posterior distribution 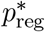 that solves the following constrained optimization problem:

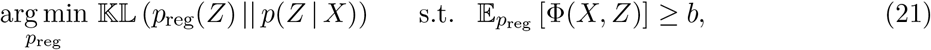

where *p*(*Z*|*X*) is the unregularized Bayesian posterior, *p*_reg_(*Z*) is the sought-after regularized posterior, and Φ(*X*, *Z*) is a specified function of raw data and latent variables that we wish to constrain below a specified value of *b* in expectation. We have implicitly grouped the model parameters together with the latent variables in *Z*. Adapted to our problem, we wish to compute a regularized posterior for noise counts, 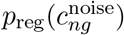, such that it is as close as possible to the regularized posterior in terms of KL divergence while the expected total noise count over all likely cell-containing droplets is controlled by the user-specified nFPR parameter (see Eq. 15):

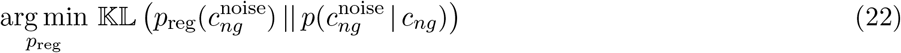

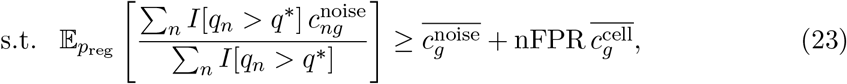

where *q*\* = 0.5 is the posterior probability threshold we have chosen for likely cell-containing droplets. As it is written, the nFPR condition is imposed separately for each gene *g*. A more relaxed version of the problem is obtained by summing both sides of the constraint over *g*, which is equivalent to imposing a dataset-wise constraint. In the dual formulation [57], the regularized posterior that satisfies Eq. (21) can be written as:

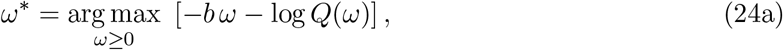

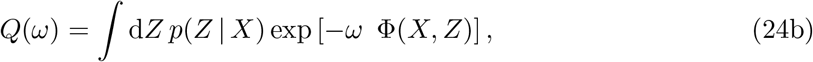

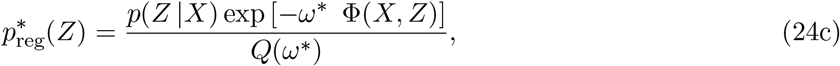

and the problem is reduced to finding an appropriate *ω\** that satisfies the constraint imposed by *b* and Φ(·). Exact posterior regularization requires separate SVI model fits for every choice of constraint threshold (*b* in Eq. 22). In theory, one could solve the optimization problem posed by Eqs. (24a)–(24c) in dual form, plugging 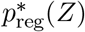 into the ELBO (Eq. (4)), and interleaving SVI updates with constrained satisfaction updates: a computationally prohibitive task. Another approach is augmented Lagrangian constrained optimization, where one concurrently updates *ω* along with model parameters using the same stochastic optimizer to minimize the ELBO while also approximately satisfying Eq. (24a).

Here, we make an approximate simplifying assumption akin to perturbation theory: so long as the user does not impose extreme values of expected nFPR compared to the FPR achieved in the unregularized problem, then we expect all latent variables to remain approximately the same, with and without posterior regularization, with the exception of perhaps 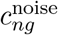, which directly appears in the constraint. By employing this approximation, we can freeze all latent variables to their unregularized posteriors and only regularize 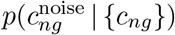 *ex post facto*. To achieve this goal, consider scaling 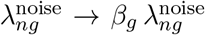, where 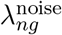 is the Poisson rate of exogenous counts given in Eq. (3) and *β_g_* ≥ 0 is a to-be-determined scale factor. We postulate that finding the optimal scale factor that satisfies the posterior constraint is equivalent to solving Eqs. (24a)–(24c). To show this, we use the following identity which can be readily ascertained using the explicit expression of the Poisson probability mass function:

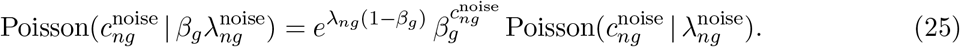

According to the dual formulation given in Eq. (24a)–(24c), we can write 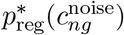 for likely cell-containing droplets, i.e. *q_n_* > *q*\*, as:

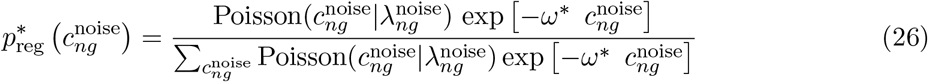

Comparing this to Eq. (25), we identify 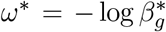. In other words, solving for the regularized posterior reduces to the problem of finding the largest noise scale factor 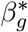 that satisfies the constraint in Eq. (22). The regularized Poisson rate for noise counts is then 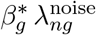. For a dataset-level nFPR constraint, the gene-wise scale factor 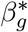 reduces to a single global scale factor *β*\*. At the moment, only the dataset-level nFPR condition is implemented in `CellBender`.

##### Locating the optimal *β*\* via binary search and estimating the integer noise matrix

We locate the optimal noise scale factor *β*\* numerically using a binary search strategy. Our goal is to identify the largest value *β*\* such that the inequality given in Eq. (22) is satisfied. Binary search is performed over the range *β*\* ∈ [0.01, 500]. At each iteration of the search, we estimate 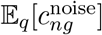 by obtaining the regularized posterior using Eq. (11) and making the replacement 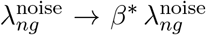. For computational efficiency, we only include a random subset of likely cell-containing droplets (128 randomly-chosen cells by default). The entire optimization procedure is repeated five times using different randomly-chosen subsets of cells. The final value of *β*\* is the average from the several repeats. Having located the optimal *β*\*, we obtain the integer noise count matrix as the MAP estimate from the regularized noise posterior. We refer to this noise estimation strategy as posterior regularization for mean-targeting, or “PR-*μ*” for short.

##### Approximate noise CDF quantile targeting via posterior regularization

A variation of the discussed PR strategy is obtained by replacing the constraint appearing in Eq. (22) with the following:

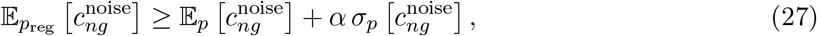

where *α* = Φ^−1^(*q*) is approximately equal to quantile *q* of noise under the normality assumption. Note that the constraint is imposed at the level of individual count matrix entries. The motivation for this approach is to allocate the extra noise budget preferentially to the count matrix entries with lower noise posterior confidence. Again, the dual form of the PR problem implies a solution 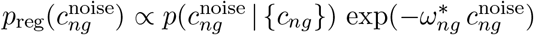 where 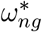 is a matrix of Lagrange multipliers to be determined in order to satisfy Eq. (27). In practice, we obtain 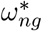 by performing a parallelized binary search as described earlier. Once the regularized posterior is obtained, the output can be summarized either by taking the posterior mean, the posterior mode, or a single sample, all of which we compare later. Note that *α* = 0 is identical to the unregularized posterior. We refer to this noise estimation strategy as posterior regularization for quantile-targeting, or “PR-q” for short.

#### S.1.8 Evaluating different noise estimation strategies

We introduced several strategies for estimating noise counts from the Bayesian noise posterior in Sec. S.1.5–S.1.7 in order to address the shortcomings of canonical Bayes estimators and allow controlling the denoising sensitivity-specificity trade-off. In this section, we evaluate these strategies on a simulated dataset that closely follows our model, see S.1.14. Concretely, we generate a test dataset consisting of three “cell types” with fixed gene expression profiles. We generate 100 cells of each type with 5000 UMIs/cell on average, and a background noise that consists of only ambient RNA for simplicity. The ambient RNA profile is taken to be the same as the average gene expression across all simulated cells, with 200 ambient UMIs per droplet on average.

**Figure S 2:**
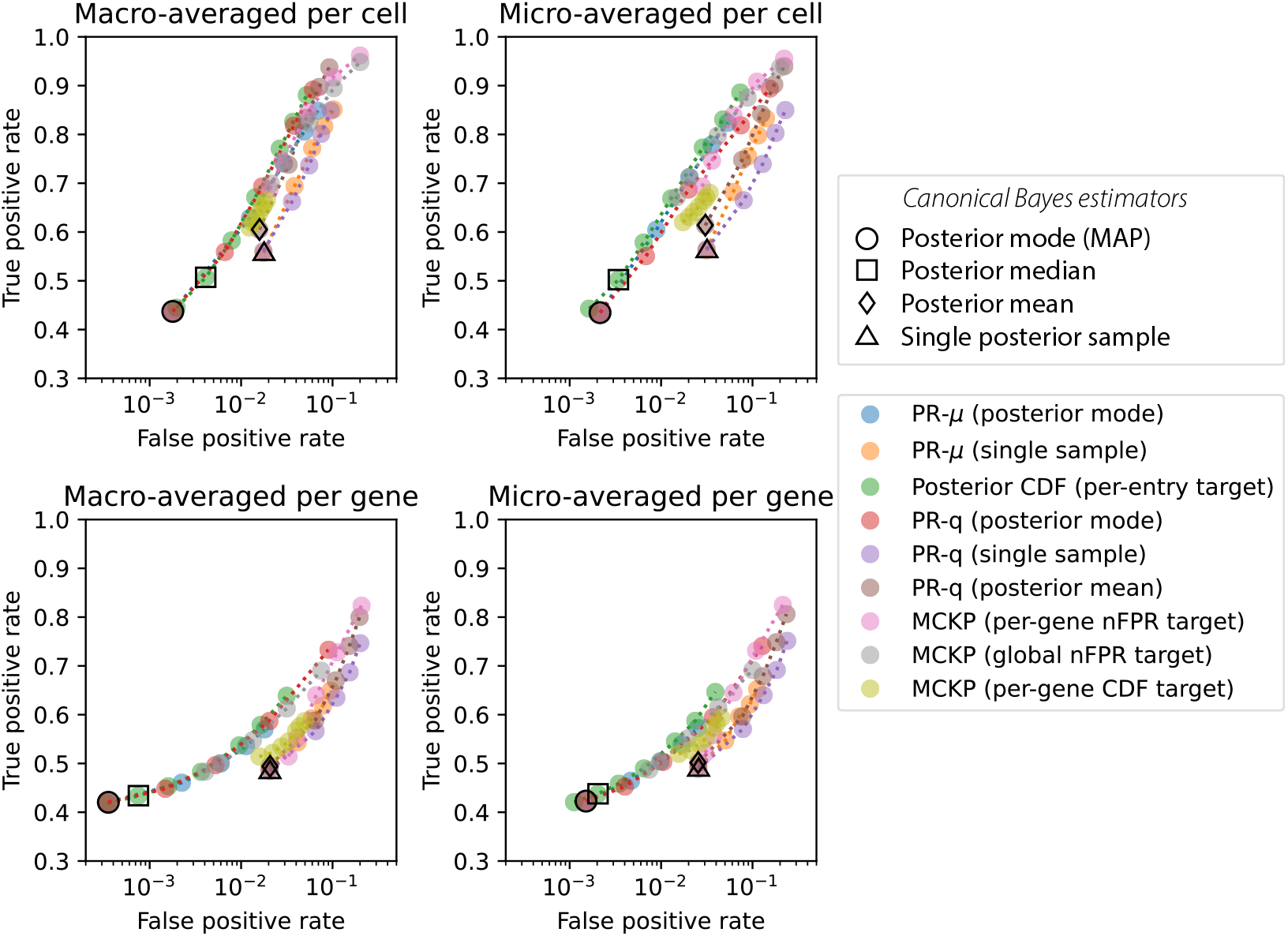
Comparison of output summarization methods from Sections S.1.5 (legend label MCKP), S.1.6 (legend label Posteior CDF), S.1.7 (legend labels PR-*μ* and PR-q). The four panels show four different ways to compute TPR and FPR to display a ROC curve. “Macro-averaged per cell” computes TPR as 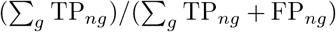, while “micro-averaged per cell” computes TPR as 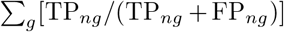. For the “per gene” cases, the sum over genes is replaced by a sum over cells. We exclude genes whose raw data counts are less than 10 summed over all cells. The dots shown represent the mean over all cells or genes as appropriate.

Here, our focus is to evaluate various noise estimation strategies after model fitting and inference. In order to sidestep confounding factors such as our ability to fit the model and infer the noise posterior (which depends on the dataset size, the degree of model faithfulness, and our variational approximations), we assume perfect knowledge of all latent variables other than 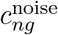. Such an oracle short-circuits the marginalization over *Z* in Eq. (11) and evaluates the integrand at the true value of *Z*. Therefore, the performance metrics given in this section are theoretical upper bounds. A comparison of such theoretical upper bounds with actually attainable end-to-end results is given in Fig. 4c-g.

**Figure S 3:**
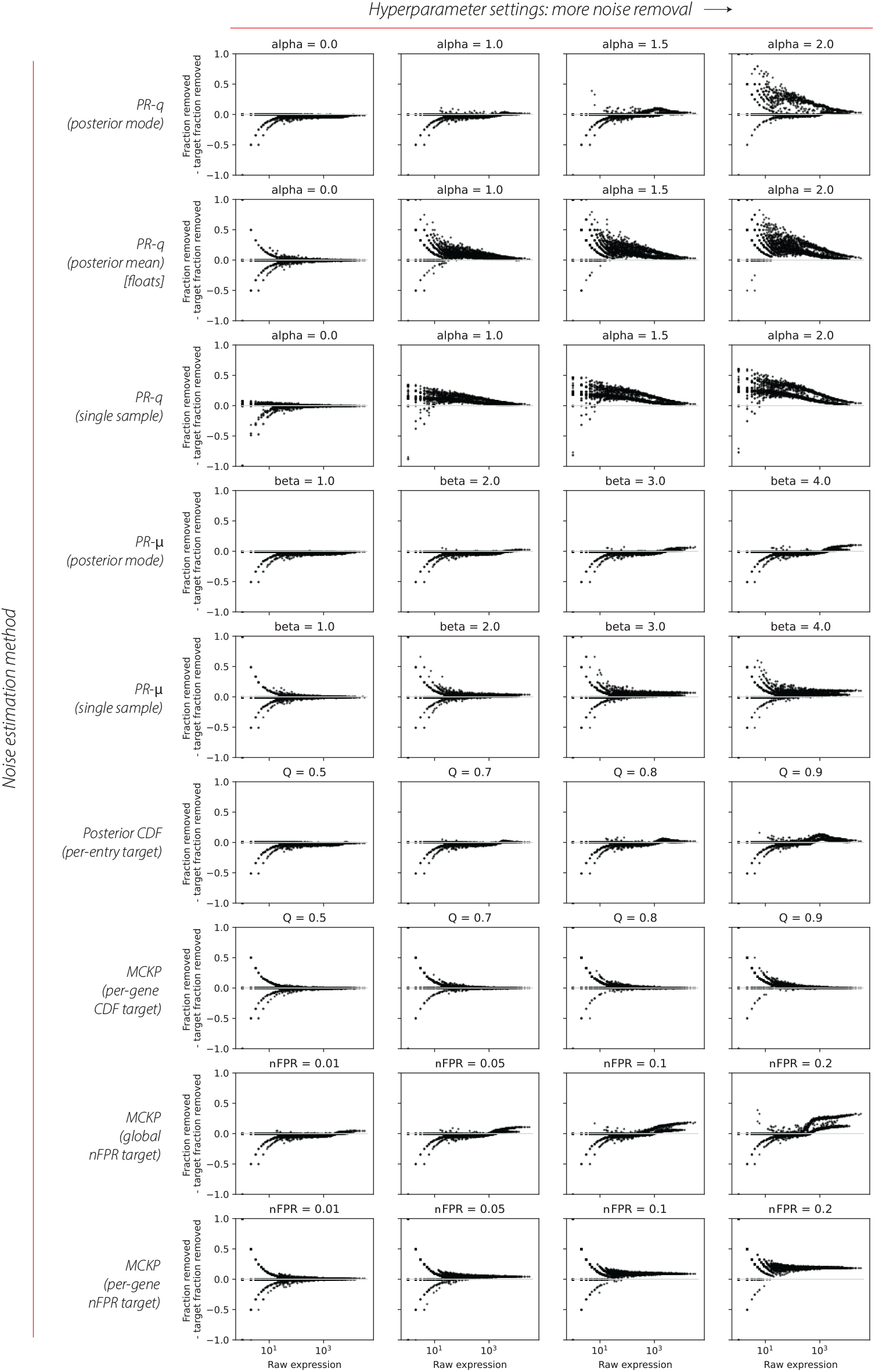
Comparison of different noise estimation methods from Sections S.1.5 (MCKP), S.1.6 (Posterior CDF), and S.1.7 (PR-*μ* and PR-q). Each plot shows the over-removal of each gene (fraction removed - fraction that should have been removed according to truth) for the given method with the hyperparameter setting specified in the title. Each dot is a gene. Positive values indicate that too many counts of the gene were removed at the level of the entire experiment. Row 1 column 1 shows the posterior mode, row 2 column 1 shows the posterior mean, and row 3 column 1 shows a single sample from the unregularized posterior (*α* = 0).

First, we evaluate the different estimators by studying their receiver operating characteristic (ROC) curves. To construct a ROC curve, we consider each *n* × *g* entry of the noise count matrix, take “noise” as the “positive” class, and calculate the 2 × 2 confusion matrix as follows:

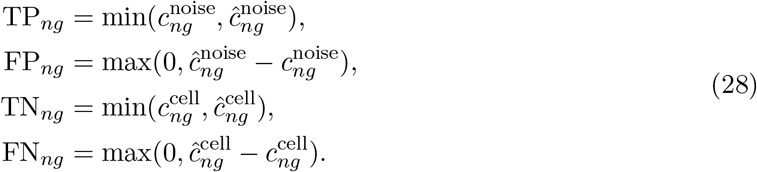

where 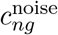 and 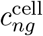 represent the simulated truth values, 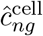 is the `CellBender` output, and 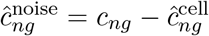. We “summarize” the resulting *n* × *g* confusion matrix either (1) as a “macroaverage” per gene or per cell, where we sum the element-wise 2 × 2 confusion matrices along *n* or *g*, respectively; or (2) as a “micro-average” per gene or or per cell, where we calculate the element-wise TPR_*ng*_ and FPR_*ng*_, remove the undetermined entries, and calculate the arithmetic mean along *n* or *g*, respectively. Fig. S2 shows the resulting ROC curves for various estimation methods. We have further reduced the obtained TPR and FPR values for per-cell (or per-gene) micro- and macro-averages to a single point via arithmetic averaging for better visibility. The canonical Bayes estimators (black circle, square, diamond) each provide a single point on the ROC plane. In contrast, each of our estimators provides a natural parameter for controlling the position on the ROC curve.

It is clear that drawing a random sample either from the actual posterior (black triangle) or from the regularized posterior (PR-*μ*, orange; PR-q, purple), is a poor strategy, while also being inconsistent and non-deterministic estimators. Posterior mean estimators, either unregularized (hexagon) or regularized (PR-q, brown circles), neither produce an integer count matrix, nor are among the top-performing estimators in terms of ROC curve. Estimators based on the regularized posterior mode (PR-*μ*, blue circles; PR-q, red circles), the element-wise posterior CDF quantiles (green circles), and MCKP estimators (per-gene nFPR target, pink; global nFPR target, gray), all do well and are practically tied in terms of the ROC curve, with the estimator based on elementwise posterior CDF quantiles showing a slight advantage in this benchmark.

To further distinguish the characteristics of the different estimators, we also study the over- or under-removal of noise counts for each gene vs. total gene expression in Fig. 3. The ideal estimator is expected (1) to exhibit the same characteristics across the entire gene expression spectrum, and (2) to not under- or over-remove noise counts when the total noise budget is chosen in a balanced way (i.e. *q* = 0.5 for CDF-based targets, or nFPR ≈ 0). Among the top-performing estimators in terms of the ROC analysis, we find that MCKP with a per-gene nFPR target satisfies both expectations (see the last row in Fig. 3). Specifying a dataset-level (global) noise budget tends to over-correct highly-expressed genes (see PR-*μ* posterior mode, and MKCP global nFPR target in Fig. 3).

In summary, our analysis highlights two estimation strategies: (1) the MCKP estimator with gene-wise nFPR control, which shows decent ROC characteristics and a consistent performance across the entire gene expression spectrum; (2) element-wise posterior CDF quantiles, which shows the best ROC characteristics although with some dependence on the gene expression rate. We have chosen the former as the default estimation strategy in the latest release of `CellBender` (v0.3.0). The previous version (v0.2.0) used the PR-*μ* strategy, which as we have shown here, is inferior to MCKP. Finally, we note that all of these estimation strategies are implemented in `CellBender`, should a user have a use case that warrants a strategy other than the default.

#### S.1.9 A fast and exact MCKP solver for strictly log-concave posterior distributions

MCKP is an NP-hard problem which admits a pseudo-polynomial dynamic programming solution. Here, we show that assuming strict log-concavity of the noise posterior distribution leads to a fast and exact solution of MCKP with time complexity 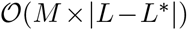, where 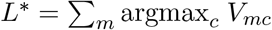.

##### Definition S.1

(Log-concave discrete distribution). A discrete probability distribution 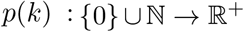 is called log-concave if and only if log *p*(*k* +1) +log *p*(*k* – 1) ≤ 2 log *p*(*k*). It is called strictly log-concave if ≤ is replaced with strict inequality.

Many common probability distributions are log-concave, including Poisson and negative binomial, most of which are also *strictly* log-concave except for a measure zero set of parameters. We do not aim to rigorously prove the conditions for strict log-concavity of our noise posterior distribution. However, we have empirically verified that this property holds in various datasets. To motivate the this empirical observation, consider the limit Φ → 0 and *q*(*Z*) → *δ*(*Z* – *Z*\*). It is easily shown that the the noise posterior tends to the Binomial distribution with a success probability of 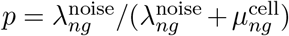 and total number of trials *N* = *c_ng_* in this limit (see Sec. S.1.10), which is a log-concave distribution. Continuity implies the existence of an extended parameter regime around this limit where log-concavity holds. Increasing Φ or the dispersion in *q*(*Z*) can be thought of as imparting uncertainty on *p*. Modeling this uncertainty as a Beta distribution, the noise posterior may then be approximated as a Beta-Binomial distribution, which is also strictly log-concave except for a measure zero set of parameters or irrelevant parameter regimes, e.g. bimodal success probability *p*. Hereafter, we assume the strict log-concavity of the noise posterior as given.

We call the MCKP problem posed by Eq. 12 a *strictly convex MCKP problem* if and only if the reward weights 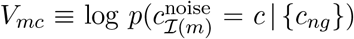 are derived from strictly log-concave distributions. We will show that the strictly convex MCKP problem admits an exact greedy solution. To set the stage, consider the unconstrained MAP estimate 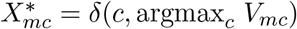 and observe that it achieves the total noise target 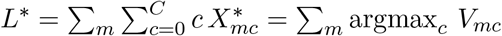. Clearly, if the specified total noise target *L* coincides with *L*\*, then 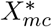 is indeed the optimal solution since each reward term is individually maximized, the constraint is satisfied with equality, and moving away from the equality constraint satisfaction implies deviating from the MAP point and thus decreasing the reward. In a nutshell, our greedy strategy is to take 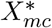 as a reference point and iteratively modify it via best local moves such that the specified noise target is met. To this end, we define Δ = *L* – *L*\* as the gap between the total noise count of the MAP solution 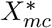 and the specified total noise target. We refer to the sought-after solution as 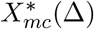. By definition, 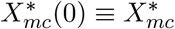. We only consider the case Δ > 0 here. The case Δ < 0 can be worked out by symmetry. Our greedy algorithm for solving this problem for Δ > 0 is as follows. To obtain 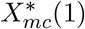 from 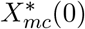, we consider 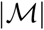 local moves where the noise count for each coordinate *m* is increased by 1 while keeping the other coordinates fixed; we chose the local move that yields the highest possible reward. Note that we are not considering all possible moves that satisfy the constraint, e.g. removing two noise counts from a coordinate and adding three counts to another. We proceed with this greedy strategy in an iterative fashion until we reach the desired Δ.

##### Theorem S.1.

The greedy iterative coordinate ascent algorithm solves the strictly convex MCKP problem exactly.

*Proof*. Consider the following objective function:

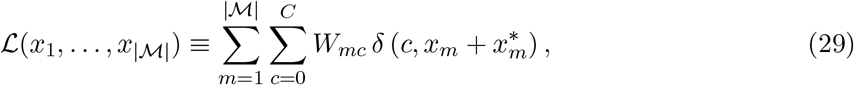

where:

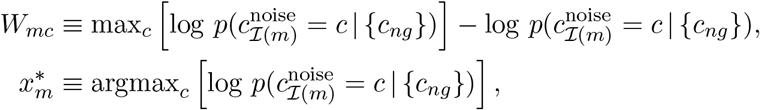

and 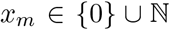 is the “extra” noise counts allocated to count matrix entry *m* on the top of the MAP point 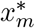. We refer to the vector of extra noise counts and MAP counts as **x** and **x**\*, respectively. Minimizing 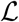 subject to the total noise constraints given in Eq. (12) is equivalent to solving the MCKP problem. In the new notation, 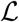 conveniently achieves its minimum value of 0 at **x** = **0**, which corresponds to the unconstrained MAP point. This is due to implicitly setting the MAP point as the reference point in the definition of *W_mc_*. The strict log-concavity of noise posterior distributions implies *strict convexity* of *W_mc_* in the following discrete sense:

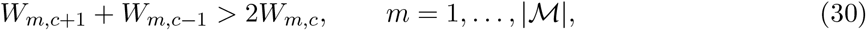

which follows from Definition S.1. As a consequence, 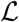 emerges as a separable function of strictly convex one-dimensional functions over non-negative integers. We will use this property repeatedly to establish the optimality of coordinate ascent moves. We define *B*(Δ) as the subspace of points that satisfy the total noise constraint with equality at *L*\* + Δ:

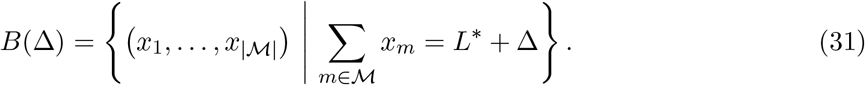

We observe that *B*(Δ) is a discrete convex set in the sense that if 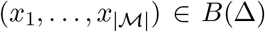, then 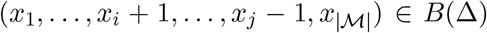 for all *i* and *j*. As a consequence, the restriction of 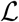 to *B*(Δ) is also strictly convex, implying that: (1) any local minimum of 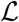 over *B*(Δ) is the global minimum; and (2) the global minimum of 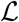 over *B*(Δ) is unique. Therefore, to prove the optimality of coordinate ascent, it is sufficient to show that the point obtained by applying coordinate ascent to the minimizer of 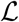 in subspace *B*(Δ), namely **x**\*(Δ), yields a *local* minimum in the next subspace *B*(Δ + 1). Global optimality and uniqueness follows from strict convexity. Consider the set of all 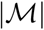 local coordinate ascent moves from **x**\*(Δ), and let 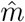 be the coordinate to which adding a noise count accrues the smallest increase in 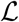. This implies:

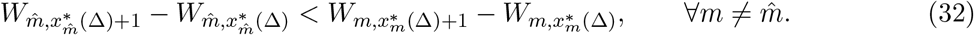

We denote the coordinate ascent update of **x**\*(Δ) as 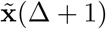:

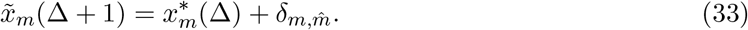

The set of nearest neighbor points of 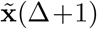, namely 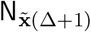, can be written as the union of three mutually exclusive set of points: (1) 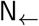 points obtained by moving backward along coordinate 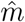, and moving forward along another coordinate 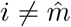, there are 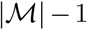 such points; (2) 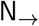 points obtained by moving further forward along coordinate 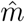, and moving backward along another coordinate 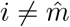; there are 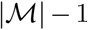 such neighbors; (3) 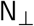 points obtained by keeping coordinate 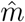 fixed, choosing two other coordinates *i,j* such that 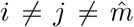, moving forward along *i* and backward along *j*; there are 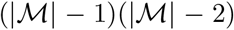 such moves. Put together, the three mutually exclusive sets comprise 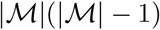 nearest neighbor points of 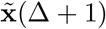,

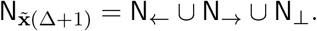

We wish to show that 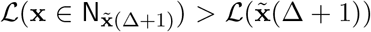. First, we note that the 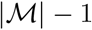 points in 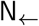 coincide with the 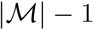 rejected forward moves which by definition lead to a higher value of 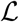 over *B*(Δ + 1), see Eq. (32). Therefore, all points in 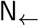 are directions of ascent. For an arbitrary point 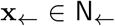 obtained by stepping backward along coordinate *i* and further forward along *m*, we have:

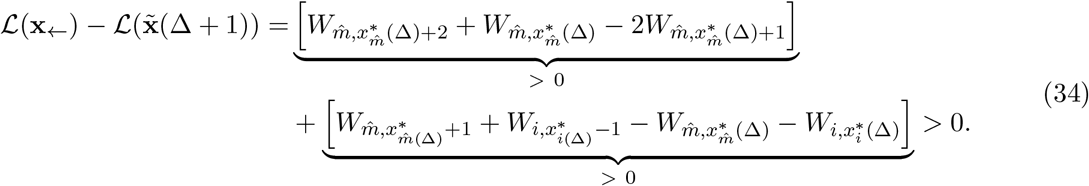

The first term is positive due to strict convexity and the second term is positive due to **x**\*(Δ) being the minimizer of 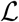 in subspace *B*(Δ). Finally, for a point 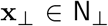 obtained by stepping forward and backward along coordinates *i* and *j*, respectively, we have:

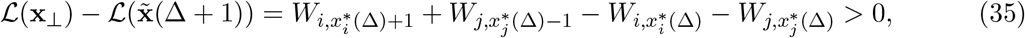

which directly results from **x**\*(Δ) being the minimizer of 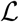 in subspace *B*(Δ). Put together, we have shown that 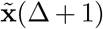 is a *local* minimizer of 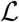 in subspace *B*(Δ + 1). Strict convexity implies that 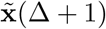 is also the unique and global minimizer:

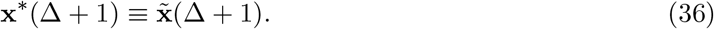

Therefore, following the iterative coordinate ascent trajectory that connects the MAP point **x**\*(0) to **x**\*(Δ) yields the unique solution of the strictly convex MCKP problem. There are Δ = |*L* – *L*\*| iterations and each iteration involves 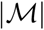 comparisons to locate the optimal coordinate. Therefore, the complexity of this algorithm is 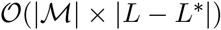. As mentioned earlier, the case Δ < 0 can be worked out by symmetry, i.e. replacing “backward” moves with “forward” moves.

In practice, we implement the coordinate ascent strategy by pre-computing, pooling, and sorting differential coordinate ascents *δ_m,c_* ≡ *W_m,c+1_* – *W_m,c_*. Even though the time complexity of this implementation is 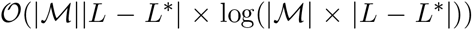, it runs faster on GPU hardware by leveraging parallelism.

#### S.1.10 On the asymptotic bias of canonical Bayes estimators

We mentioned the shortcomings of canonical Bayes estimators as part of our motivations for developing application-specific integer noise estimators. These include the non-integral estimates obtained by posterior mean (PM), and the asymptotic bias of posterior mode estimator, also known as the maximum *a posteriori* (MAP) estimator. In this section, we study these estimators in more detail in a simple setting that is related our application. We consider the simplifying limit Φ → 0 and *q*(*Z*) → *δ*(*Z* – *Z*\*) in Eq. (11), focus on a single count matrix entry, and drop the *n* and *g* indices for brevity. In this limit, the posterior is found to be:

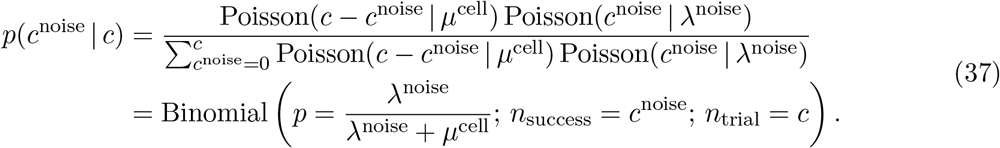

Here, λ^noise^ and *μ*^noise^ correspond to the noise count and cell count rates at the latent variable concentration point *Z*\*. We have also used lim_Φ→0_ NegBinom(*x*|*μ*, Φ) = Poisson(*x*|*μ*). The binomial equivalence can be either derived by interpreting Poisson variables as the sum of Bernoulli variables, or by resorting to the algebraic expression of the Poisson probability mass function. In this limit, we find the PM and MAP estimators to be:

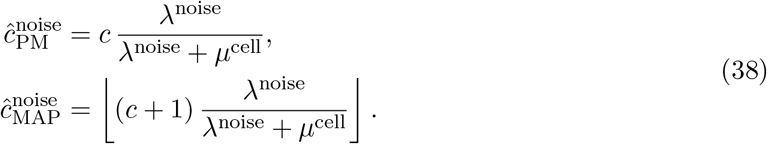

Note that the expression for 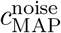 is only valid when the expression appearing in the floor function is non-integer, which is the case except for a measure zero set of points.

We consider *N* i.i.d. realizations of *c*^noise^ and *c*^cell^ and study the asymptotic bias of the two estimators in sample mean. This analysis is an idealization of taking a population of *N* → ∞ droplets containing identical cells, and checking whether or not the empirical mean of a given noise estimator converges to λ^noise^. For the PM estimator, we have:

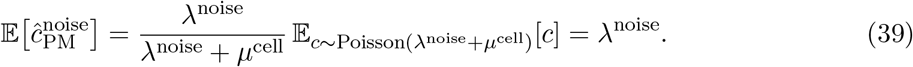

Therefore, we find PM to be consistent and asymptotically unbiased. However, the estimator clearly yields non-integer values, which is undesirable. For the MAP estimator, we have:

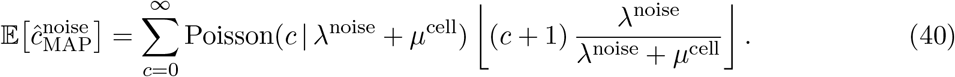

It is easy to see that this estimator is asymptotically biased. The floor term is identically vanishing for 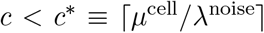. In the relatively low-noise limit λ^noise^ ≪ *μ^cell^*, *c*\* becomes arbitrarily larger than the mode of *c*, which is ≈ *μ*^cell^ in this limit, and subsequently 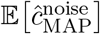 becomes arbitrarily smaller than the expected value of λ^noise^. While the asymptotic bias of the MAP estimator can be studied analytically, we find it more straightforward to resort to a numerical study. We define the relative asymptotic bias of the MAP estimator as:

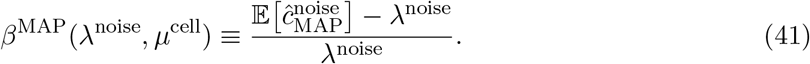

Fig. 6 shows *β*^MAP^ for a range of noise count and cell count prior rates. We notice *β*^MAP^ ≈ – 1 in the regime λ^noise^ ≪ *μ*^cell^, as expected from the pathological behavior of the MAP estimator in the low-noise regime. In this regime, 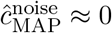, implying that no noise count is removed from any cells.

#### S.1.11 Implementation Details and Technical Remarks

The default architecture for the encoder network NN_*z*_ that maps *c_ng_* to the bundle of **z**-posterior location and scale, (**z**_*n;μ*_, **z**_*n;σ*_), has one hidden layer of 500 units, and the encoded dimension of Z of **z**_*n*_ is 100. Similarly, the decoder network NN_χ_ that maps **z**_*n*_ to χ_*ng*_ has one hidden layer of 500 units, a linear readout, followed by a `softmax` operation to bring the output to the (*G* – 1)-simplex of normalized endogenous feature frequencies. The encoding network for *y_n_*, 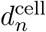, and *ϵ_n_*, denoted by NN_enc_ for brevity, works as follows. Inputs to the network consist of raw counts as well as three additional features which are hand-crafted: (1) the log of total counts per droplet, (2) the log of the number of nonzero genes per droplet, and (3) the overlap with the current estimate of the ambient RNA profile (which is calculated as a log probability that the observed droplet counts were drawn from a Poisson with rate equal to 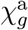). Hand-crafted features are concatenated to counts to form the input to the network. By default, the network has two hidden layers [100, 50]. From the last hidden layer, three separate linear transformations take the hidden state and produce (1) logit cell probability logit *q_n_*, (2) the inverse variance of the gamma distribution for *ϵ_n_*, and (3) the log of mean cell sizes 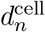. Weights are initialized using PyTorch defaults, except for the weights which connect the handcrafted log-counts per droplet input feature to the output for *q_n_*, which are initialized to 1, so that the network starts with an condition that cell probability should closely follow log counts. Softplus non-linearities are used throughout. In practice, `CellBender` results are not very sensitive to the architecture of the encoders, and network architectures can be changed from the default values using command-line arguments.

We note that initially learned biological gene expression landscape, NN_χ_(**z**_*n*_), may itself be contaminated with background RNA counts. However, as the inference procedure progresses and as the estimate of the background RNA profile improves, the maximum likelihood principle encourages the neural network to correct in a self-consistent fashion and learn to represent background-free gene expression profiles.

For numerical stability and to preclude vanishing gradients, we handle all probabilities in logit-space in our implementation. During training, the log probability of **z**_*n*_ is only added to the ELBO for droplets which have been found to contain cells (that is, for droplets *n* where a sample of *y_n_* is 1). The discrete latent variable *y_n_* cannot be re-parameterized, and so we use full enumeration over cell / no cell (*y_n_* being 1 or 0) in our variational posterior to reduce variance. This is achieved using the `TraceEnum_ELBO` SVI objective available in Pyro. Integration over the continuous latent variables appearing in the ELBO (Eq. 4) is done using a single Monte Carlo sample.

Training happens in random mini-batches. Each full epoch trains on a fixed subset of barcodes from the dataset as well as a randomly-sampled subset of empty droplet barcodes that changes each epoch. This is done in order to cover the tens of thousands of empty droplets without taking excessive computation time. The fraction of each minibatch that is composed of these randomly-sampled empty droplets can be specified using a command line argument (by default, we use 20 percent).

The training loop converges typically within about 150 epochs. For a typical 10x scRNA-seq experiment containing 5-30 thousand cells, the total runtime of the tool ranges around 20 min - 1 hour using an NVIDIA Tesla T4 or K80 GPU, depending on the size of the dataset and chosen parameters. The stochastic optimizer used is a version of the Adam optimizer with gradient clipping. A OneCycle learning rate scheduler is used by default. Optimization proceeds for a pre-defined number of epochs, which can be set via command line arguments. Default is 150 epochs, and the OneCycle scheduler increases the learning rate to 10 times the user-defined `--learning-rate` at maximum. The default learning rate is 1e–4.

The tool saves checkpoints at user-defined intervals, which can be used to resume training or create a new output with a different false positive rate. Checkpoints enable the use of cheaper preemptible cloud machines via the Terra platform (app.terra.bio). More generally, any workflow deployment using Cromwell (https://github.com/broadinstitute/cromwell) version 55+ can automatically benefit from this checkpointing functionality, so that a preempted workflow can pick up where it left off instead of starting from scratch.

#### S.1.12 Single-cell analysis workflow and cell quality control details

Analysis workflows for single-cell data were carried out in scanpy [17] version 1.9.1. We employed a rudimentary cell quality control (QC) post-`CellBender`, i.e. removing cells using percentile-based thresholds on UMI count, gene count, and mitochondrial read fraction. UMAPs were created after (1) finding highly variable genes using the `seurat_v3` algorithm implemented in scanpy, (2) normalizing counts per cell, (3) log-scaling counts, (4) scaling counts of 2000 highly-variable genes, and (5) performing PCA on those scaled values for the highly-variable genes. A nearest neighbor graph was constructed with 20 neighbors based on cosine distance in PC space (top 25 PCs). Clustering was performed using the Leiden algorithm at the same resolution for both raw and post-`CellBender` data. Dataset-specific cell QC thresholds and the statistics of initial and final cell calls are as follows:

##### pbmc8k scRNA-seq dataset

We remove top 5% of high UMI count droplets and top 5% of high unique gene count droplets (to eliminate doublets), as well as top 10% of high mitochondrial read fraction droplets, and no lower cutoff for total number of genes per droplet. This left 7515 cells remaining out of an initial 8903 droplets.

##### rat6k snRNA-seq dataset

We remove top 15% of high UMI count droplets and top 15% of high unique gene count droplets (to eliminate doublets), as well as top 10% of high mitochondrial read fraction droplets, and eliminate droplets with fewer than 100 genes. This left 5868 cells remaining out of an initial 10445 droplets.

##### pbmc5k CITE-seq dataset

We remove top 5% of high UMI count droplets and top 5% of high unique gene count droplets (to eliminate doublets), as well as top 10% of high mitochondrial read fraction droplets, and a lower cutoff of 300 genes per droplet. This left 4451 cells remaining out of an initial 5754 droplets.

##### hgmm12k scRNA-seq dataset

No cell QC was performed before creating the `hgmm12k` results plots: all the `CellBender` “non-empty” droplets are included.

#### S.1.13 pbmc5k CITE-seq dataset quality control and normalization

For the plot in Fig. 5e, the following antibody features were omitted due to low correlation between antibody counts and mRNA counts per cluster in the raw data: `CD34_TotalSeqB` (also has very low mRNA counts), `CD45RA_TotalSeqB` and `CD45RO_TotalSeqB` (where the poor correlation to *PTPRC* mRNA counts which is understood given the high splicing specificity of *PTPRPC* in different immune subtypes which make the expectation of having a linear correlation meaningless in principle), `CD69_TotalSeqB`, `CD137_TotalSeqB`, `CD197_TotalSeqB`, `CD274_TotalSeqB`, `IgG1_control_TotalSeqB`, `IgG2a_control_TotalSeqB`, and `IgG2b_control_TotalSeqB`. Low correlation was defined as a slope of less than 1 for a fit using weighted ordinary least squares when plotting log1p antibody counts versus log1p mRNA counts. The following features were omitted due to low mRNA counts in the raw data: `CD15_TotalSeqB`, `CD25_TotalSeqB`, `CD278_TotalSeqB`, and `PD-1_TotalSeqB`. Low mRNA counts was defined as the maximum mean-expression value over all clusters being ≤ 0.2 counts. Leaving out these features was for clarity of presentation (the scaling transformation, below, does not work well for those outliers), but the excluded features are all plotted in Supplementary Fig. S16a-b.

The scaling transformation used to plot data in Fig. 5e, by collapsing all data onto a single line, is as follows. The raw RNA expression data is *x*, while the raw antibody data is *y*:

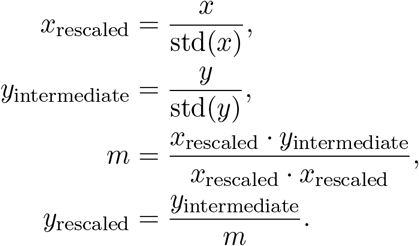

#### S.1.14 Simulated data generation

Data were simulated according to a model which was slightly and intentionally mis-specified for `CellBender`’s model, in that each cell within a cell type *k* is not given the exact same underlying expression profile 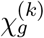, but instead each cell has its expression profile drawn from a Dirichlet distribution with a common set of concentration parameters for each cell type, 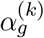. Thus the data will be a bit overdispersed compared to `CellBender`’s model. Details of the simulations are included in a notebook for code reproducibility, and the data simulation function is included as part of the `CellBender` package.

The simulator first samples the base gene expression profiles for *k* cell types, 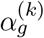, from flat Dirichlet distributions, e.g. 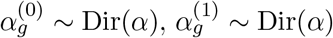, etc. These *k* cell-type expression profiles are then optionally made to be artificially similar to 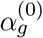 via a parameter *η* by applying the transformation 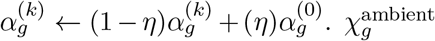 is set to the (normalized) average of the 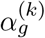, weighted by the number of simulated cells of each type and the average UMI per cell type. Then, for a given cell type *k* with *n* cells, the simulation proceeds as:

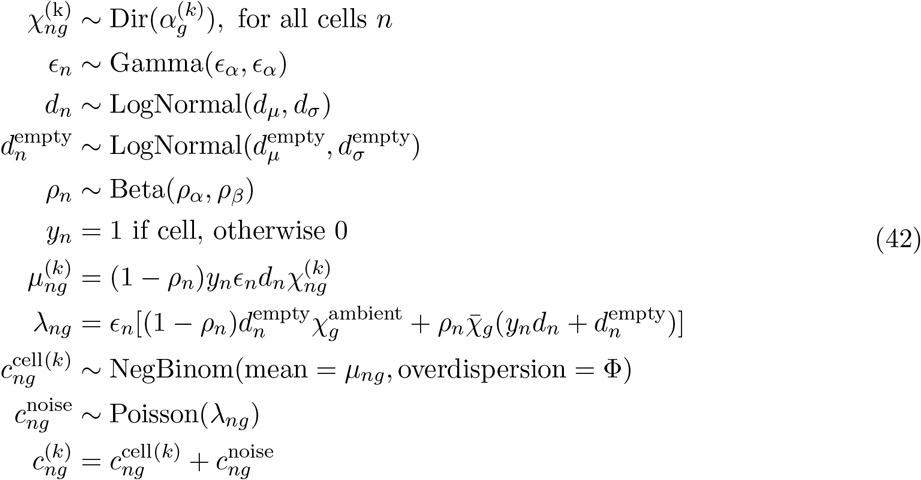

where the simulated counts for cell type *k* are 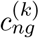, and 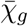 is the same as 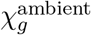 in these simulations, and all the other variables not specified above are hyperparamter inputs to the simulation. Cells are simulated with *y_n_* = 1 and empty droplets are obtained by setting *y_n_* = 0. Cell counts are simulated, one cell type *k* at a time, followed by empty droplets, in order to obtain a full dataset.

### S.2 The phenomenology of systematic background noise counts in droplet-based single-cell assays

In this section, we review the phenomenology of systematic background noise counts by examining four exhibits in different experiments. Next, we review a number of mechanisms that satisfactorily explain all aspects of the phenomenology. Some of these mechanisms have been noted by other authors, though we provide them in one place for completeness.

#### Exhibit 1

Examining the counts of total unique UMIs per droplet in a typical 10x scRNA-seq experiment reveals that there are thousands of high-count droplets followed by a much larger number of low-count droplets (See Supplementary Fig. S4a,e and note the logarithmic scale of the axes). Here, the word “counts” is used as shorthand for counts of unique UMIs summed over all genes. The number of high-count droplets typically agree in order of magnitude with the expected number of cells given the protocol [2]. The low-count droplets typically have tens to hundreds of UMIs each (i.e. far fewer counts than high-count droplets), and significantly outnumber the expected number of cells. Therefore, these droplets are unlikely to have their counts originating from a physically encapsulated cell.

#### Exhibit 2

Experiments with mixtures of different cell types have shown that some of the transcripts in each droplet do not originate from the cell encapsulated within the droplet. That is, even for droplets that do contain cells, there is still some exogenous background noise in the count matrix. Fig. 4a shows a scRNA-seq dataset generated using a mixture of human and mouse cells from 10x Genomics. It is noticed that for a few percent of the count data, human transcripts are assigned to a droplet where the vast majority of transcripts are mouse, and vice-versa. This mixing can happen when a human cell and a mouse cell are captured in the same droplet, but these “doublets” can be easily identified due to the fact that they have tens of thousands of counts from each species (doublets are excluded from Fig. 4a). Even droplets that do not contain doublets still have nonzero counts from transcripts of the other species (see the inset of Fig. 1c, for example). Droplets with tens of thousands of human counts typically have a few hundred mouse counts, and vice-versa.

#### Exhibit 3

The phenomenon of non-zero RNA counts in empty droplets is even seen in experiments where the preparation is entirely devoid of cells but rather contains a high concentration of spike-ins (e.g. see the publicly available `ercc` dataset from 10x Genomics [2]). In this experiment, approximately 1000 droplets were prepared from the same spike-in soup and used for library production. Quite curiously, the total counts vs. sorted barcode plot looks similar to experiments that include cells: a first region including approximately 1000 high-count droplets, followed by thousands of droplets with approximately 100 UMIs each. The appearance of the second region resembling “empty droplets” is unexpected, since all droplets are filled uniformly with the same amount of spike-in transcripts.

#### Exhibit 4

The phenomenon of increased exonic read fraction among empty droplets in snRNA-seq data. When cells are lysed to liberate nuclei, lots of cytoplasmic RNAs are also liberated, becoming cell-free ambient RNA. Despite washing steps, it is still clear that these cytoplasmic RNAs, enriched for higher exon fraction, are present at higher proportions in empty droplets. See Fig. 1d for an example demonstrating this for a snRNA-seq dataset.

Based on these four exhibits, we review and identify several distinct mechanisms that explain the full phenomenology of background RNA in droplet-based scRNA-seq experiments. In what follows, we assume an mRNA capture assay for concreteness, however, as mentioned earlier, the same phenomenology identically applies to all other barcoded molecular features, including protein (CITE-seq), chromatic accessibility (scATAC-seq), and Perturb-seq sgRNA guides, as should become clear from the discussion.

#### Sequencing or synthesis errors in the droplet barcode

The presence of uncorrected sequencing errors in droplet barcodes or impurity of synthesized barcodes on capture beads will result in a spreading of transcripts across droplets. In particular, one expects a net flow of transcripts from RNA-rich (cell-containing) droplets to otherwise RNA-free (cell-free) droplets. Quantitative estimates of barcode sequencing error indicate that far more empty droplets are observed than can be explained by sequencing error alone. The effective provisions for barcode error-correction employed by the 10x Genomics scRNA-seq protocol (using a whitelist, no homo-polymers, and a Hamming distance ≥ 2 between droplet barcodes) allows most barcode sequencing errors to be corrected. In our simulations using typical base substitution and insertion/deletion error rates, we found that at most 2 percent of erroneous droplet barcodes were corrected to the wrong barcode. Given that a typical 10x v2 scRNA-seq experiment yields less than 5 percent invalid barcodes, we estimate that at most 1 in 1000 transcripts would be mis-assigned due to wrong barcode error correction. This rate is 3 orders of magnitude lower than what is required to produce non-zero transcript counts in empty droplets as seen in typical experiments. The presence of error or impurity in barcode synthesis, however, might still explain part of the background RNA phenomenology. Unfortunately, details of the 10x barcode synthesis protocol are not public.

#### Presence of ambient molecules in the cell suspension

Cell-free “ambient” RNA that is physically present in the cell suspension and is encapsulated in a droplet will clearly contribute to the background while generating non-zero transcript counts in otherwise empty droplets. This mechanism is shown schematically in Fig. 1b. Cell-free RNA is present in the aqueous cell suspension, either as a result of normal biological processes or as a result of tissue dissociation, cell death, or other stresses experienced by cells during the isolation protocol which may cause cells to die or lyse (see Fig. 1a). Such a mechanism has been proposed by others as well [1, 11, 32, 42].

#### Barcode swapping and PCR chimera formation

Swapping of droplet barcode between transcripts during mixed-template PCR amplification via formation of heteroduplex/chimeric molecules [12–14], and/or on the flowcell during sequencing [58], will spread transcripts across droplets and generate a background. Chimeric fragments incorporate mRNA sequences from one original molecule and a droplet barcode (and UMI) either from a different original molecule or from a previously unused barcoded capture oligo. In the 10x Genomics protocol, there is a large amount of sequence complementarity, both in the Illumina primers as well as in the poly(T) region (the means by which these molecules were captured in the first place). As PCR progresses through many rounds, primers are depleted. Eventually, extension could be primed by (1) incomplete extension products from other molecules, as suggested by Dixit [13], or by (2) unused and inadequately washed capture oligos that were used to capture poly(A)-tailed mRNAs at the outset. These mechanisms would both result in transcripts which are assigned to the wrong droplets. This process of chimera formation is prone to occur in all mixed-template PCR reactions, and is not unique to scRNA-seq library preparation protocols [14]. The hallmark of PCR chimeras is a relatively lower read per UMI (“family size”) compared to non-chimeric and properly amplified cDNA molecules. It has been shown that identifying and removing library fragments with small read per UMI (e.g. 1) is highly effective in removing off-target cross-species counts in species-mixing experiments [13].

#### Cross-contamination of capture oligo beads on the microfluidic device

The capture oligo gel beads (referred to as GEMs in the 10x Genomics scRNA-seq protocol) flow in a microfluidic channel (see Fig. 1b, green hexagons). The GEMs are tightly packed in the channel to achieve a precise flow control that allow their super-Poisson loading into droplets [2]. Since these gel beads are soluble in certain conditions in aqueous solution, it is reasonable to expect that some small number of capture oligos could be released from the GEM in the channel, leading to cross-contamination due to “ambient” capture oligos from other GEMs. Therefore, even if the GEMs were synthesized with high barcode purity to begin with, there could be some mixing in the microfluidic device. The downstream effect is similar to GEM impurity or barcode error, and produces a background. The appearance of thousands of low-count droplets in the spike-in experiment (cf. Exhibit 3 above) is likely to be associated with this mechanism.

We may summarize the above mechanisms in two main categories:

#### Physically encapsulated exogenous molecules

The mRNAs were physically present in the droplet at the time the droplet was formed. This is the “soup” or cell-free ambient RNA hypothesis (depicted in Fig. 1a-b). A small amount of cell-free ambient RNA was present in solution (due to cell death, lysis, etc.) at the time the droplets were formed, and some of this ambient RNA was packaged into each droplet, along with cells.

#### Barcode misassignment

The mRNAs were not physically present in the droplet at the time the droplet was formed, but were later assigned to that droplet. This could happen in one of two ways: (1) a molecule’s droplet barcode was physically swapped to a cell-containing droplet barcode at some point in the protocol, (2) a molecule was mis-assigned to a different droplet barcode due to sequencing error or capture oligo impurity or contamination.

These two explanations could lead to different “background RNA” profiles. If cell-free ambient RNA was physically packaged into each droplet, then each droplet should contain a small sample of this *same* RNA profile, which could be related to cell expression or could be slightly different (for example, it could in principle incorporate an exogenous contaminant or a higher proportion of mitrochondrial mRNA if the source of cell-free RNA is related to cell death). If the cause of background RNA is instead barcode swapping, sequencing error, or capture oligo impurity, then it would be expected that the background RNA profile would be *exactly* the average of all the RNA sequenced in the experiment, because these mechanisms act at random.

### S.3 Obtaining and Using CellBender

We have implemented the model and the inference method using the Pyro probabilistic programming language [16] and PyTorch [59] and presented it as a user-friendly, production-grade, and stand-alone command line tool. We internally refer to the background noise removal algorithm implemented in `CellBender` as `remove-background`. In the future, together with the community of `CellBender` users and developers, we hope to extend `CellBender` in exciting new directions and to other high-throughput cell biology assays.

`CellBender` can be obtained from https://github.com/broadinstitute/CellBender. Additional documentation is available at https://cellbender.readthedocs.io. `CellBender` modules are also available as workflows on Terra (app.terra.bio), a secure open platform for collaborative omics analysis, and can be run on the cloud with zero setup.

#### S.3.1 CellBender remove-background inputs

The current version of `CellBender remove-background` (0.3.0) takes the following file formats as input: (1) raw HDF5 file from 10x Genomics’ CellRanger v2+ count pipeline, (2) raw MTX file, with accompanying TSV files, in CellRanger format, (3) raw DropSeq DGE file, (4) H5AD file in AnnData format [17], (5) raw BD Rhapsody CSV file, (6) Loom file readable by AnnData. Ensure that empty droplets are included in the file. The AnnData, DropSeq DGE, and CellRanger MTX formats are particularly general, and data from other sources can be massaged into one of those formats.

#### S.3.2 CellBender remove-background outputs

The output of `CellBender remove-background` provides several useful quantities: (1) inferred background-subtracted count matrix, (2) probability that each droplet contains a cell, (3) lowdimensional latent representation of gene expression for each cell, and (4) the ambient profile, among other latent variables.

There is an input parameter `--fpr` which controls the expected “nominal false positive rate”, where a false positive is a real count that has erroneously been identified as background and removed. Setting nFPR to 0.01 means that the algorithm will remove as much noise as possible while controlling the expected removal of real signal to ≈ 1% above the estimated dataset-wide noise level. It is to be understood that this constraint is enforced in expectation and is approximate: assuming the model fits the data perfectly (no model misspecification), the estimate will be correct. There is an inherent trade-off in noise-reduction where the removal of more noise comes at the expense of removal of more signal. The nFPR parameter allows the user to control that tradeoff. Multiple false posistive rate inputs will result in multiple output count matrices. Since we marginalize over 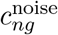 and 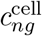 during training, constructing the output 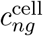 at a given nFPR is a several-step process, and is detailed in Supplemental Sec. S.1.4.

The probability that each droplet contains a cell is given by *q_n_*, the latent variable encoded by NN_*y*_. The low-dimensional latent representation of gene expression is given by the encoded **z**_*n;μ*_ for each cell. And the ambient RNA profile is inferred as 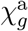. By default, `CellBender remove-background` creates an HTML output report, showing several diagnostics including the progress of the inference procedure and salient changes in the output count matrix, making recommendations and issuing warnings as necessary.

### S.4 Data Availability

The datasets used in this study are the following:

`pbmc8k`: The publicly available PBMC 8k dataset from 10x Genomics called “8k PBMCs from a healthy donor”, run with v2 chemistry and analyzed with CellRanger 2.1.0, available at https://www.10xgenomics.com/resources/datasets/8-k-pbm-cs-from-a-healthy-donor-2-standard-2-1-0.
`heart600k`: The published dataset from the Broad-Bayer Precision Cardiology Lab called “Single-nuclei profiling of human dilated and hypertrophic cardiomyopathy” [23], run with 10x Genomics 3’ capture v3 chemistry and analyzed with CellRanger 4.0.0, available at https://singlecell.broadinstitute.org/single_cell/study/SCP1303/single-nuclei-profiling-of-human-dilated-and-hypertrophic-cardiomyopathy.
`hgmm12k`: The publicly available HGMM 12k dataset from 10x Genomics called “12k 1:1 Mixture of Fresh Frozen Human (HEK293T) and Mouse (NIH3T3) Cells”, run with v2 chemistry and analyzed with CellRanger 2.1.0, available at https://www.10xgenomics.com/resources/datasets/12-k-1-1-mixture-of-fresh-frozen-human-hek-293-t-and-mouse-nih-3-t-3-cells-2-standard-2-1-0.
`pbmc5k`: The publicly available PBMC 5k dataset with antibodies from 10x Genomics called “5k Peripheral Blood Mononuclear Cells (PBMCs) from a Healthy Donor with a Panel of TotalSeq™-B Antibodies (Next GEM)”, run with v3 Next GEM chemistry and analyzed with CellRanger 3.1.0, available at https://www.10xgenomics.com/resources/datasets/5-k-peripheral-blood-mononuclear-cells-pbm-cs-from-a-healthy-donor-with-cell-surface-proteins-next-gem-3-1-standard-3-1-0.
`rat6k`: A snRNA-seq dataset from healthy Wistar rat left atrium, comprising approximately 6000 nuclei, processed on 10x Genomics platform using v2 chemistry and analyzed with CellRanger 3.1.0. Dataset provided by Patrick Ellinor’s group at the Broad Institute as part of the Broad-Bayer Precision Cardiology Lab. Experiment performed by authors Alessandro Arduini and Amer-Denis Akkad.. The dataset will be made publicly available upon publication.

### S.5 Software packages used

Details of all software packages used are included in Supplementary Table S1.

**Table S 1:** Details of software packages used.

| Package | Version | Reference | URL |
| --- | --- | --- | --- |
| CellBender | 0.3.0 | (this work) | https://github.com/broadinstitute/CellBender |
| Pyro | 1.8.1 | Ref. 16 | https://github.com/pyro-ppl/pyro |
| PyTorch | 1.11.0 | Ref. 59 | https://github.com/pytorch/pytorch |
| Scanpy | 1.9.1 | Ref. 17 | https://github.com/scverse/scanpy |
| Harmony-pytorch | 0.1.7 |  | https://github.com/lilab-bcb/harmony-pytorch |
| Limma | 3.48.3 | Ref. 60 | https://rdrr.io/bioc/limma |
| Voom | (Limma) | Ref. 55 | https://rdrr.io/bioc/limma/man/voom.html |

### S.6 Supplementary Results

A set of Jupyter notebooks used to make all plots in this paper is available, showing all details of every analysis.

#### S.6.1 Examining inputs and CellBender outputs

Here we show two datasets: one scRNA-seq, `pbmc8k`, and the other snRNA-seq, `rat6k`. Fig. S 4 panels (a) and (e) show the UMI curves from the raw CellRanger data. Several “regions” of the UMI curves can be identified and have been labeled: (1) cells, in green, (2) the “empty droplet plateau”, in red, and (3) putative barcode errors, in blue/gray. The two datasets share these three regions, but the UMI curves are quite different. In the PBMC dataset, very deep sequencing has led to a large number of UMIs per cell (large for 10x 3’ capture v2 chemistry). The fact that this is whole-cell and that the PBMCs exist as separate cells suspended in blood makes it relatively easier to isolate these cells and capture them, whole, in droplets. This leads to a sharp distinction between cells and empty droplets. Fig. S4b shows that, between droplet 8000 and 10,000 on a rank-ordered plot, the UMI count (rather) sharply transitions from cells, with around 4000 UMI counts, to empty droplets, with around 100 UMI counts. The “empty droplet plateau” in the UMI curve in panel (a) can be seen to extend several tens of thousands of droplets, to near droplet 80,000. This number reflects the total number of whitelisted 10x barcodes that were physically present in the library. Beyond this plateau region is a long stretch of possibly several hundred thousand more barcodes, most with fewer than 10 UMI counts, and the vast majority with only 1 or 2 counts. This region, labeled “putative barcode errors”, is hypothesized to contain a mixture of barcode sequencing errors (which were unable to be corrected to their true whitelist barcode) as well as potential impurities in barcode / bead manufacture.

The snRNA-seq dataset, on the other hand, is much more challenging in terms of calling cells. Fig. S4 panels (e) and (f) show that there is a large region of several thousand droplets, from rank-ordered droplet 8000 to 15000, where the UMI count is quite low, and very similar to the UMI count in empty droplets. For many of these droplets, it is unclear if a nucleus is present at all. These particular nuclei were isolated from Wistar rat heart tissue. During tissue dissociation and cell lysis (which takes place in order to obtain nuclei), a large amount of cytoplasmic material is liberated and becomes free in solution. Nuclei also have fewer starting mRNA molecules than whole cells. All of these experimental details commonly result in snRNA-seq experiments having the median UMI count per nucleus (here 472) much closer to the median UMI count per empty droplet (here 78). Calling cells (nuclei) is much more difficult, and the `CellBender` result is shown in Fig. S4f. A comparison with other cell calling algorithms is shown in Fig. 3.

After cell quality control steps (see S.1.12), we plot the top two principal components of the `CellBender` latent space in Fig S4c,g. The labels obtained from the traditional scanpy workflow (S.1.12) are shown as colors, and happen to agree quite well with the groupings of cells in the `CellBender` latent space. This provides some evidence that `CellBender` is able to learn a prior that encodes “cell type”.

Comparing panel (h) with (d) shows that a larger fraction of counts were removed from many genes in the snRNA-seq dataset (h). The higher ratio of noise to signal in the nuclei dataset is responsible for this greater removal of counts. These plots, as well as several others, are included in an HTML output report produced by `CellBender`.

#### S.6.2 Downstream differential expression in a simulated sample cohort

One major use case for `CellBender` is to remove noise from datasets in such a way that several datasets can be fairly compared afterward using a differnetial expression analysis. This is especially important for large-scale atlas-building efforts, and case-control studies where several samples have been measured. In a cohort setting, it is important to set nFPR = 0 to avoid over-correction beyond the expected noise budget. Using larger values of nFPR naturally imparts a bias on the output by preferentially keeping only the most certain cell counts, which is unsuitable when aggregating data from many samples.

In order to show the effect of `CellBender` on such downstream differential expression testing in a cohort setting, we have constructed a simulated cohort of six samples (A1, A2, A3, B1, B2, B3), split into two batches of three samples each. Data are simulated as in S.1.14, using the full noise model with a mean of 500 UMI counts per empty droplet in all simulated datasets. Each simulated dataset is composed of two cell types, whose expression profiles are taken to be the average expression profiles of the cardiomyocytes and fibroblasts in the `rat6k` dataset. This is done for the purpose of being able to examine a dotplot with real gene names. Simulated cardiomyocytes in these simulations have a mean UMI count of 10000, while fibroblasts have a mean UMI count of 5000. The only difference between samples from batch A and batch B is that samples from batch A have 1000 cardiomyocytes, while samples from batch B have 4000 cardiomyocytes. All samples have 2000 fibroblasts, and the noise profile in all samples is taken to be the average expression over the entire experiment, which only differs between batches due to the different number of cells present. The three samples in each group (A or B) are not simulated using identical parameters. Instead, a “batch effect” between samples labeled (1, 2, 3) is simulated by raising the expression profiles to the power (0.75, 1.0, 1.25) respectively, and normalizing. The batch effect generated this way is quite large, and harmony-pytorch is used to correct for this batch effect and align the cell types across samples.

The simulated cohort is constructed this way in order to have a clear expected outcome: that when we perform a differential expression test between cardiomyocytes from batch A and batch B, there should be no differentially-expressed genes. The same for fibroblasts when testing is performed between batch A and batch B. The samples are simulated so that the true expression profiles of these cell types are the same in both batches. Any differentially-expressed genes that show up in the raw data are due to background noise. Given that simulated cardiomyocytes have 2x more UMIs and that the background noise is taken as the average expression over the entire experiment, we expect to see cardiomyocyte genes (which vary in the background noise across batches due to the different numbers of cardiomyocytes) as spurious differentially-expressed genes in fibroblasts. We also expect this spurious differential expression to disappear with `CellBender` pre-processing.

The simulated cohort is shown in Fig. S5a. As expected, when fibroblasts are tested for differentially expressed genes between batches A and B, the raw data shows that many genes come up significant (Fig. S5d). However, after `CellBender`, these spurious results have been eliminated and no new spurious results have been introduced (Fig. S5g, where labeled genes are the same as in panel d). The `CellBender` data shown here used nFPR = 0.01. Note that nFPR settings of 0.02 and higher do result in over-correction, which results in spurious differential expression results that are different from the raw data (while the spurious results from the raw data still disappear). While nFPR 0.01 did not lead to over-correction in this case, we still recommend setting nFPR = 0 in a cohort as a conservative measure. Differential expression testing was performed with limma-voom as in [23], summing over cells in the same cluster and sample to create pseudobulk vectors (as recommended in [61]), fitting the model ‘~ 0+cluster : batch’, and examining contrasts for cluster comparisons across batches.

#### S.6.3 pbmc8k additional information

A violin plot of *LYZ* is shown in Supplementary Fig. S8b for each cluster in the dataset `pbmc8k` before and after `CellBender`, and shows markedly improved cell-type-specificity of *LYZ* expression as compared to the raw data in (a).

UMAP plots of per-cell expression before and after `CellBender` are included in Fig. S9 for four genes which were highly expressed and which were targeted by `CellBender` due to high ambient expression. Comparison of the UMAP with the labeled version in Fig. 2c allows identification of these cell types. It can be seen, for example, that *NKG7* expression becomes more specific to NK cells and CD8+ T cells and T gamma delta cells (Fig. S9a), while *CST3* becomes much more specific to monocytes, progenitors, and pDCs.

Differential expression tests (here a simple Wilcoxon test in scanpy using counts from either the raw CellRanger data or `CellBender`, after normalizing and log-transforming) show marked increases in specificity of four monocyte genes (*LYZ, CST3, S100A8, S100A9*) to the cluster “0: Monocytes C” as compared with cluster “4: B Naive”. The log2-fold changes typically increase substantially (by a factor of two), showing that even in a dataset that is relatively clean to begin with, `CellBender` significantly sharpens the differential expression signal. The gene *PTPRC* is also included as a “control” gene, since the expression of *PTPRC* is ubiquitous in immune cells, and it should not become more specific to one cell type or another after `CellBender`. We see that `CellBender` appropriately changes this log fold change estimate very little.

The data plotted in Fig. 2e quantifies the probability that each cell type is responsible for the expression of genes which have been removed by `CellBender`, broken down into bins of “how much” of each gene was removed. For example, the highly-removed genes (> 22%) are preferentially coming from basophils. A breakdown of this data on an individual-gene level is shown in Fig. S 6. Panel (a) shows the top ten genes by removed fraction, and the HPA reference shows that those genes are likely, more often than not, to come from basophils and neutrophils. As a point of comparison, panel (b) shows genes which `CellBender` has largely left alone, and they are much more randomly distributed across cell types in the reference. Additionally, some genes of interest, including some major PBMC marker genes, are shown in Fig. S 7.

**Table S 2:** Differential expression effect size (log2 fold change from scanpy Wilcoxon test) between “Monocytes C” (cluster 0) and “B Naive” (cluster 4).

|  | Raw data | After CellBender |
| --- | --- | --- |
| *LYZ* | 6.07 | **13.56** |
| *CST3* | 5.44 | **10.03** |
| *S100A8* | 6.06 | **12.95** |
| *S100A9* | 5.87 | **11.60** |
| *PTPRC* | 0.53 | 0.68 |

#### S.6.4 heart600k additional information

The `heart600k` dataset was published with `CellBender` v0.2.0 results available as part of the dataset on the Single Cell Portal (see S.4). We did not re-run `CellBender`’s v0.3.0 version on this dataset, but instead used the published results, which are the data shown in Fig. 2i-q. The main difference is using posterior regularization noise estimation strategy in `CellBender` v0.2.0 compared to to MCKP-based noise estimator in `CellBender` v0.3.0 which is expected to leave the results unchanged for the purposes of our discussion.

Similar to the previous PBMC analysis, Fig. S 10 shows that top-removed genes by `CellBender` can be attributed mainly to cardiomyocytes, and to some extent epicardial cells (this time, based not on an external reference, but on the dataset itself). This is quite consistent with observations of high amounts of background noise in cardiomyocyte marker genes including *TTN* and *RYR2*. Several marker genes of interest, including these two, are shown in Fig. S 11.

#### S.6.5 Remark on cell QC and mitochondrial reads

The intended use of `CellBender remove-background` is as part of a data processing pipeline, where `CellBender` comes after creation of a count matrix and before cell QC. Importantly, using all of the `CellBender` “non-empty” droplets downstream may not be appropriate, since not all “non-empty” droplets are high quality. We note that mitochondrial reads are often used as a cell QC metric downstream. In Fig. S12, we show that the low-UMI-count cells in the “arms” of the log-log plot of cross-species counts actually have very high mitochondrial read fraction compared to the high-quality cells.

#### S.6.6 Cells called on the single-nuclei rat6k dataset

**Table S 3:** Cells called by various methods for the `rat6k` dataset, before and after cell QC.

| Method | Before cell QC | After cell QC | Fraction surviving QC |
| --- | --- | --- | --- |
| CellRanger v2 | 5852 | 3805 | 0.650 |
| CellRanger v3 | 5869 | 3823 | 0.651 |
| EmptyDrops | 3109 | 1161 | 0.373 |
| CellBender | 10351 | 5878 | 0.566 |
| Exclusive to CellBender | 3922 | 1645 | 0.419 |

We run `CellBender remove-background`, `CellRanger v2.1.1`, `CellRanger v3.0.2`, and `EmptyDrops` (as in `Bioconductor v3.9`) with the default arguments other than (retain = 1000, lower = 80, ignore = 10). We removed all droplets outside of the union of cell calls made by the four methods. Next, we ran a standard `scanpy` analysis to cluster the cells and to find marker genes. Cells with mitochondrial read fraction greater than the 90th percentile have been removed, as well as cells with fewer than 100 unique genes expressed at nonzero levels. Cells with UMI count or gene count higher than the 85th percentile are also eliminated. These filters are employed to remove outliers and doublets from the analysis.

**Table S 4:** Cells called by various methods for the `rat6k` single-nuclei RNA-seq dataset, after cell QC, and clustered at Leiden resolution 0.5.

| Method | Detected Cells (per cluster, after cell QC) |  |  |  |  |  |  |  |  |  | Total |
| --- | --- | --- | --- | --- | --- | --- | --- | --- | --- | --- | --- |
|  | 0 | 1 | 2 | 3 | 4 | 5 | 6 | 7 | 8 | 9 |  |
| Union | 1842 | 1658 | 688 | 631 | 362 | 302 | 190 | 155 | 28 | 22 | 5878 |
| CellRanger 2.1.1 | 1226 | 1221 | 559 | 237 | 213 | 167 | 92 | 57 | 19 | 14 | 3805 |
| CellRanger 3.0.2 | 1234 | 1228 | 560 | 239 | 214 | 166 | 92 | 57 | 19 | 14 | 3823 |
| EmptyDrops | 36 | 198 | 49 | 232 | 161 | 287 | 21 | 149 | 18 | 10 | 1161 |
| CellBender | 1842 | 1658 | 688 | 631 | 362 | 302 | 190 | 155 | 28 | 22 | 5878 |

We notice after cell QC that (1) `CellBender` calls more cells than CellRanger or `EmptyDrops`, and (2) `CellBender` does not miss any of the cells called by the other methods. We argue below that (1) the extra calls made by `CellBender` are valid, and (2) excluding these cells implies discarding a *significant*, and *biased*, slice of the dataset. Cells called by each method, after cell QC, are shown in Table S4.

On the one hand, we notice that the extra cell calls made by `CellBender` are distributed essentially uniformly across the ten clusters. Crucially, the extra cell calls do not form a cluster of their own: had these extra cells actually been empty droplets, we would expect their expression profile to regress toward the background profile and cluster together. Even for clusters enriched with extra calls by `CellBender` (e.g. cluster 6), we find very specific marker genes (Fig. 3d shows that cluster 6 cells are pericytes). Again, this would not be expected if `CellBender` were erroneously identifying empty droplets (which would not be marked by unique marker genes that are absent from the other cell clusters).

We notice that the other methods, in particular `EmptyDrops`, fail to call a large fraction of cells in the most populous clusters. For instance, `EmptyDrops` has detected only 45 cells from cluster 3 (after quality-controlling cells as described above), compared to 680 cells called by `CellBender remove-background` (see Table S 4). This cluster, which can be unambiguously identified as cardiomyocytes, is populous while also producing disproportionately more transcripts per nucleus. This implies that the ambient background profile is likely to closely resemble that of cardiomyocytes.

`EmptyDrops` (run with lower UMI threshold set to 100, surely-cell retain threshold set to 1000, and Bonferroni-corrected FDR < 1%) calls many low-UMI-count cells that CellRanger v2 and v3 miss, though it also misses a large number of relatively high-UMI-count droplets along the rank-ordered UMI plot. We notice that the most populous clusters are also the most enriched in cell calls missed by `EmptyDrops` (Clusters 0, 1, 3; see Table S4). We believe that the issue originates from the frequentist approach used in `EmptyDrops`. Since the background profile indeed resembles the gene expression profile of the most abundant and transcript-rich cell types, the expression profiles of these cells are *accidentally* compatible with the background. Therefore, the Dirichlet-Multinomial p-values obtained on a single-droplet basis may not reach statistical significance for droplets that contain one of the major cell types, in particular, if the background pseudo-count scale *α* (a model parameter in `EmptyDrops`) is determined to be too large. By default, `EmptyDrops` determines *α* via a maximum likelihood procedure. We found that overriding *α* manually and using a smaller value generates more statistically significant cell calls, as expected.

`CellBender` does not suffer from this caveat since it effectively performs Bayesian model comparison using informative priors for both hypotheses (empty model *y_n_* = 0, non-empty model *y_n_* = 1; see Methods). The expression profile and the expected UMI count of the abundant cell types (and the background) are initially learned from the low- and high-count droplets, this prior information is used in comparing the two models, and the priors and posteriors are updated until a self-consistent solution is achieved.

Cell type labels in Fig. 3c were provided by author SF, as these cell types are closely related to ongoing projects in the Precision Cardiology Lab at the Broad Institute, and several manuscripts involving these cells are in preparation.

#### S.6.7 Denoising mixed-species hgmm12k

Additional quantification of removal of cross-species background noise in the `hgmm12k` dataset is shown here in several ways. Linear versions of the log-log plots from Fig. 4a-b are shown in Fig. S 13. The linear axis plots highlight the remaining linear trend in the DecontX data in panel (b), where cross-species counts increase in proportion to the total UMI count in a droplet.

The quartiles of remaining cross-species “contaminating” counts can be found in Table S 5. The suffixes on the `CellBender` data refer to the specified nominal false positive rate (nFPR) when the tool was run. The data show that truly only a handful of cross-species counts remain, even at the default nFPR of 0.01. These same data are shown graphically in Fig. S 14. It can be seen that `CellBender` removes nearly twice as much background noise using default settings than DecontX using default settings.

**Table S 5:**
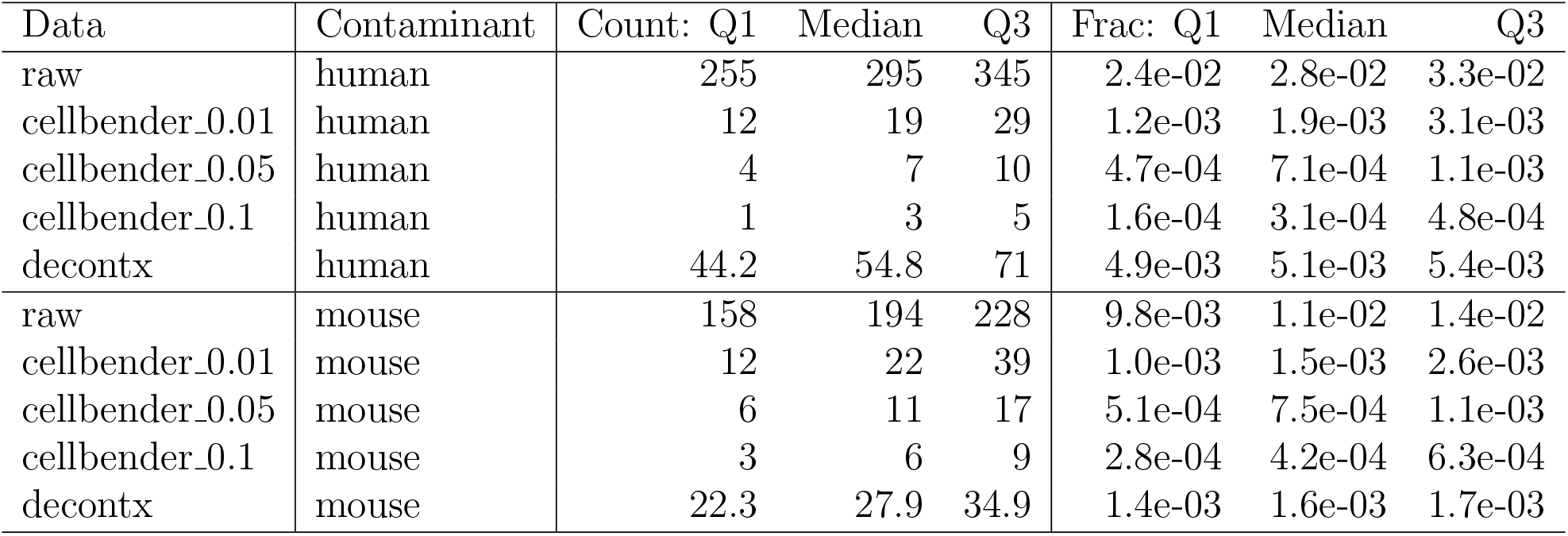
Removal of cross-species comtaminant counts from 10x Genomics human-mouse mixture dataset `hgmm12k` by `CellBender` and DecontX. Removal summarized as quartiles of fraction of cross-species counts remaining in cells (Frac), as well as raw counts (Count).

#### S.6.8 Concordance between nFPR and the empirical FPR

The nFPR parameter controls the total noise target in aggregate whereas the empirical FPR in simulated datasets is assessed at a more granular level, depending on one’s choice of micro- or macro-averaging, per cell or per gene. Nevertheless, we expect a good degree concordance between the two. This is shown for a particular simulated dataset in Table S6. The offset at nFPR = 0 is caused by the fact that the nominal false positive rate is a target for “extra” noise counts removed in addition to the level of noise expected to be present in the dataset, see Supplementary Section S.1.5.

#### S.6.9 CITE-seq pbmc5k additional information

Fig. S15 shows some additional information about CD45 isoform expression in the `pbmc5k` dataset. Panel (a) is a dotplot showing marker genes for several cell types common in PBMC datasets, along with which clusters these genes show up in. Panel (b) tentatively names the clusters and shows a UMAP. Panel (c) shows the expression of *PTPRC* (the gene for CD45), along with the CD45RO isoform and the gene *HNRNPLL*. CD45RO is expected to correlate with *HNRNPLL* expression, since *HNRNPLL* is a related splicing factor, which we see it does. This trend is less visually clear in a UMAP of the raw data. The major cell types expressing CD45RO appear to be monocytes C/NC/I, along with T CD8+ EM / TE / gamma delta, and T reg / helper.

**Table S 6:** Target nominal false positive rate (nFPR) parameter versus the empirical FPR of the `CellBender` output for the simulated data shown in Fig. 4c-g. Note that the results are dataset-dependent.

| Target nFPR | Empirical FPR |
| --- | --- |
| 0 | 0.026 |
| 0.01 | 0.033 |
| 0.02 | 0.040 |
| 0.03 | 0.049 |
| 0.04 | 0.057 |
| 0.05 | 0.067 |
| 0.06 | 0.076 |
| 0.07 | 0.086 |
| 0.1 | 0.116 |
| 0.2 | 0.217 |
| 0.5 | 0.456 |
| 0.6 | 0.516 |
| 0.7 | 0.569 |
| 0.9 | 0.648 |

Fig. S16 shows the concordance of RNA counts and antibody counts before and after `CellBender` for all the “Antibody Capture” features in the `pbmc5k` dataset. Panel (a) is the raw data, showing that correlations do exist, but that for every antibody, there are significant counts even when the RNA counts go to zero. Panel (b) shows the same data after `CellBender`, and we see that the antibody counts are more often near zero in clusters where the RNA counts are zero. This leads to an increase in the Pearson correlation between the fraction of cells per cluster with nonzero RNA and antibody counts, as shown in panel (c), which leads to greatly sharpened biological inferences about cell-type specificity of expression.

**Figure S 4:**
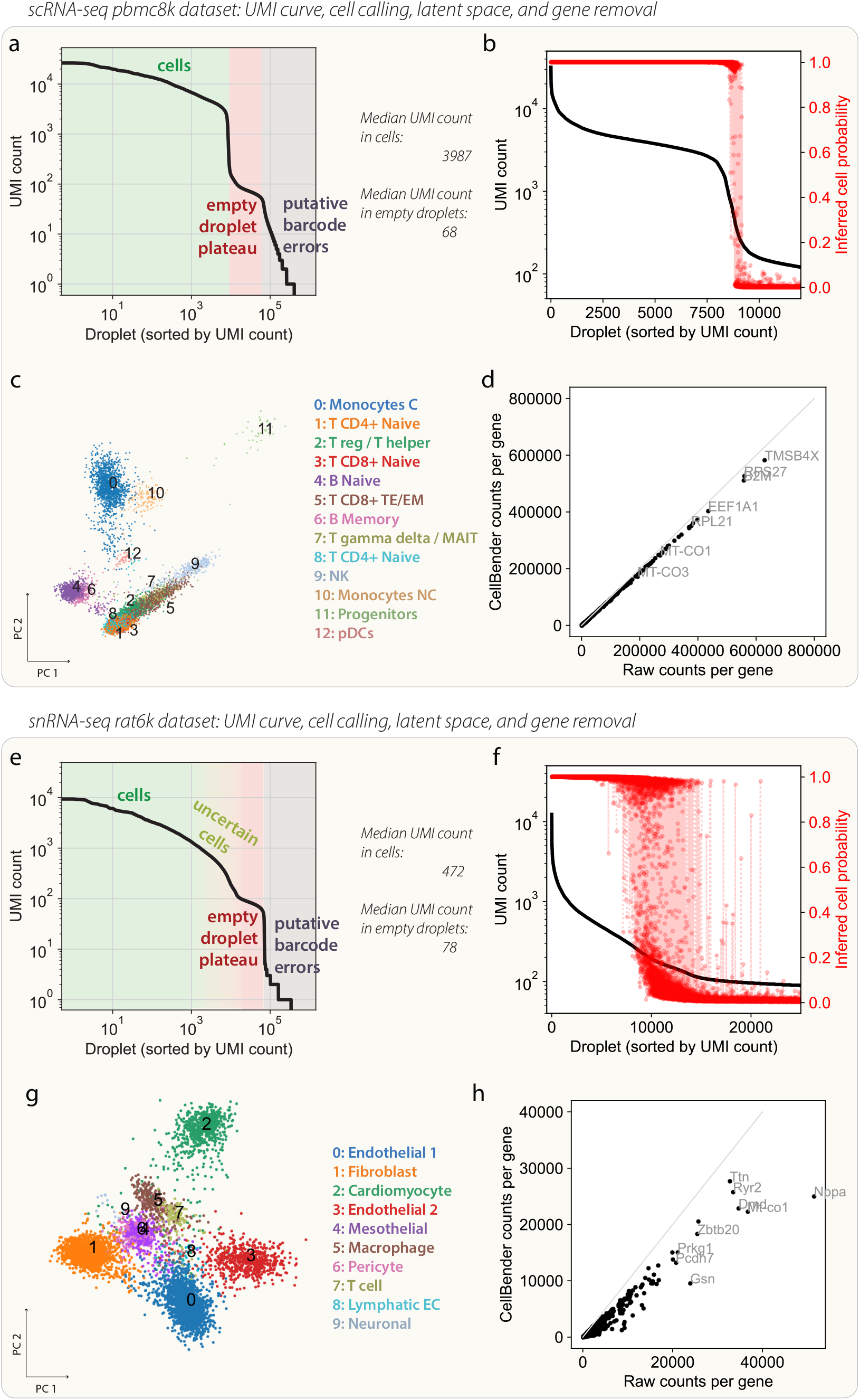
UMI curves from the raw data together with various `CellBender` outputs for the (a-d) `pbmc8k` and (e-h) `rat6k` datasets. (a,e) The raw UMI curves, annotated with areas of cells and empty droplets. Notably, the distinction is much more difficult in (e), the nuclei dataset extracted from heart tissue. (b,f) Cells probabilities inferred by `CellBender` on same UMI curves from (a,e) respectively. The region of transition from “surely-cell” to “surely-empty” is much broader in the snRNA-seq dataset. (c,g) First two principal components of the latent gene expression embedding inferred by `CellBender`, colored by Leiden clustering from a separate scanpy analysis. The structure very closely reflects the labels attributed by that separate analysis. (d,h) Scatter plots showing removal of each gene by `CellBender` (each dot is a gene, *MALAT1* is off-scale). Several top denoised genes are indicated.

**Figure S 5:**
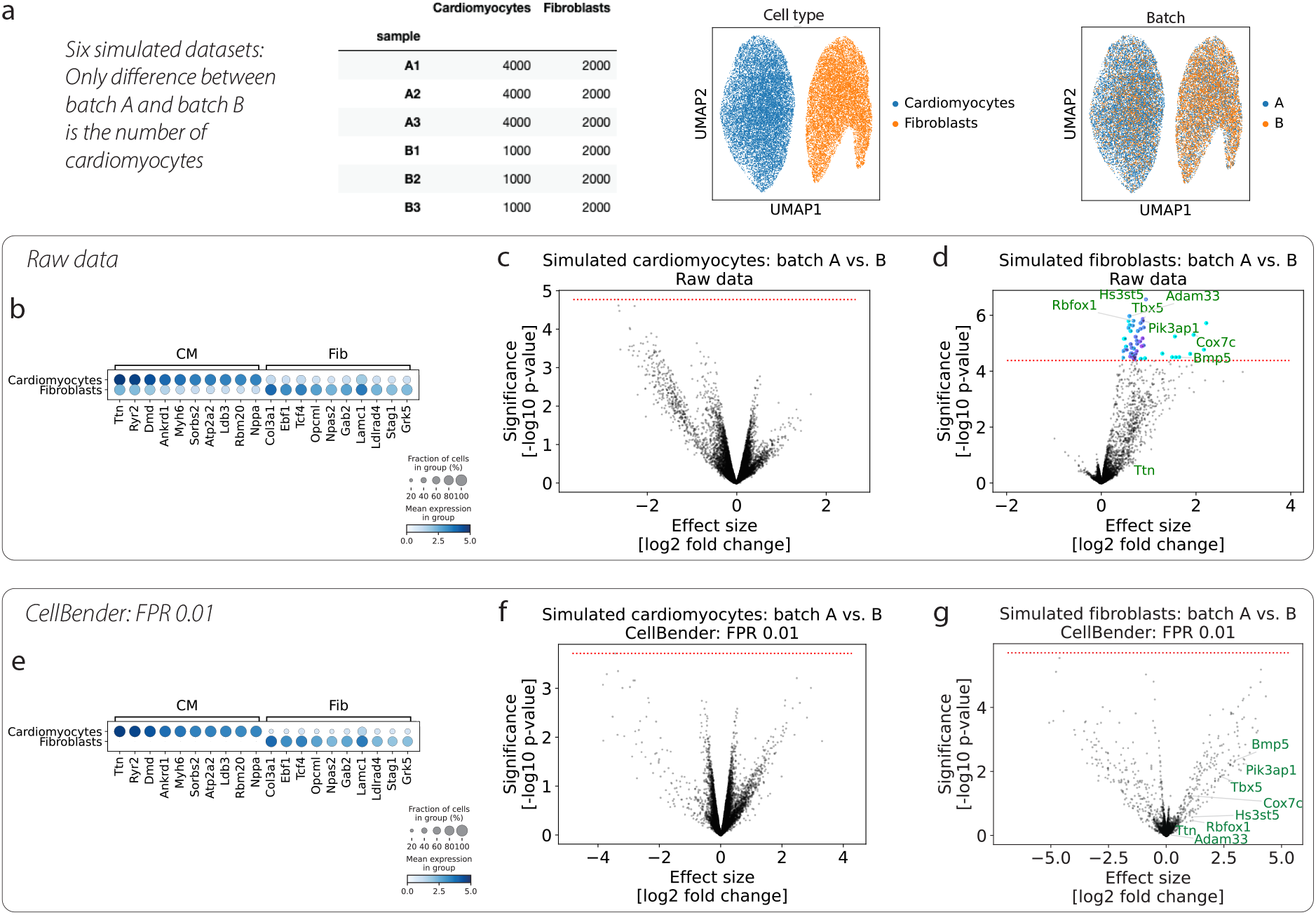
Systematic background noise as a source of batch variation and spurious differential expression across batches. (a) Setup of the cohort of simulated datasets, where there are two cell types whose expression profiles are taken from real data (`rat6k`) for cardiomyocytes and fibroblasts. The only difference between simulations from batch A and batch B is the number of cardiomyocytes. Noise ends up being different in the two batches due to these cell number differences. The “truth” in this simulated cohort is that there are no differences between a cell type’s expression profile between batches. (b-d) Raw data. (e-g) `CellBender` denoised data. (b) Dotplot showing top cardiomyocyte and fibroblast marker genes. Background noise causes marker genes to show up in the off-target cell type at a low level. (e) Marked cleanup of the dataset at an aggregate level. (c,f) The cardiomyocytes show no differentially expressed genes between batch A and B, before or after `CellBender`. (d) In the raw data, many genes show up as being significantly differentially-expressed due to background noise. (g) After `CellBender`, these spurious results have disappeared (a few of which are labeled). Benjamini-Hochberg-corrected FDR value for significance (red dotted line) is 0.01 in all volcano plots.

**Figure S 6:**
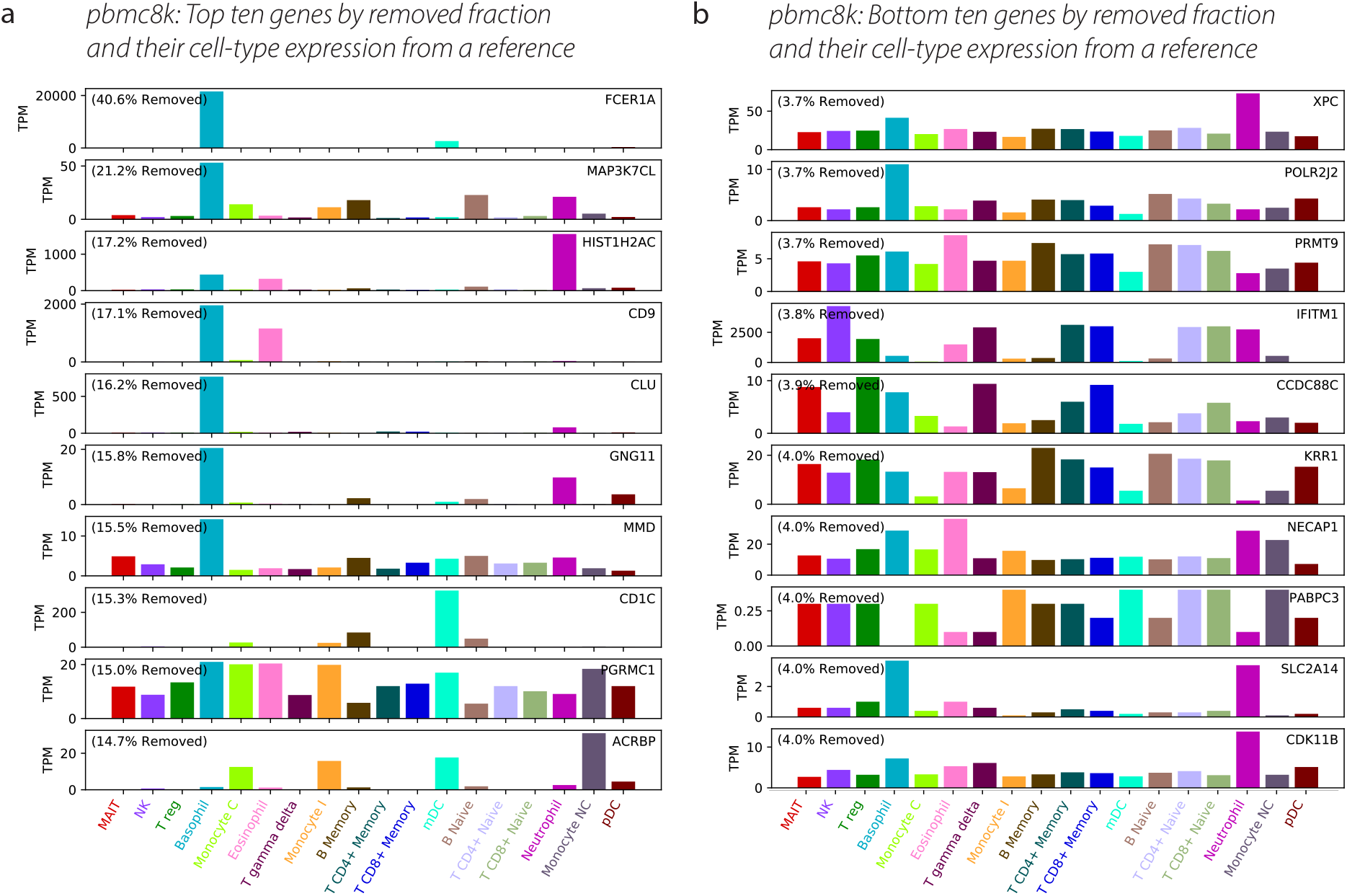
(a) The top ten genes ranked by fraction removed by `CellBender` (above an expression cutoff) from the `pbmc8k` dataset, indicating that many of these highly-removed genes have high expression in basophils and neutrophils. (b) For comparison, the bottom ten genes ranked by fraction removed (above an expression cutoff). These genes are much more random as far as cell-type specificity. The HPA immune reference [38] was used to attribute TPM expression for these genes to the immune cell types in the reference.

**Figure S 7:**
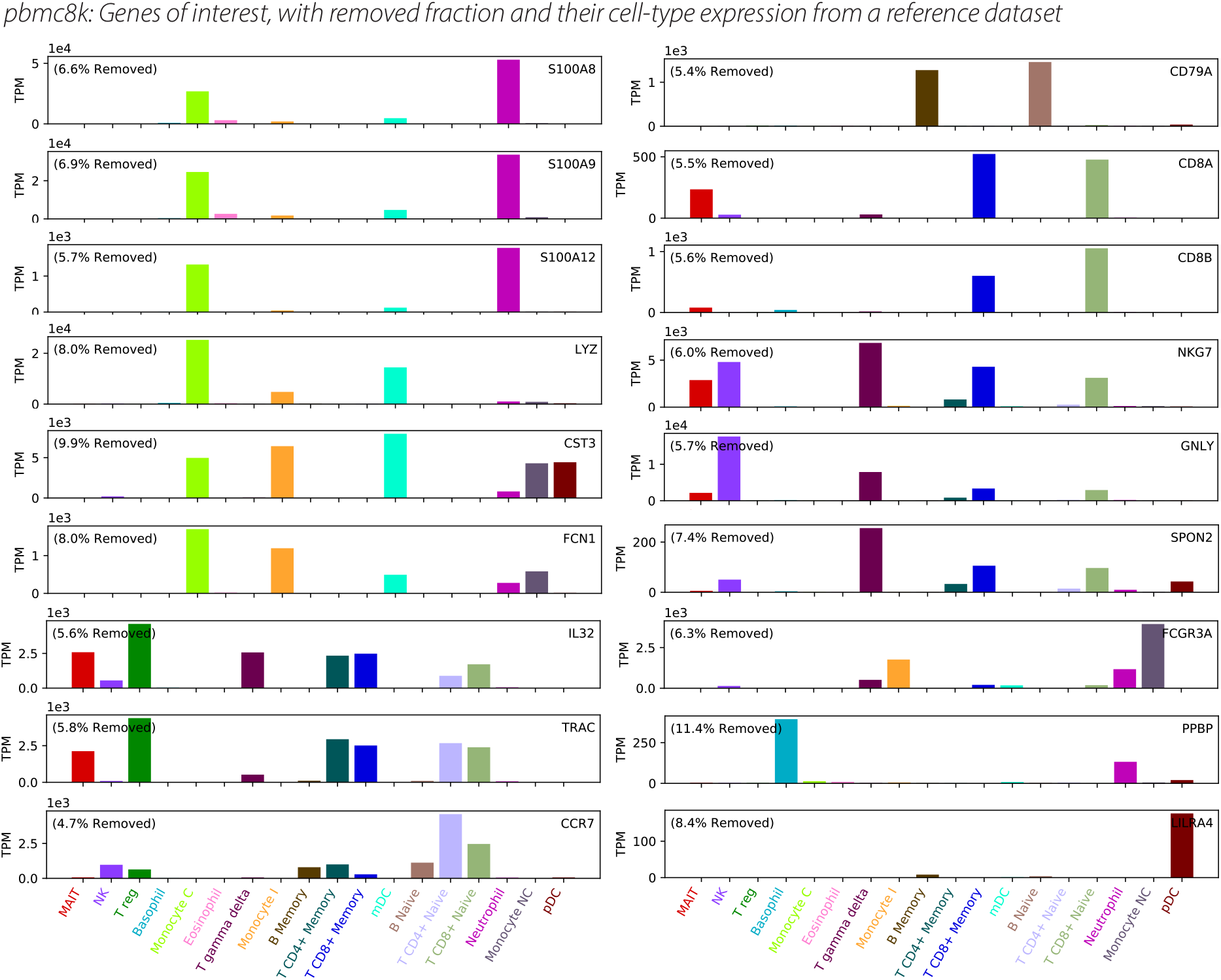
Selected genes of interest, showing fraction removed by `CellBender` from the `pbmc8k` dataset, along with expression per cell type. The HPA immune reference [38] was used to attribute TPM expression for these genes to the immune cell types in the reference.

**Figure S 8:**
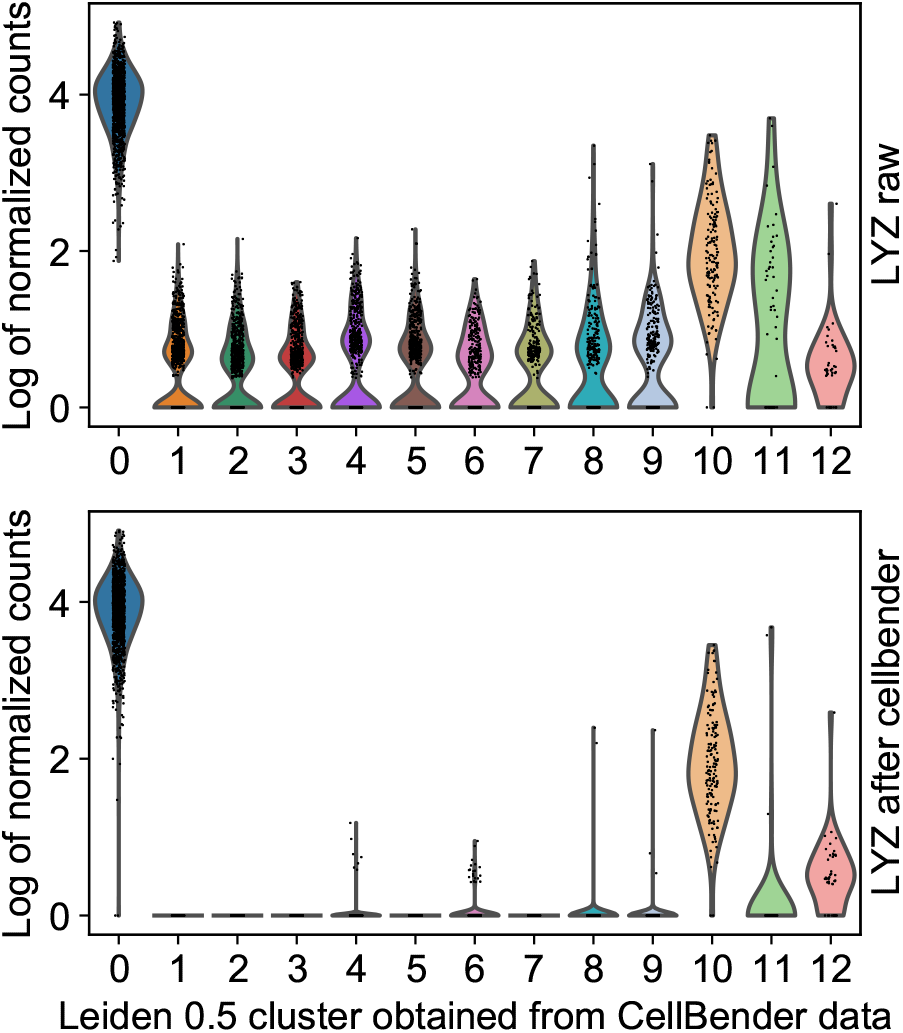
Violin plots showing the count distributions of lysozyme, *LYZ*, per cluster before and after `CellBender` denoising (nFPR 0.01). The off-target counts are effectively removed, with counts remaining in clusters 0 (CD14^+^ monocytes C), 10 (FCGR3A+ monocytes NC), and 12 (plasmacytoid dendritic cells).

**Figure S 9:**
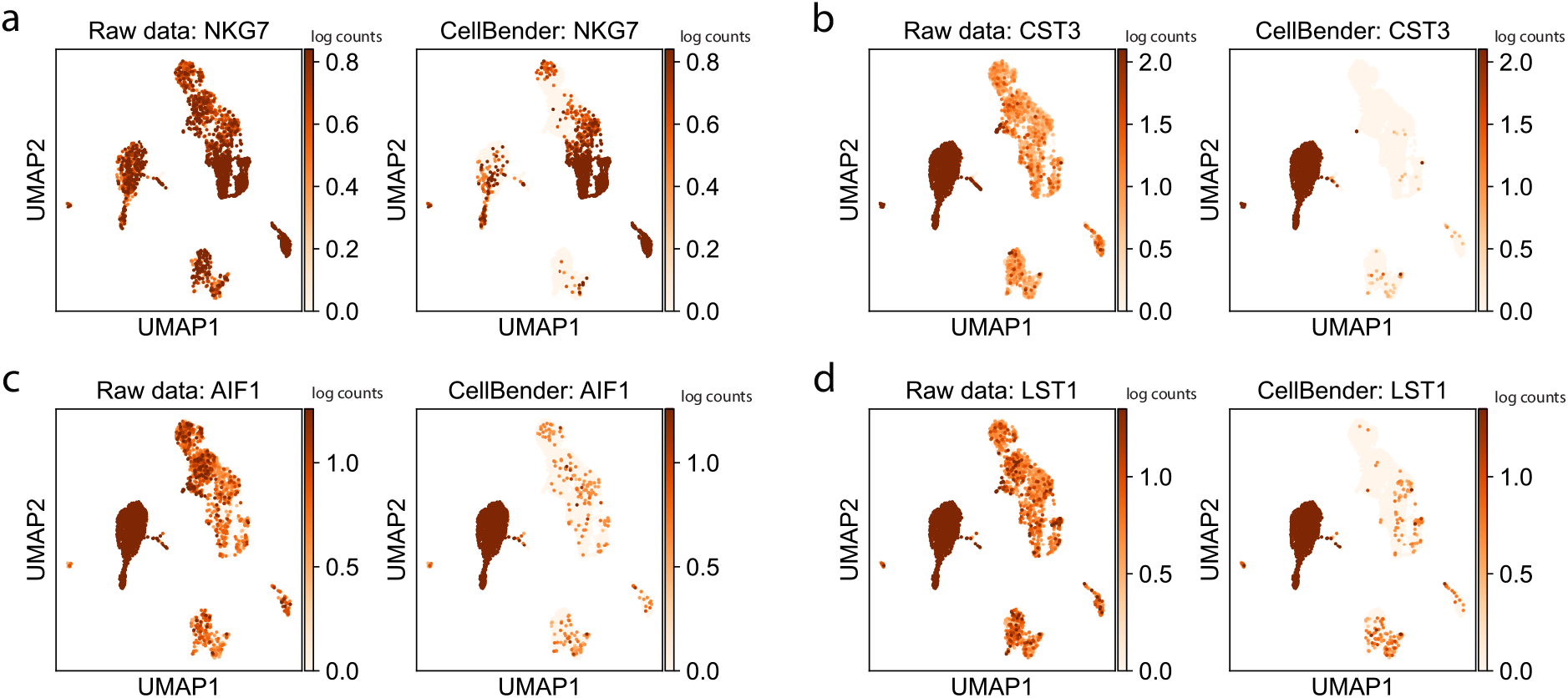
UMAPs created from the `CellBender`-analyzed `pbmc8k` data, showing increased expression specificity of marker genes for different cell types after `CellBender` denoising as compared to the raw data.

**Figure S 10:**
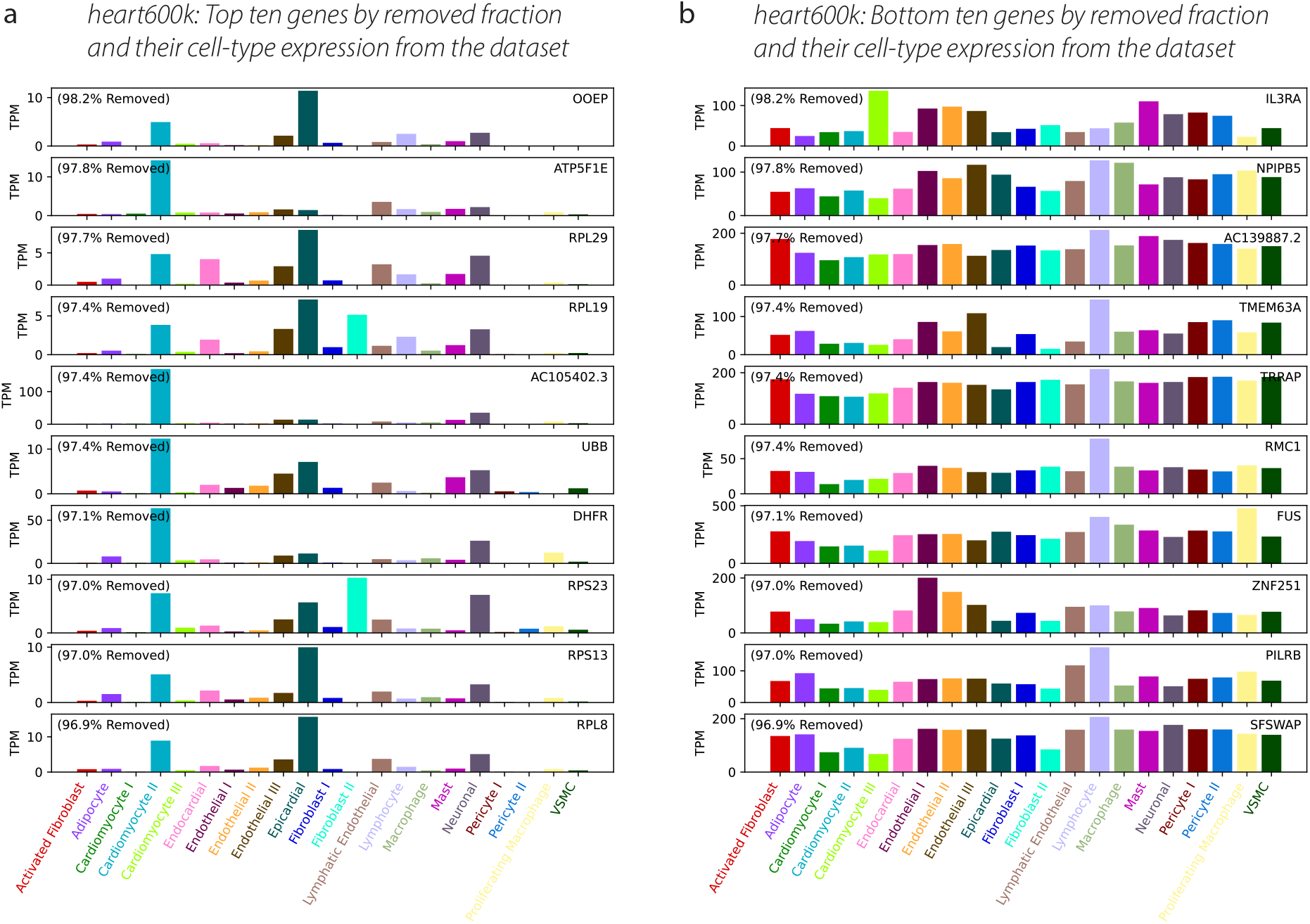
(a) The top ten genes ranked by fraction removed from `heart600k` by `CellBender` (above an expression cutoff) from the `heart600k` dataset, indicating that many of these highly-removed genes have high expression in cardiomyocytes (as well as epicardial cells). (b) For comparison, the bottom ten genes ranked by fraction removed (above an expression cutoff). These bottom genes are fairly random in terms of cell-type specificity. The `heart600k` dataset was used to attribute TPM expression (computed as normalized gene expression per cell type times 1e6) for these genes to each cell type.

**Figure S 11:**
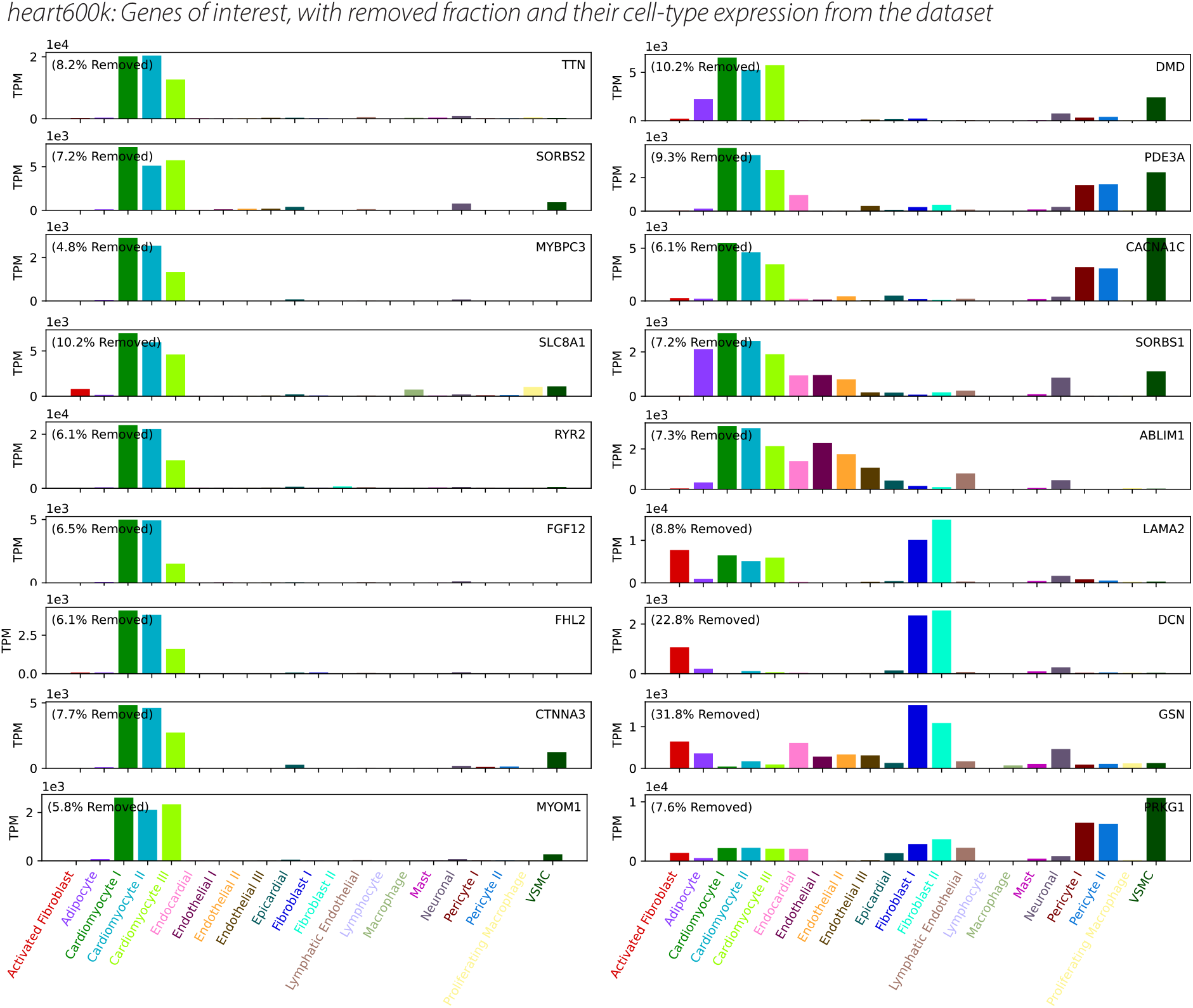
Selected genes of interest, showing fraction removed by `CellBender` from the `heart600k` dataset, along with expression per cell type. The `heart600k` dataset was used to attribute TPM expression (computed as normalized gene expression per cell type times 1e6) for these genes to each cell type.

**Figure S 12:**
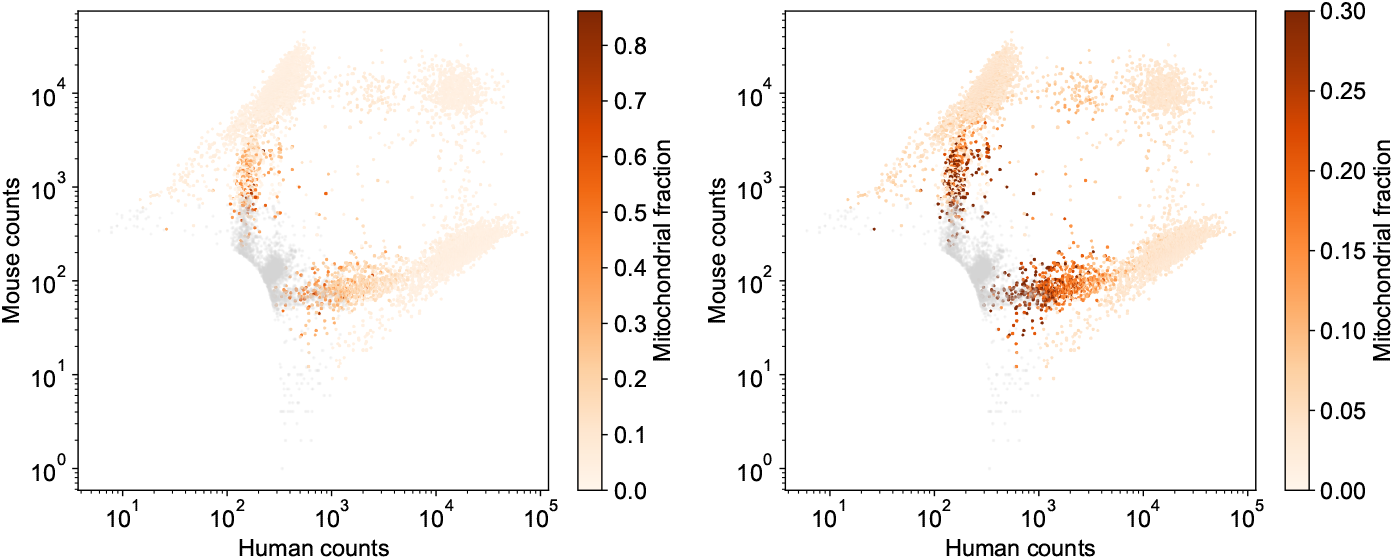
Fraction of reads per droplet that align to mitochondrial genes in the `hgmm12k` dataset. Gray dots are determined to be empty droplets by `remove-background`. (Right) Colorbar axis is truncated to a maximum of 0.3 to enhance contrast. This highlights the fact that, while the high-mitochondrial-fraction droplets are clearly not empty, they may not be desirable for downstream analysis either. Further cell QC is needed above and beyond `remove-background`’s elimination of empty droplets.

**Figure S 13:**
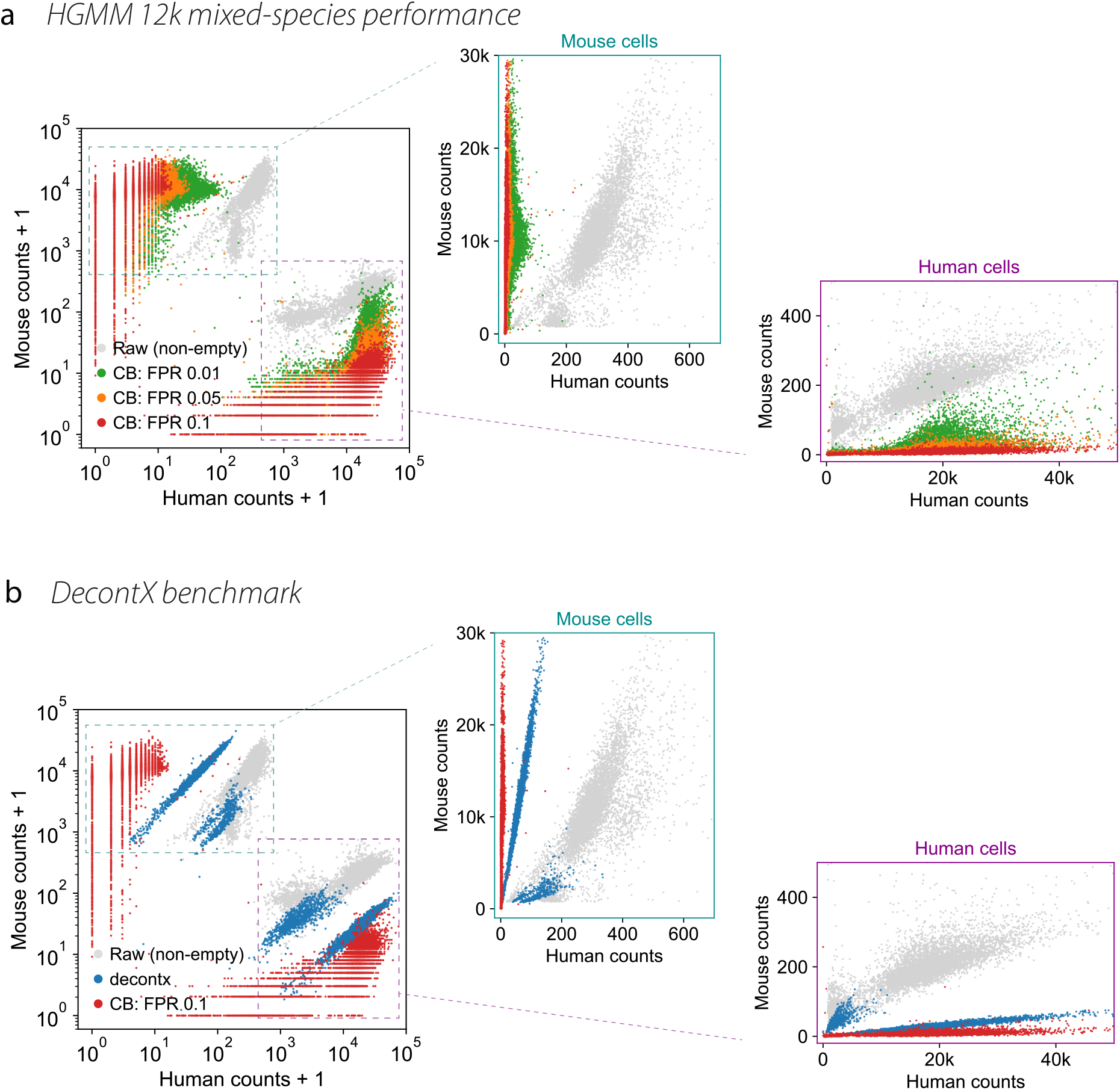
`hgmm12k` dataset, showing same results as in Fig. 4a-b, along with data plotted on linear axes. (a) `CellBender` (CB) run at several nFPR values. (b) DecontX as a benchmark. A residual linear trend of remaining cross-species counts is visible (note the slope of blue points).

**Figure S 14:**
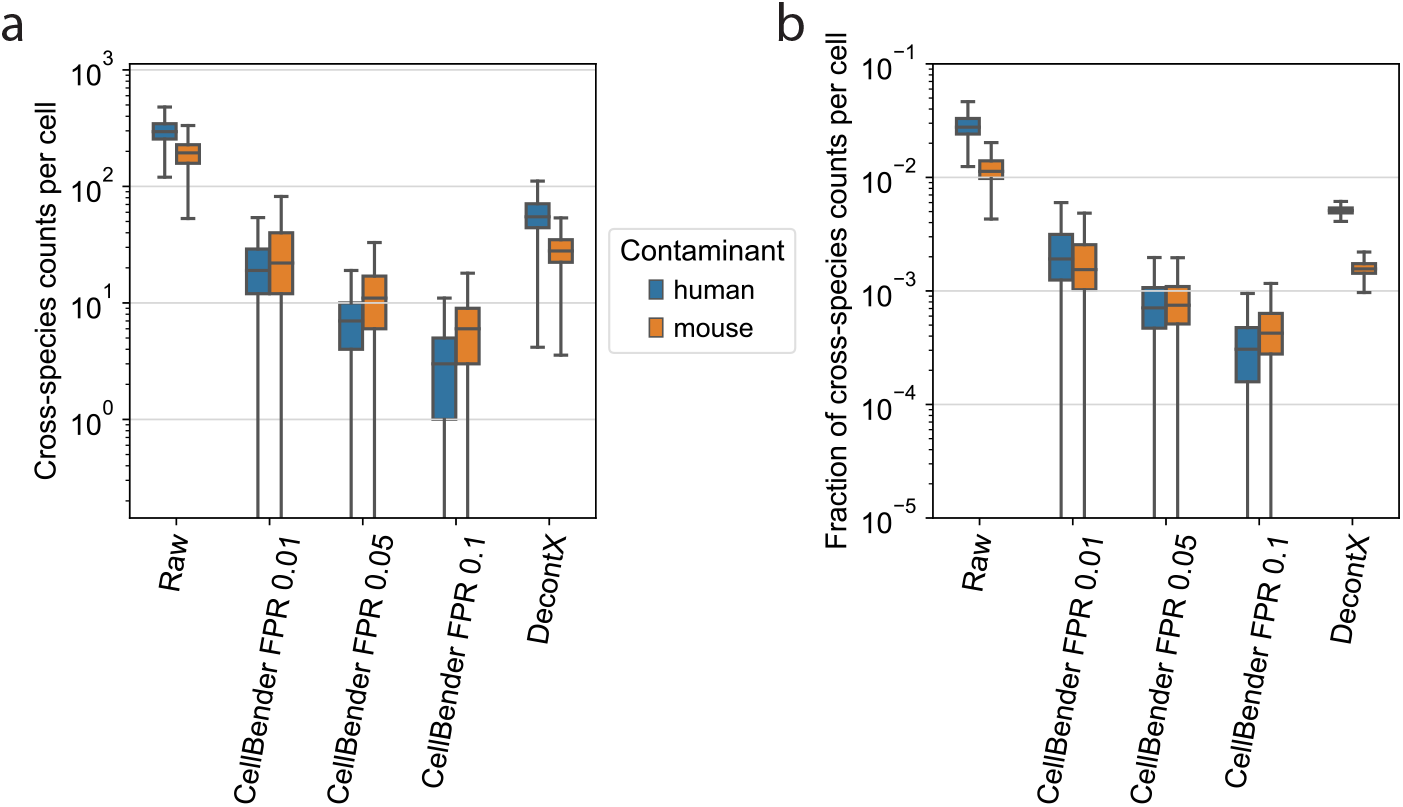
Quantification of removal of cross-species counts from `hgmm12k` dataset (data are the same data as Fig. S 13) (a) The number of contaminant counts in cells from the other species. Boxes represent the first to third quartile, with the median drawn as a line. The whiskers are 1.5 times the interquartile range. (b) Same data as panel (a), but as a fraction of the total cell counts.

**Figure S 15:**
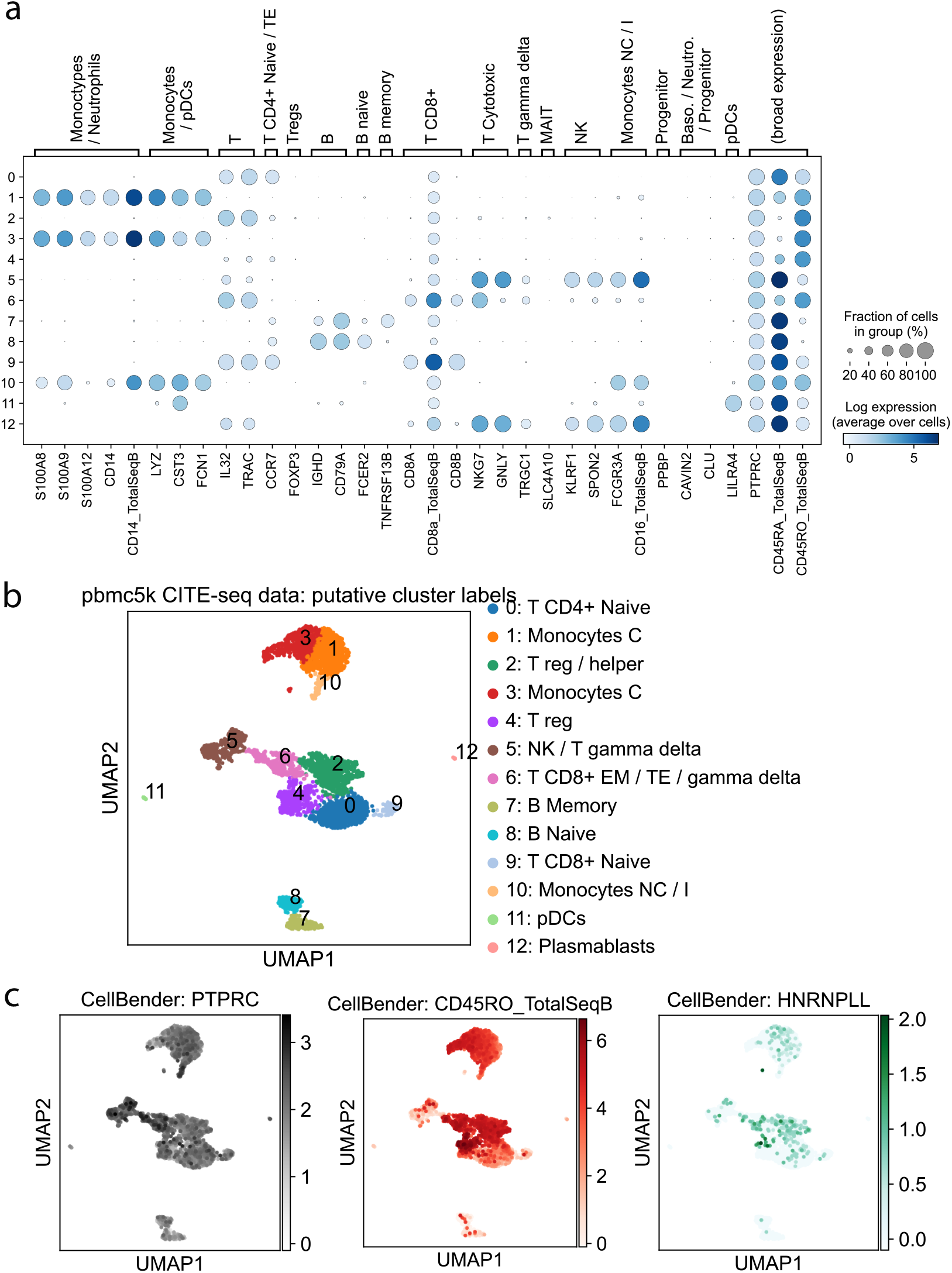
`pbmc5k` CITE-seq dataset. (a) Marker genes and proteins for several immune cell types commonly observed in PBMC datasets, showing in which Leiden cluster they appear (post-`CellBender` data shown). (b) UMAP of the `pbmc5k` data, colored by Leiden cluster, and labeled based on the dotplot in (a) along with other marker genes. (c) Distribution of the gene *PTPRC*, one measured isoform CD45RO (after `CellBender`), and the distribution of *HNRNPLL*, a positive splicing factor for CD45RO. `CellBender` increases the specificity of CD45RO expression (raw data can be found in Fig. 5c for comparison).

**Figure S 16:**
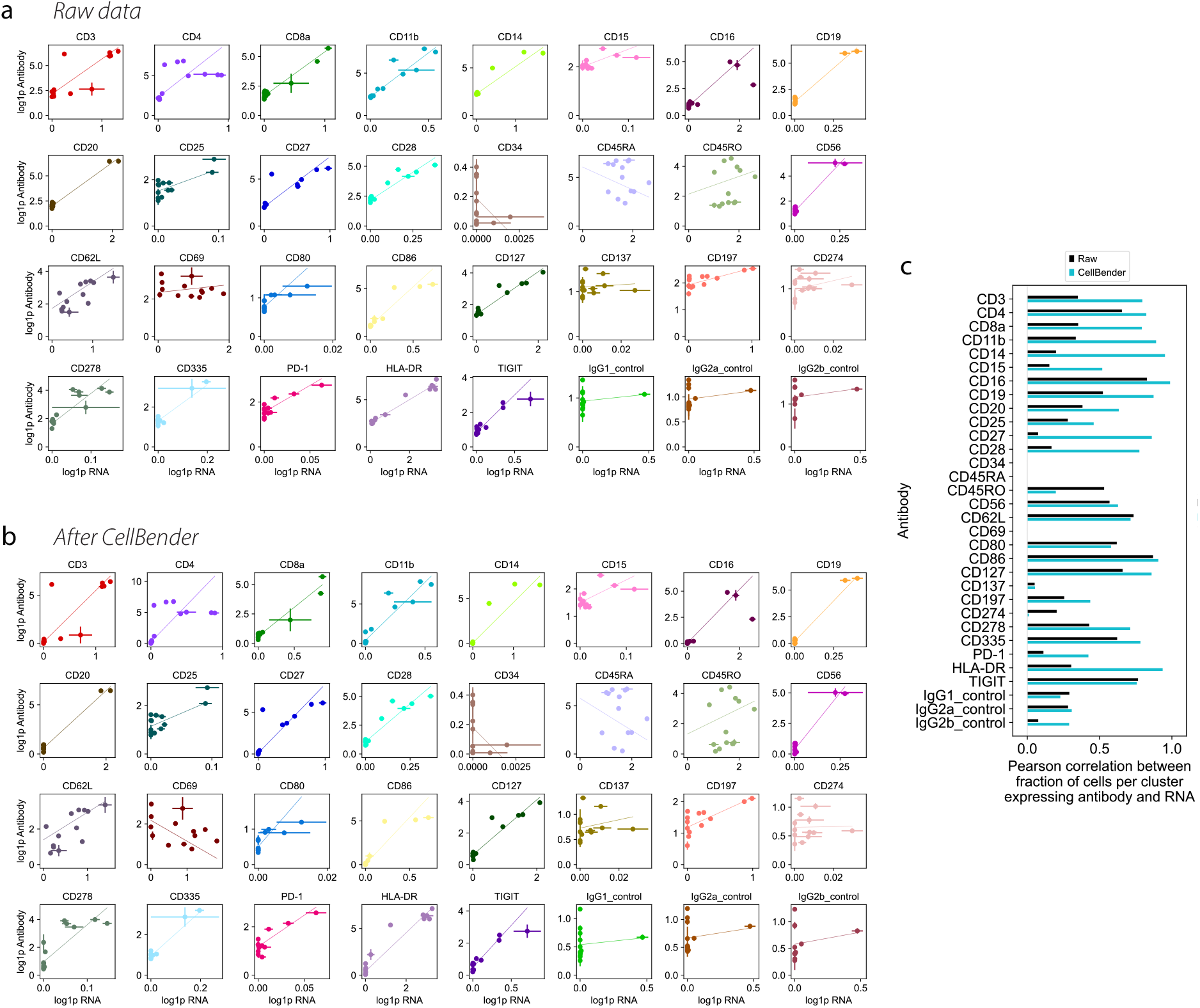
(a) Scatter plots of each measured antibody in the `pbmc5k` CITE-seq dataset. Raw data from CellRanger, where dots are mean counts in each of the 14 clusters, along with linear fits using weighted ordinary least squares. Numeric values on axes are log1p of unnormalized counts. Most antibodies exhibit a reasonably strong correlation between RNA counts and antibody counts. Notable exceptions include CD45RA and CD45RO, which are different and typically mutually exclusively expression isoforms of the same gene, *PTPRC*, and CD34, CD69, CD137, and CD274, where antibody counts and RNA counts do not seem well-correlated in the raw data. (b) Same as (a), but using the output data from `CellBender`. Most fits have intercepts very near zero, and the linear relationships are preserved. (c) Pearson correlation between fraction of cells with nonzero RNA and fraction of cells with nonzero counts of the corresponding antibody, for raw data and after `CellBender`. The increased correlation leads to much greater cluster specificity. Bars with Pearson correlation < 0 are omitted.

